# Mesoscale landscaping of the TRIM protein family reveals a novel human condensatopathy

**DOI:** 10.1101/2025.01.02.630836

**Authors:** Hari R. Singh, Vineeta Sharma, Jik Nijssen, Andrei Pozniakovski, Alexander Rubin, Lorrin Liang, David Ball, Sunwoo Hong, Victoria Gauntner, Guanghao Yu, Arathi Ranga, David Salant, Friedhelm Hildebrandt, Anthony A. Hyman, Amar J. Majmundar

**Author notes:** Equal contribution first authors (Hari R. Singh, Vineeta Sharma). Co-corresponding authors **Correspondence should be addressed to**, Amar J. Majmundar M.D. Ph.D. Division of Nephrologym, Boston Children’s Hospital 300 Longwood Avenue, Boston, Massachusetts 02115; Anthony A Hyman, Max Planck Institute of Molecular Cell Biology and Genetics Pfotenhauerstrasse 108, Dresden 01309.

## Abstract

The mesoscale organization of cells is central to cellular physiology and pathology. Cellular condensates often form via biomolecular phase separation, mediated by intrinsically disordered regions (IDRs) and represent a key mechanism for mesoscale organization. The TRI-partite Motif (TRIM) family of ubiquitin ligases is implicated in diverse cellular functions and disease, yet the role of biomolecular condensation in TRIM family organization remains understudied. Here, we systematically investigate the mesoscale localization of 72 TRIM proteins, revealing that a majority form condensates in distinct cellular compartments. IDR content correlates with dynamic condensate formation, suggesting a critical role in mesoscale organization. Focusing on TRIM8, associated with a neuro-renal disorder, we demonstrate that disease-causing truncations of the TRIM8 C-terminal IDR result in a *condensatopathy*, characterized by disrupted condensation, proteasomal regulation, and TAK1/NFκB signaling. Functional assays in cellular and animal models link these disruptions to podocyte dysfunction and impaired response to injury. Our findings establish a framework for understanding *condensatopathies* and the mesoscale principles governing TRIM family organization and function.

## INTRODUCTION

Mesoscale organization of the cell dictates its physiology which can go awry in disease. Membraneless biomolecular condensates provide one mechanism for this organization by enriching biomolecules within and by interfacing them with external complexes and systems^1,2^. Condensates play critical roles in normal cellular functions^3^, in pathologic states^4–6^, and even in cellular partitioning of therapeutics^7^. These structures are formed through weak multivalent interactions. Frequently, this occurs through macromolecular phase separation mediated by intrinsically disordered regions (IDRs)^1,2^. However, the prevalence of condensates as a general mechanism for mesoscale organization (e.g. within a class of proteins) is not well defined. It also remains unclear what precise physiochemical properties/cellular context within functionally defined IDRs and their surrounding domain architecture would govern biomolecular condensation.

The <u>TRI</u>-partite <u>M</u>otif (TRIM) family of ubiquitin ligases represents the largest subclass of human ubiquitin ligases^8–12^ (**Figure 1A, Suppl Figure S1, Suppl Table ST1, Suppl Files SF1-SF78**). The family is characterized by a conserved multi-valent N-terminal portion comprised of (i) a **R**ING domain mediating ubiquitination of substrates; (ii) one or two **B**-**B**ox domains, and (iii) frequently a **c**oiled-coil domain required for homo/hetero-dimerization. In contrast to this classical **RBBC** structure, the C-terminal portions of TRIM family proteins are far more variable with a number of ligase-specific ordered domains that mediate interaction with proteins (e.g. SUMO1)^13^ and supra-molecular polymers such as chromatin (e.g. PHD-BROMO domain of TRIM33), RNA (e.g. NHL repeats of TRIM2 or 3, PRY-SPRY domain of TRIM25), and microtubules (e.g. COS domain of TRIM46) (**Figure 1B**). While some TRIM family proteins were anecdotally observed to form foci^14–16^, it remains unclear if biomolecular condensation is a frequent mechanism for mesoscale organization of TRIMs.

**Figure 1.**
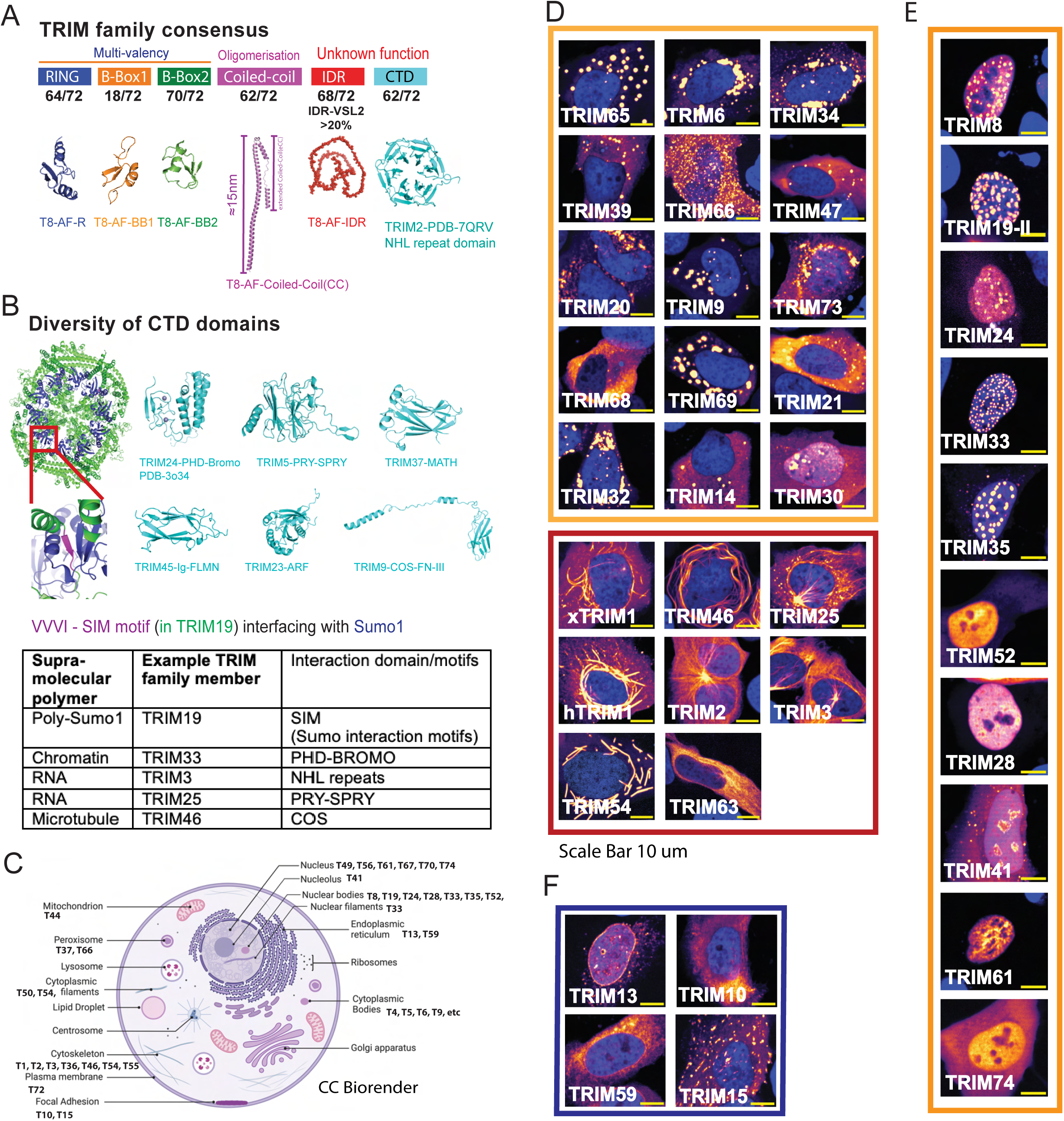
Overview of the TRIM family proteins that display a variety of specific mesoscale organizations in cells. **A.** The TRIM family consensus domain architecture is shown with N-terminal multivalent RBBC domains that consists of RING, Type I B-Box, Type 2 B-Box, and Coiled-coil domains. This is followed by generally short/long IDR. (Note that IDRs are also present as spacers between RBBC domains and at the N-terminus, generally short i.e. <30 a.a.) The IDR is then followed by a C-terminal structured domain, which frequently interacts with a supra-molecular polymer like (SMPL) surfaces. The relative frequencies of these structures within the TRIM family are shown (see **Suppl Table ST1** for domain structures based on primary sequence interpretation and evaluation of Alpha-fold generated structures in **Suppl Figure S1** and **Suppl Files SF2-SF78**.) **B.** A tetramer of TRIM19 modelled with a concatenated 10-mer (with GSG linker) of Sumo1 protein is shown after generation in Alpha-fold3. Inset shows the SIM (Sumo interaction motif) in purple interfacing the Sumo1 protein monomer (in blue). Additional structures are shown of exemplary C-terminal domains within the TRIM family proteins and summarized below. **C.** A visual summary of the consensus global cellular mesoscale localization patterns that were observed in HELA and/or U2OS cells (**see Suppl Figure S2**) for the TRIM family. **D.** Representative examples are shown of the TRIM family proteins that localize to the cytoplasm, including cytoplasmic filaments and the cytoplasmic bodies, in HELA cells. (For U2OS cells, **see Suppl Figure S2.**) **E.** Representative examples of the TRIM family proteins that predominantly localize to the nucleus in bodies or diffuse patterns in HELA cells are shown. (For U2OS cells, **see Suppl Figure S2.** While localization patterns were generally concordant, note that TRIM52 in HELA cells show diffused localization, while in U2OS cells it shows nuclear bodies.). See further analysis of these proteins in Figure 2E. **F.** Representative examples are shown of the TRIM family proteins that localize to the membranous compartments.

TRIM family proteins regulate a myriad of cellular functions including protein quality control through proteasomal degradation and autophagy, signal transduction, nuclear organization and transcriptional regulation^11,12,17–19^. These functions mediate critical tissue and organism scale roles in humans including innate immunity and host-pathogen interactions, cardiac function and disease, and carcinogenesis^11,12,17–20^. This is well evidenced by the array of Mendelian disorders caused by germline variants in TRIM family genes (11/72 TRIM in 9 OMIM genetic phenotypes, 4 provisional OMIM genetic phenotypes, and 1 additional sourced condition) (**Suppl Table ST2**).

Previously, we and others had discovered that dominant variants in *TRIM8* cause a novel neuro renal syndrome (OMIM 619428)^21–27^. We showed that germline variants in this severe pediatric onset disorder lead to aberrant mesoscale organization of TRIM8^23^. In this condition, the kidney disease manifests during childhood with proteinuria, renal scarring known as focal segmental glomerulosclerosis (FSGS), and progressive renal dysfunction. *TRIM8* is highly expressed in critical kidney filtering cells, podocytes^9,23^. Podocyte dysfunction is a hallmark of the majority of proteinuric kidney disease genes^28^, suggesting TRIM8 may regulate important functions in this cell type. Disease variants cause truncation of the TRIM8 C-terminus, and this results in loss of localization to nuclear bodies^23^. However, it remains unknown whether (i) TRIM8 nuclear foci are, in fact, biomolecular condensates and (ii) if this mesoscale organization is mediated by IDR-directed condensation that is disrupted in *TRIM8-*associated disease.

In this study, we sought to systematically investigate the mesoscale localization of the TRIM family of proteins within the framework of biomolecular condensation. Specifically, we aim to: (1) map the mesoscale organization of TRIM family proteins to assess their localization and behavior in cellular systems; (2) explore how TRIM family organization bridges molecular-scale features to mesoscale structures; (3) examine how disease-causing IDR deletions in TRIM8 underlie a *condensatopathy*, linking molecular disruption to human genetic disease; and (4) conduct a multi-scale mechanistic characterization of TRIM8-loss of condensation associated neuro-renal syndrome. This allows us to establish a framework for understanding condensatopathies and their connection to mesoscale organization.

## RESULTS

### TRIM family proteins display various sub-cellular localizations in cells

To evaluate the mesoscale organization of TRIM family members, 72 TRIM family proteins were either acquired or cloned into fluorescent tagged expression vectors (**Suppl Table ST1, Suppl File SF1**). The plasmids were transiently transfected into HELA (**Figure 1C-F**) and U2OS (**Suppl Figure S2**) cell lines and images acquired for at least 100 cells. We assessed and divided the mesoscale organization of TRIM family proteins into (i) specific patterns (bodies greater than 300 nm in diameter, filaments, membranous or multiple types) or diffuse only as well as (ii) sub-cellular localizations. Predominant mesoscale patterns were defined as being present in >50% in at least one cell type (**Suppl Table ST3**).

Most TRIM family proteins (50/72, 69%) exhibited specific mesoscale localization patterns in >50% of HELA and/or U2OS cells, while a minority exhibited a diffuse pattern only (22) (**Suppl Table ST3**). We observed that a large majority of the 72 proteins localize to some extent to nuclear or cytoplasmic bodies representing potential condensates (60 TRIMs, 41 with predominant localization to cellular bodies) (**Suppl Table ST3**). Several TRIM family proteins displayed predominant filament localization (7 members) or membranous patterns (6) in the cytoplasm (**Figure 1D, 1F, Suppl Figure S2B**). Across all localization patterns, most TRIMs (60) were in the cytoplasm while the remainder were observed in primary mesoscale patterns within the nucleus, pan-cellularly, or a combination. Of the 12 members exhibiting any nuclear localization in >50% of cells, eight TRIMs (TRIM8, −19, −24, −28, −33, −35, −41, −52) were robustly present in nuclear bodies (**Figure 1E, Suppl Figure S2A**). This screen revealed the diversity of mesoscale organization in the TRIM family and indicated that a high fraction of TRIM proteins localize to potential biomolecular condensates, consistent with prior examples^14–16^.

We next assessed the dynamic behavior of 48 TRIMs including 39 of the putative condensates forming TRIMs and 6 of the filaments forming TRIMs (**Figure 2A-D, Suppl Movies SM1-SM51, Suppl Figure S3-S11, Suppl Table ST3**) using live cell microscopy and conducting FRAP assays. Of the TRIM family proteins that localize to filaments, some were dynamic with a range of 50% recovery times (varying between seconds to minutes), while others were non-dynamic with minimal to no recovery over the measurement time (30 seconds to 5 minutes). TRIMs localizing to cellular bodies, similarly, were split between those displaying dynamic behavior in cells with 50% recovery times varying between seconds to minutes (20/39) and those showing non-dynamic behavior in this measurement period (19/39). In conclusion, TRIM proteins display the whole spectrum of cellular mesoscale organization with a significant subset exhibiting dynamic behaviors consistent with biomolecular condensates.

**Figure 2.**
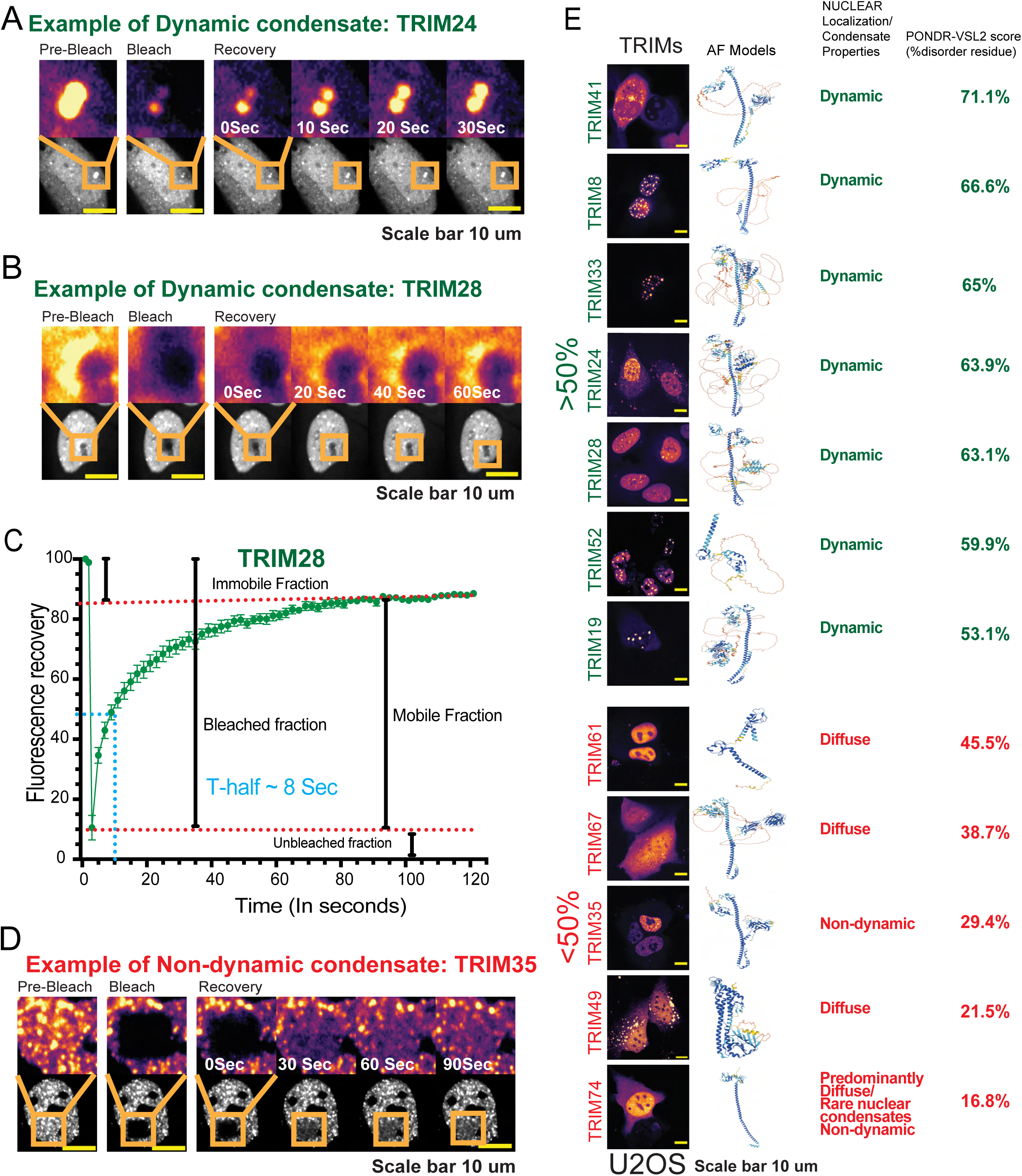
Systematic evaluation of TRIM dynamics by FRAP reveals nuclear TRIMs forming biomolecular condensates. **A** TRIM24 nuclear foci are shown dynamically recovering upon photo-bleaching. **B.** TRIM28 nuclear foci are shown dynamically recovering upon photo-bleaching. **C.** Fluorescence recovery of TRIM8 signal after photobleaching is shown reflecting the significant mobile fraction relative to the immobile fraction and the half-life for recovery. **D.** The representative profile for the non-dynamic TRIM35 nuclear foci is shown. **E.** Panel shows the relationship for TRIMs predominantly localizing to the nucleus between dynamic recovery from photo-bleaching for those forming condensates and their disordered content (based on PONDR-VSL2). Each row shows a representative image and predicted Alpha-fold structure for a given nuclear TRIM. Those TRIMs with disordered content >50% were noted to form dynamic nuclear condensates, while TRIMs with disordered content less than 50% were found to have diffuse localization within the nucleus or to form non-dynamic nuclear bodies.

To look at the relationship between dynamic behavior and IDR content, we compared the disorder content of TRIM family proteins predicted using PONDR-VSL2^29,30^ (**Figure 1A, Suppl Figure S12, Suppl Table ST1, ST3**). Interestingly, the disordered content of TRIM proteins forming dynamic bodies was significantly higher than that of TRIMs forming non-dynamic bodies (**Suppl Figure S12D**). However, there was clearly overlap in disordered content between dynamic and non-dynamic proteins. There were TRIM proteins that formed dynamic bodies (7/19) with disordered content below 50% as well as family members with disordered content above 50% that organized into non-dynamic bodies (3/19).

Nonetheless, the relationship between disordered content and dynamic behavior was most apparent for TRIMs forming nuclear bodies, for which the disordered content of the dynamic TRIM proteins (TRIM8, −19, −24, −28, −33, −41, −52) ranged from 53.1-71.1% whereas that of the non-dynamic TRIMs (TRIM35, −74) were 16.8-29.4% (**Figure 2E**, **Suppl Figure S5, S9, S10, S11, S12, Suppl Table ST3**). (TRIM74 rarely forms nuclear condensates and typically appeared diffuse in the nucleus and cytoplasm.) In the remaining three TRIMs with any predominant nuclear localization, these displayed diffuse nuclear localization without nuclear bodies and had disordered content less than 50% by PONDR-VSL2 (**Figure 2E, Suppl Table ST3**). Similar trends were observed using other *in silico* prediction tools for disordered content (PICNIC, FuzDrop-pLLPS)^31,32^ (**Suppl Figure S12**). These analyses suggested a potential relationship between IDR content of the TRIMs and their ability to form biomolecular condensates, particularly in the nucleus.

### Emergent mesoscale properties of TRIM nuclear condensates associated with molecular complexity

We can illustrate the relationship between the molecular complexity (e.g. IDR content and modular structural complexity) of TRIM family proteins and emergent mesoscale physicochemical properties by examining four nuclear proteins that robustly form condensates: TRIM52, TRIM8, TRIM19/PML and TRIM33 (**Figure 3A, Suppl Figure S5, S9, S10, Suppl Table ST1**). TRIM52 has simple domain complexity with a RING finger domain, Type 2 B-Box domain and an IDR. TRIM8 has a complete RBBC motif and C-terminal IDR. Relative to TRIM8, TRIM19/PML has an additional SUMO interaction motif on its C-terminus, while TRIM33 has an additional C-terminal PHD-BROMO domain that interacts with chromatin. (As a side note, TRIM19/PML molecular complexity is further diversified by its alternative splicing as shown in **Figure S13**.) Therefore, these four TRIMs represent a ladder of complexity with TRIM8 having intermediate complexity (**Figure 3A**).

**Figure 3.**
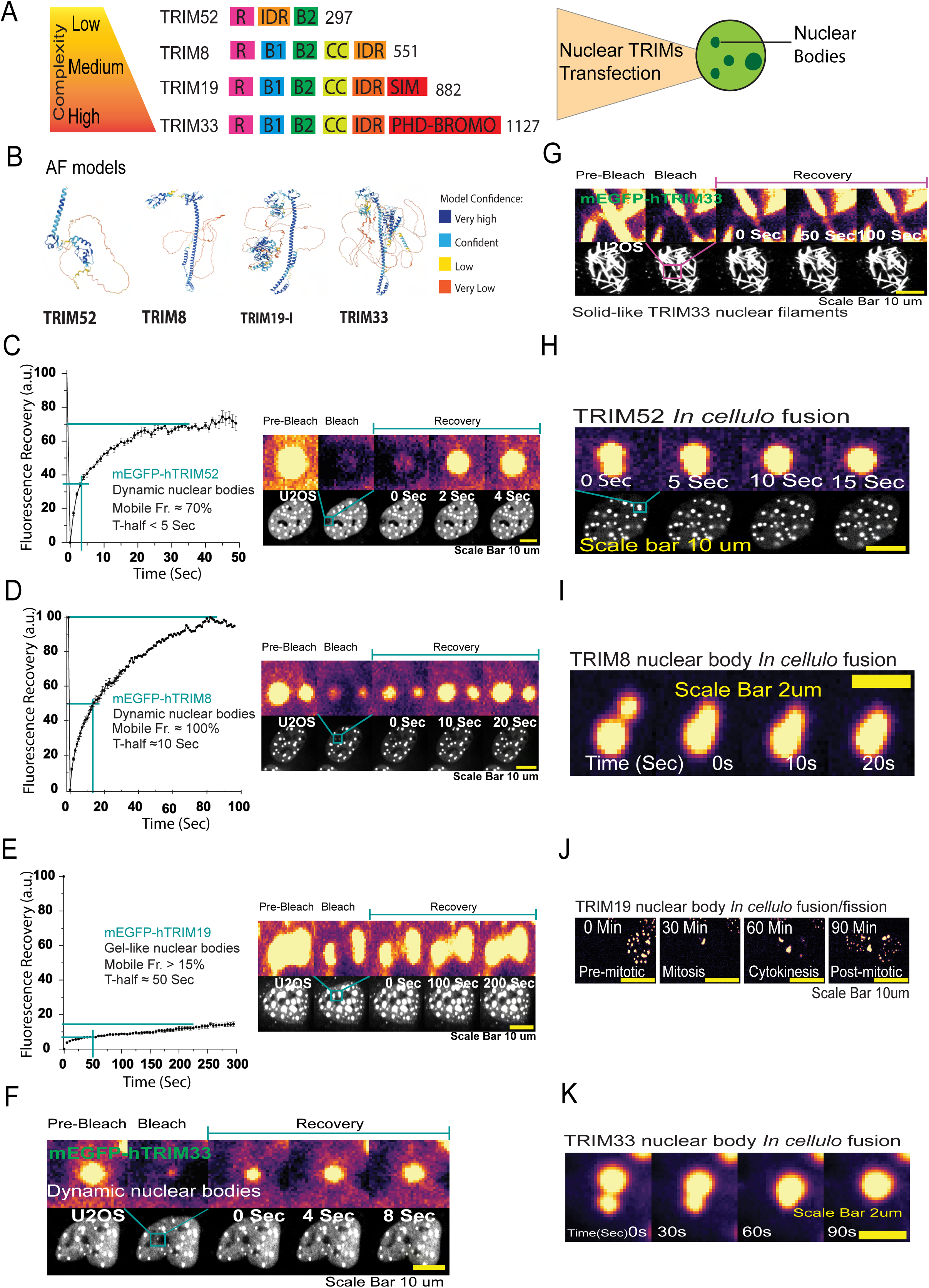
TRIM family nuclear proteins display complexity dependent condensate organization. **A** Panel shows the natural domain architecture for the nuclear TRIM family proteins (TRIM8, −19, −33, 52), in a hierarchically increasing complexity. **B.** Alpha fold model of TRIM52, TRIM8, TRIM19 and TRIM33 are shown. **C.** Live cell imaging of the TRIM52 nuclear bodies is shown upon fluorescently labeled TRIM52 encoding plasmids were transfected in U2OS cells and imaged after 24 hours. Dynamic recovery after photo-bleaching is shown for TRIM52 within seconds to minutes. **D.** Evaluation performed as in (C) for TRIM8 nuclear bodies, showing similar dynamic recovery. **E.** Evaluation performed as in (C) for TRIM19 nuclear bodies, showing dynamic recovery albeit less rapid than for TRIM8 and TRIM52. **F.** Evaluation performed as in (C) for TRIM33 nuclear bodies, showing dynamic recovery. **G.** Evaluation performed as in (C) for TRIM33 nuclear filaments, showing no recovery. **H.** Fusion/fission behaviors shown for TRIM52 over seconds to minutes. **I.** Fusion/fission behaviors shown for TRIM8 as in **H.** **J.** TRIM19 fusion (mitosis) and fission(cytokinesis) occurring with cell cycle over minutes to hours. **K.** Fusion/fission behaviors shown for TRIM33 as in **H.**

As part of our screen, we characterized these 4 nuclear body-forming TRIMs with respect to (1) diversity of the mesoscale foci observed in cells, (2) dynamics in the FRAP assay (**Figure 3C-G**), and (3) Fusion of the mesoscale foci (**Figure 3H-K**). With respect to the diversity of the condensate foci we observed a clear correlation between the increase in molecular scale complexity and diverse kind of mesoscale localization patterns. TRIM52, TRIM8 and TRIM19/PML always form round shaped condensates (**Figure 3C-E**). However, TRIM33 forms both condensates and filaments in the nucleus (**Figure 3F-G**). TRIM52 and TRIM8 show fast recovery rates (**Figure 3C-D**) while TRIM19/PML shows less dynamic behavior (**Figure 3E**). TRIM52 and TRIM8, similarly, showed fusion events over seconds consistent with liquid-like behaviour (**Figure 3H-I**), while TRIM19/PML exhibited fusion events over minutes to hours during the cell cycle (**Figure 3J**). TRIM33, on the other hand, showed a range of behaviors from foci that displayed fast dynamics (**Figure 3F**) with fusion events (**Figure 3K**) to filaments that did not show any recovery over the course of the measurement (**Figure 3G**). Taken together, these data suggest that molecular complexity and, in particular, the domain context surrounding IDRs can dictate emergent properties at the mesoscale via biomolecular condensation.

So far, we have correlated the molecular scale complexity of TRIM family proteins with mesoscale complexity at a variable range of protein concentrations using transient transfection. However, the localization pattern of any protein will likely depend on its precise bulk concentration. In addition, protein concentration is known to vary many folds in in different cell types, in disease, in different developmental stages. To examine the relationship between protein concentration and localization, we decided to construct phase diagrams of localization as a function of protein concentration. For this purpose, we generated a knockout (KO) of a gene of interest using CRISPR/Cas9 technology and transiently transfect the plasmids that encode the particular fluorescently tagged protein, for which the KO was generated, providing a range of protein concentrations. We, then, correlate the localization with the measured protein concentration of >300-1000s individual cells (**Figure 4A, Suppl Figure S14**). We observed that TRIM8 and TRIM33 show an increase in condensate appearance as a function of their concentration with fixed enrichment ratios while TRIM19/PML shows condensates at all detectable concentrations increasing enrichment (**Figure 4B, Suppl Figure 15A**). This demonstrates that TRIM molecular complexity results in distinct condensate properties.

**Figure 4.**
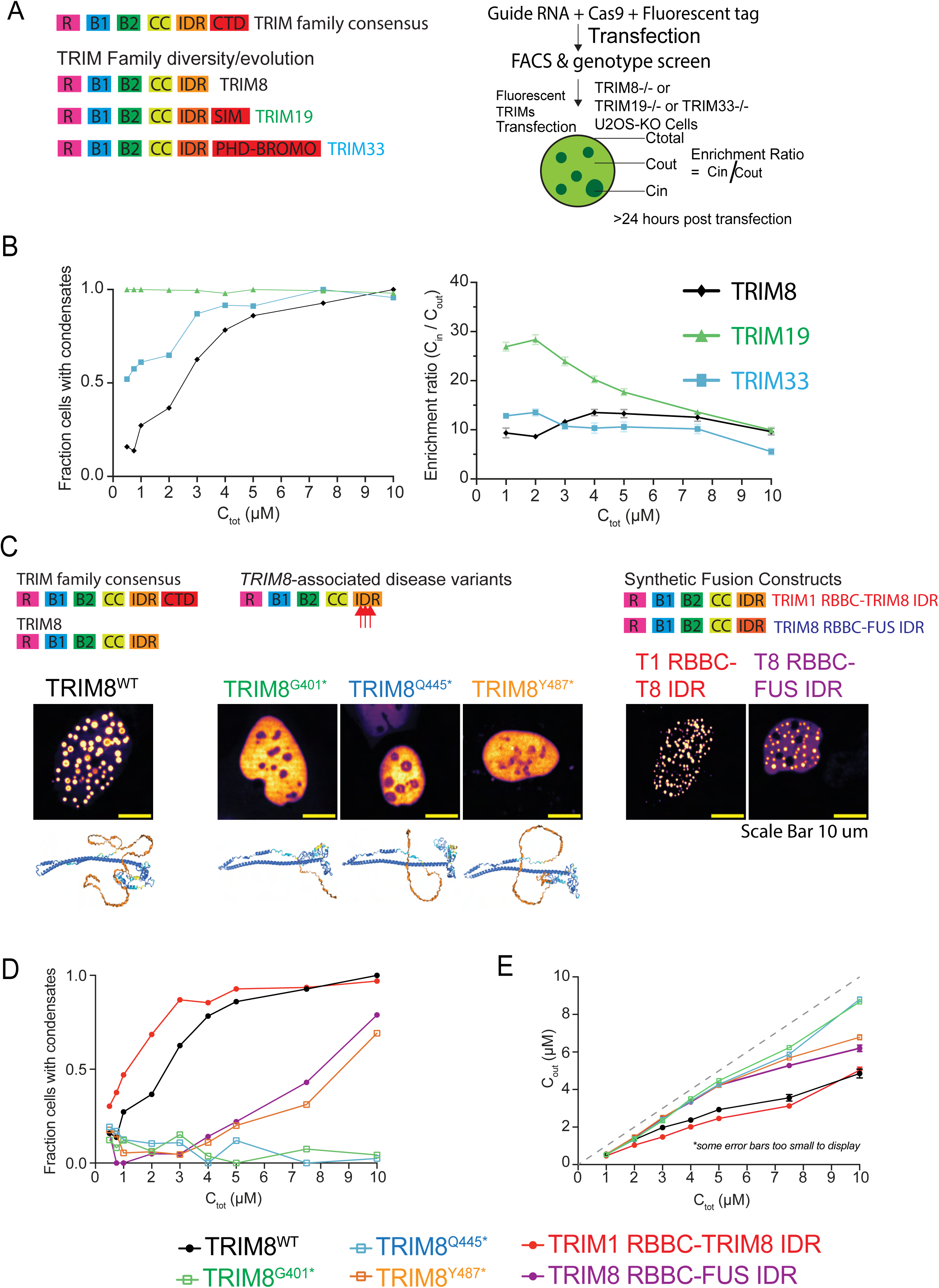
Phase diagrams reveal divergent features of TRIM nuclear condensates and indicate *TRIM8-*associated neuro-renal syndrome is a condensatopathy. **A** Domain architectures of the Nuclear TRIM proteins (TRIM8, 19 and 33) in an increasing order of complexity and the protein chain length (left), and on the right an outline for the nuclear TRIM protein condensation as a function of their concentration **(see also Suppl Figure S14)**. U2OS-KO cell lines of the corresponding proteins were transfected with fluorescently labelled TRIM8, 19 and 33 plasmids and the *in cellulo* concentrations were quantified. **B.** On the left shows a fraction of cells that form condensates as a function of protein concentration. While TRIM8 and TRIM33 condensates form as function of protein concentration, TRIM 19 shows constitutive condensates at all detectable concentrations. On the right shows, enrichment ratio (partition coefficient correlate) as a function of the protein *in cellulo* concentration, while TRIM8 shows a typical concentration dependent saturation-like behavior, meaning as the total concentration of TRIM8 increases, so does the concentration in the condensed phase. On the other hand, TRIM19 and TRIM33 show decrease in the enrichment ratio, meaning a much quicker ripening and therefore an increase in the dilute phase concertation. **C.** Domain architectures of wildtype TRIM8, patient mutations, and synthetic constructs are shown. Below the schema are representative images indicating disease variants disrupt formation of condensate. Synthetic constructs indicate that the TRIM8 IDR is sufficient to promote condensation but can be rescued with an alternative IDR. **D.** On the left, shows the fraction of cells with TRIM8 condensates. This shows that TRIM8 disease mutants cause loss of protein condensation, that the TRIM8 IDR can support condensation in a condensate-defective TRIM1 construct and that the TRIM8 IDR function can be rescued partially by the alternative FUS IDR. On the right, a plot displays the resulting increase in the dilute phase protein concentration by disease mutants. The TRIM8 IDR is sufficient to restore a low dilute phase concentration in condensate-defective TRIM1 and can be partially rescued by an alternative FUS IDR.

### Pathogenic variants *of TRIM8 IDR results in a loss of condensation*

We used this approach to quantitatively characterize the pathogenic variants in *TRIM8*-associated neurorenal syndrome^21–27^. These variants reduce the length of the TRIM8 IDR length and cause an expected change in its localization in a variety of constructs generated by deletion of the C-terminus coding sequence or introduction of patient variants, consistent with our previous observations^23^ (**Figure 4C, Suppl Figure 15B**). We evaluated three disease-causing TRIM8 variants (G401*, Q445*, and Y487*) in more depth (**Figure 4C**). The two earlier truncation mutants (G401*, Q445*) fail to form discernable condensates across a wide range of concentrations and cause marked increases in the dilute phase protein concentration. The shortest truncation (Y487*) causes an intermediate degree of condensate formation and an increase in dilute protein phase concentration relative to tagged wildtype TRIM8 (**Figure 4D-E**). This supports that patient variants cause defective phase separation through IDR loss.

To verify the loss of condensation was caused specifically by loss of an IDR, we performed complementation studies by fusing TRIM8 amino acids 1-401 to the established FUS IDR^6^ (TRIM8 RBBC-FUS IDR) (**Figure 4C-E**). Indeed, this fusion partially restores condensate formation and reduces the dilute protein phase concentration. Lastly, to ask whether TRIM8 IDR is sufficient to induce condensate formation, we fused the N-terminal RBBC domains of TRIM1 (amino acids 1-401) to the TRIM8 IDR with a nuclear localization signal (TRIM1 RBBC-TRIM8 IDR) (**Figure 4D**). TRIM1 itself normally has a filamentous localization (**Figure 1D**). As predicted, addition of the TRIM8 IDR to the TRIM1 RBBC domain was sufficient to induce concentration-dependent condensate formation and appropriately reduce the dilute protein phase concentration (**Figure 4C-E**).

### Loss of TRIM8 condensation leads to aberrant protein quality control

We next determined the cellular consequences of loss of TRIM8 condensation. TRIM family ligases are known to undergo proteasome-mediated degradation in parallel with substrate modification^33,34^. Therefore, we hypothesized that TRIM8 concentration, and thereby condensation, is regulated by the cellular protein quality control systems (**Figure 5A**).

**Figure 5.**
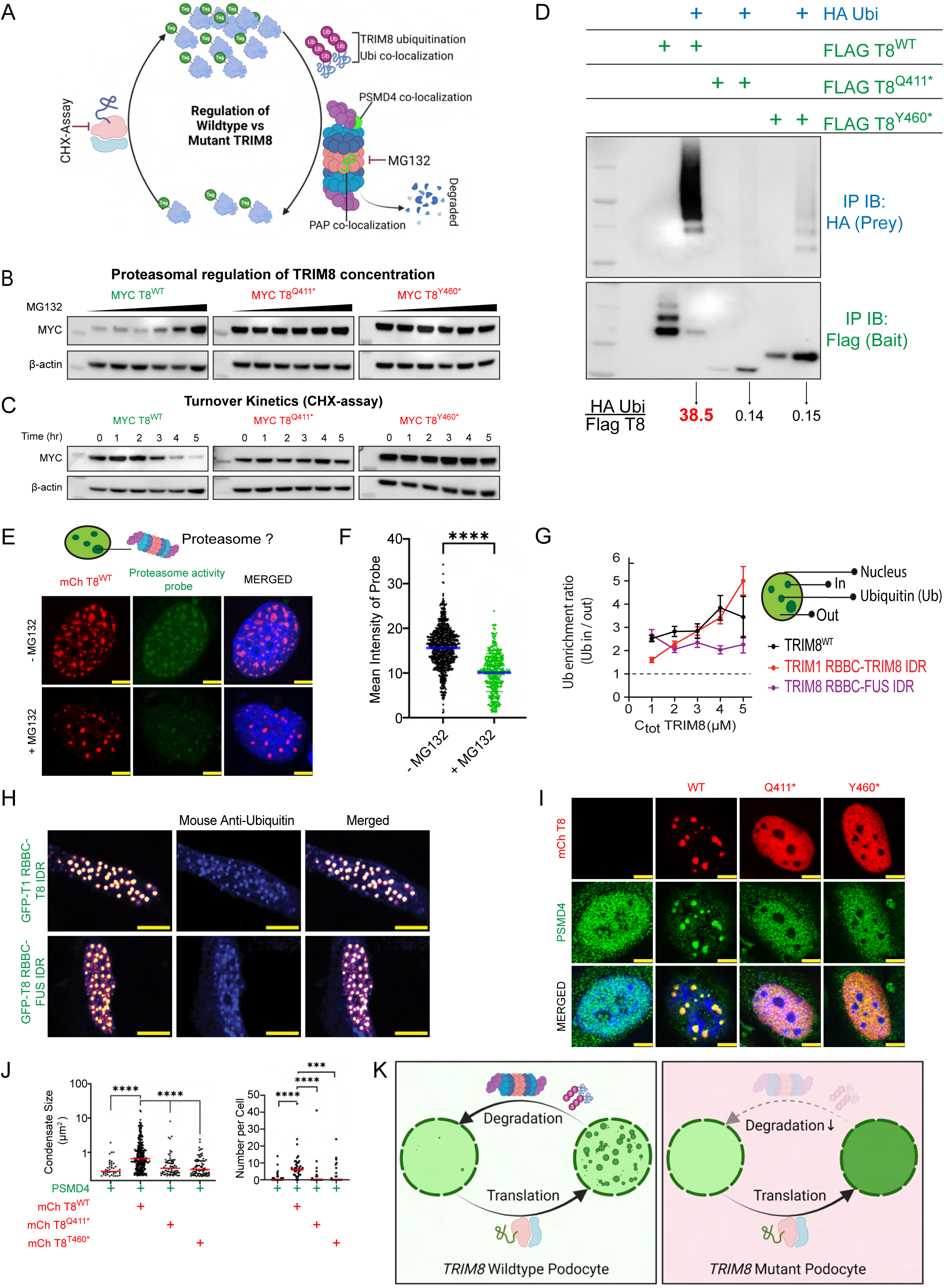
Loss of TRIM8 condensation leads to aberrant protein quality control. **A** Schematic diagram of the TRIM8 regulation in cells illustrating ubiquitination and proteasomal degradation. TRIM8 ubiquitination is studied by IP and co-localization and proteasomal degradation is demonstrated via the co-localization of proteasome components and proteasome activity probe (PAP) (Me4BodipyFL-Ahx3Leu3VS). The cycloheximide chase assay is performed to study the half-life of the protein. **B.** pIND-MYC TRIM8 wildtype or patient variants were induced by doxycycline and treated with 0-1µM MG132. Protein levels of MYC-tagged wildtype TRIM8 but not patient variant TRIM8 were increased by the 26S proteasome inhibitor MG132 in dose-dependent manner, while TRIM8 patient variant protein levels were unaffected by MG132 treatment, indicating a loss of proteasome regulation. Β-actin levels demonstrate equal loading **C.** Conversely, in the setting of protein translation inhibitor cycloheximide, tagged wildtype TRIM8 protein levels decrease within five hours while patient variant TRIM8 levels are preserved. **D.** Flag-tagged wildtype or patient variant TRIM8 was co-expressed with HA-tagged ubiquitin and immunoprecipitated to assess the ubiquitinated fraction of TRIM8. immunoblotting for HA revealed increased poly-ubiquitination of wildtype TRIM8 compared to patient variant TRIM8 when normalized to tagged TRIM8 levels. (See **Suppl Figure S16E** for input samples.) **E.** Cherry-tagged wildtype TRIM8 promotes localization of Proteasome Activity Probe (PAP) to nuclear condensates in immortalized podocytes, which is abrogated by MG132 treatment. **F.** The number of PAP nuclear condensates (H) is quantified, showing wildtype TRIM8 promotes the number of these condensates, while MG132 treatment lowers the condensate formation. **G.** Enrichment ratios for ubiquitin are shown, indicating there is enrichment for ubiquitin in wildtype TRIM8 condensates as well as condensates generated by TRIM8 synthetic constructs. **H.** Representative images are shown demonstrating co-localization of endogenous ubiquitin to synthetic constructs of TRIM8 indicating the TRIM8 IDR is sufficient to recruit ubiquitin to its condensates and can be rescued by an alternative FUS IDR. **I.** Cherry-tagged wildtype TRIM8 promotes localization of endogenous proteasome component PSMD4 to nuclear condensates in immortalized podocytes, which is abrogated by patient variants lacking the TRIM8 IDR. **J.** The number and size of PSMD4 nuclear condensates are quantified (I), showing wildtype TRIM8 promotes the number and size of these condensates, while patient variant TRIM8 do not. **K.** Schematic summary showing that in wildtype podocytes TRIM8 undergoes polyubiquitination and proteasomal degradation whereas mutant TRIM8 has reduced polyubiquitination and hence evades proteasomal degradation. Stabilization of wildtype TRIM8 shifts the localization to nuclear foci, whereas mutant TRIM8 retains pan-nuclear expression. The increased stability of mutant TRIM8 relative to wildtype TRIM8 represents a possible gain of function.

We first evaluated the impact of the 26S proteasome inhibitor MG132 on tagged-TRIM8 protein expression levels. TRIM8 protein was expressed ectopically in immortalized human podocytes, as a more disease relevant cell type, by transient transfection as well as by doxycycline-induced expression (**Figure 5B, Suppl Figure S16A**). Wildtype TRIM8 protein levels increased upon treatment with increasing concentration of MG132, suggesting its concentration is negatively regulated by the proteasome. In contrast, the protein levels of two independent mutant TRIM8 proteins, reflecting patient variants, were not reduced by MG132, indicating that truncation of the TRIM8 IDR by patient variants abrogates proteasome-dependent regulation. To further corroborate these results, we looked at the rate of degradation of TRIM8 using cycloheximide chase assay. A tagged wildtype protein levels exhibited a half-life of ∼3 hours, while patient mutant TRIM8 protein half-life is >5 hours (**Figure 5C, Suppl Figure S16B**). Therefore, we conclude that condensation competent wildtype TRIM8 concentration can be regulated by the ubiquitin-proteasome system (UPS), while the condensation incompetent disease mutants are not regulated.

Consistent with the results in immortalized cells, proteasome inhibition also increased condensate formation of tagged wildtype TRIM8 protein, while mutant protein remained in a pan-nuclear (dilute phase) pattern in immortalized podocytes (**Suppl Figure S16C-D**). Taken together our data so far show that the TRIM8 IDR regulates its (1) condensation, (2) half-life and (3) proteasomal degradation, all of which are disrupted by disease variants.

We further dissected the mechanism by which TRIM8 protein stability in physiology and disease is regulated by assaying differential polyubiquitination of TRIM8 and its variants. For this purpose, we performed co-immunoprecipitation assays upon co-expression of tagged TRIM8 with ubiquitin. We observed a characteristic smear of wildtype TRIM8 consistent with its polyubiquitination (**Figure 5D, Suppl Figure S16E**). This polyubiquitination was markedly reduced in two independent TRIM8 disease mutants, suggesting the IDR is mediating TRIM8 modification. We considered whether the critical lysine residues for ubiquitination were within the IDR after the truncation variants. However, mutation of lysine residues 464 and 546 did not alter proteasome-dependent regulation of TRIM8 (**Suppl Figure S16F-G**), suggesting other residues are the sites of polyubiquitination.

To look at whether TRIM8 condensates are sites of active ubiquitination, we treated cells with proteasome activity probe Me4BodipyFL-Ahx3Leu3VS^35,36^ and observed co-localization of functionally active proteasomes within TRIM8 condensates. This signal was impaired as anticipated by MG132 (**Figure 5E-F**). Consistently, ubiquitin colocalizes with TRIM8 condensates in U2OS cells as well as immortalized human podocytes, while this localization is abrogated by patient variants (**Suppl Figure S17A-B, S18A-D**). We, furthermore, evaluated ubiquitin localization with synthetic constructs TRIM8 RBBC-FUS IDR and TRIM1 RBBC-TRIM8 IDR, observing co-localization with overexpressed ubiquitin (**Suppl Figure S18E-F**) and that exhibited enrichment within TRIM8 condensates (**Figure 5G**). This co-localization was confirmed with endogenous ubiquitin (**Figure 5H**), indicating the TRIM8 IDR is sufficient to organize ubiquitin but can be complemented by alternative IDRs. Furthermore, endogenous components of the proteasomal machinery (PSMD4, PSMD12) localized to wildtype TRIM8 condensates and not in disease variants (**Figure 5I-J**, **Suppl Figure S17C-D**).

In summary our results suggest that, as TRIM8 protein concentration increases, the TRIM8 IDR promotes TRIM8 condensation, which recruits the UPS machinery to polyubiquitinate and degrade TRIM8 (**Figure 5K**). This negative feedback mechanism is disrupted by TRIM8-associated disease variants, resulting in a more stable protein that no longer forms nuclear condensates.

### Loss of TRIM8 condensation by pathogenic variants disrupts TAK1/NFκB signaling

Previous work has shown that TRIM8 interacts with and mediates K63-linked polyubiquitination of transforming growth factor β-activated kinase 1 (TAK1), thereby promoting its activity and downstream NFκB signaling^37–43^. Mutations in *Tak1* and *Nfkb1* cause defective podocyte homeostasis and response to injury^44,45^. This mirrors the human podocyte disorder caused by *TRIM8* variants. Therefore, we hypothesized that TRIM8 condensation is required for TAK1/NFκB signaling and that this signaling is disrupted by TRIM8 disease variants (**Figure 6A**).

**Figure 6.**
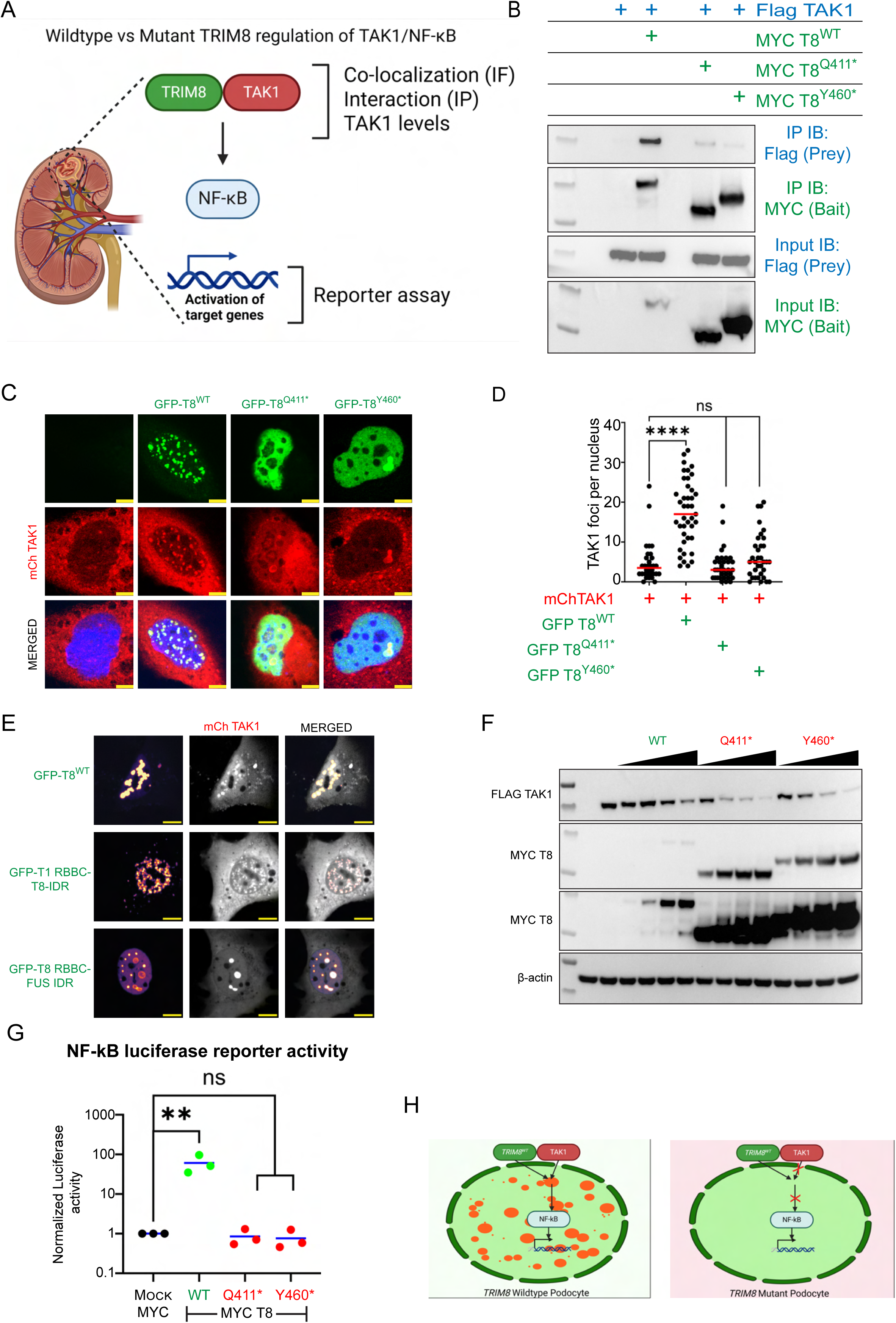
Loss of TRIM8 condensation by pathogenic variants disrupts TAK1/NFκB signaling. **A** Schematic diagram of NF-kB regulation in podocytes. **B.** MYC-tagged wildtype or patient variant TRIM8 was co-expressed with Flag-tagged TAK1 and immunoprecipitated. Input blots (10% of IP) confirm co-expression of MYC-tagged TRIM8 and Flag-tagged TAK1. immunoblotting of IP eluates for Flag-tagged wildtype TRIM8 revealed co-precipitation of TAK1, while tagged mutant TRIM8 exhibited reduced co-precipitation with TAK1, suggesting TRIM8 and TAK1 interact through an IDR-dependent manner. **C.** GFP-tagged wildtype TRIM8 promotes localization of Cherry-tagged TAK1 to nuclear condensates in immortalized podocytes, which is abrogated by patient variants lacking the TRIM8 IDR. **D.** The number of Cherry-tagged TAK1 nuclear condensates in (C) is quantified, showing wildtype TRIM8 promotes the formation of these condensates while patient variant TRIM8 does not. **E.** Representative images of synthetic TRIM8 constructs are shown co-localizing with the TAK1 and indicating that the TRIM8 IDR is sufficient to induce TAK1 recruitment but can be rescued in TRIM8 by an alternative IDR to mediate TAK1 recruitment. **F.** Western blot analysis of immortalized podocyte lysates was performed after co-transfection with Flag-tagged TAK1 and increasing amounts of MYC-tagged wildtype TRIM8 or patient variant TRIM8. TRIM8 patient variants exhibited higher expression levels than the wildtype protein. TAK1 levels were reduced in a dose dependent manner with increasing TRIM8 mutant protein, while only modest reduction in TAK1 levels was observed with wildtype TRIM8 protein. β-actin levels demonstrate equal loading. **G.** NF-κB transcriptional activity was assessed by a 4xNF-κB binding site firefly luciferase reporter in HEK293T cells with renilla luciferase co-transfected as an internal control. The ratio of firefly: renilla activity normalized to the level for the mock MYC-tagged plasmid transfection group of each of three biological replicates is shown. NF-κB is increased by co-transfection with wildtype TRIM8 while TRIM8 patient variants failed to modulate reporter activity, indicating this effect is dependent on the TRIM8 IDR. **p<0.01 by t-test. **H.** Schematic summary showing the interaction of TAK1 and TRIM8 in wildtype podocytes which is reduced in the mutant podocytes. TAK1 and TRIM8 co-localize in the wildtype podocytes (yellow dots) however the co-localization is abrogated in the mutant podocytes. Downstream signaling of TAK1 activation by TRIM8 activates the NF-κB activity in wildtype TRIM8 podocytes which is absent in the mutant TRIM8 podocytes.

We first tested the impact of disease mutants on TRIM8-TAK1 interaction using co-immunoprecipitation. We observed TRIM8 loss of IDR disease variants disrupted the TRIM8:TAK1 interaction in immortalized human podocytes (**Figure 6B**). Consistently, TAK1 co-localized to TRIM8 condensates, and this was abrogated by patient variants (**Figure 6C-D, Suppl Figure S19A**). We, also, observed co-localization of tagged TAK1 with synthetic constructs TRIM8 RBBC-FUS IDR and TRIM1 RBBC-TRIM8 IDR (**Figure 6E**), indicating the TRIM8 IDR is sufficient to recruit TAK1 but can be complemented by alternative IDRs. Lastly, co-expression of mutant TRIM8 and TAK1 led to reduced TAK1 levels (**Figure 6F**).

We tested the impact on the downstream NFκB signaling using luciferase reporter assays. As previously established, ectopic wildtype TRIM8 expression in multiple kidney cell lines induced NFκB reporter activity, which was abrogated by two independent patient variants (**Figure 6G, Suppl Figure S19B**). In conclusion, condensate incompetent TRIM8 disease variants impair interaction with and recruitment of TAK1, cause reduced TAK1 levels, and result in downstream failure to induce NFκB signaling (**Figure 6H**).

### Cellular and Organismal effects of TRIM8 pathogenic variants in cell culture and in animals

We postulated that *TRIM8* disease variants cause kidney disease in humans by causing podocyte dysfunction at the cellular and organismal levels (**Figure 7A**). CRISPR/Cas9 mediated gene editing was performed in immortalized human podocytes to generate homozygous N-terminal truncating variants within exon 1 to cause loss-of-function (*TRIM8^Ex1-^*) or homozygous C-terminal truncating variants within exon 6 that mimic patient variants and truncate the TRIM8 IDR (*TRIM8^Ex6-^*) (**Suppl Figure S20A-G**). Immunoblotting of clarified lysates from wildtype and *TRIM8* mutant cell lines confirmed loss of TRIM8 expression in *TRIM8^Ex1-/Ex1-^* podocytes and truncated TRIM8 protein in *TRIM8^Ex6-/Ex6-^* podocytes (two independent cell lines per genotype) (**Suppl Figure S20H-J**).

**Figure 7.**
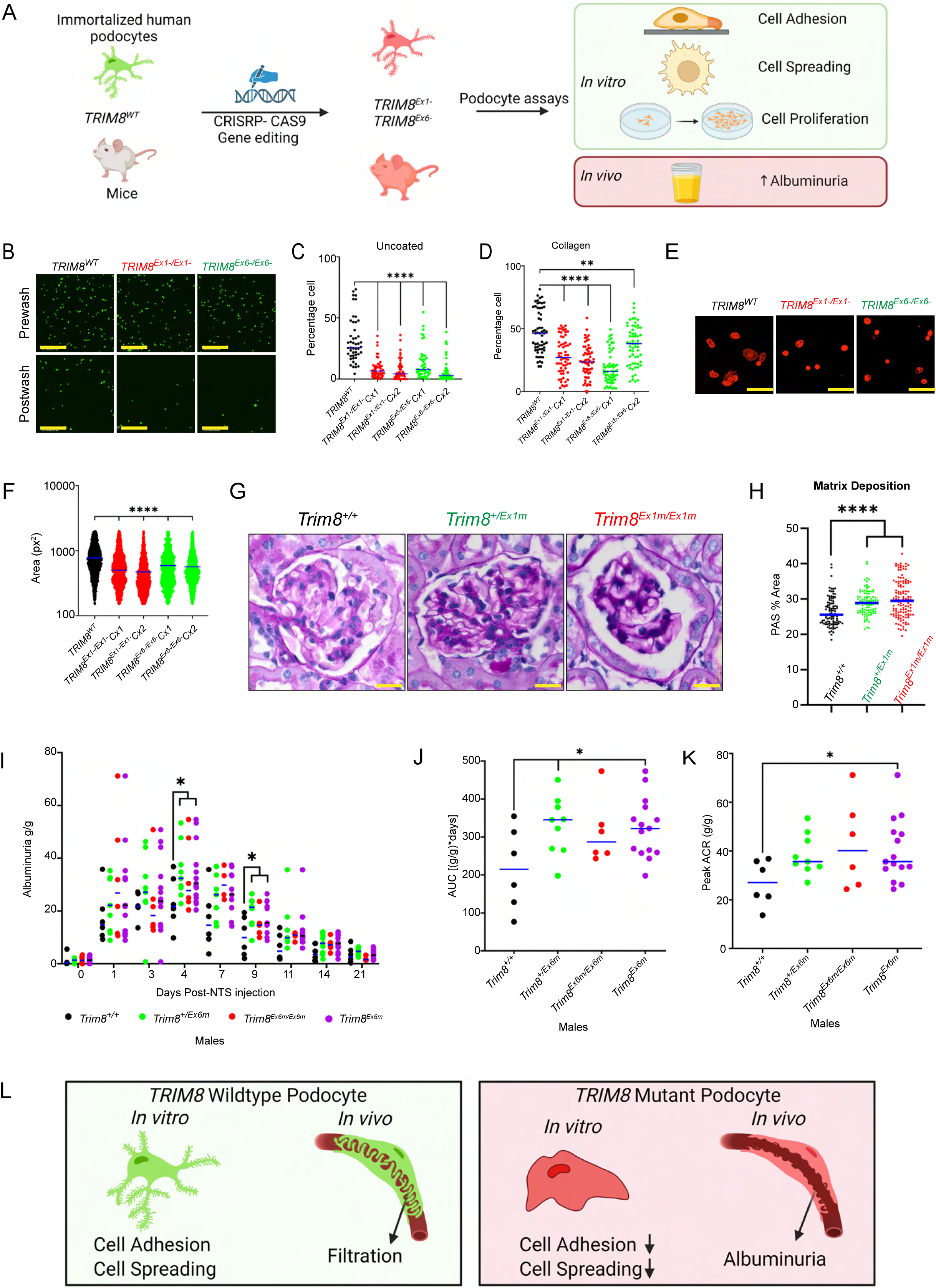
TRIM8 regulates podocyte adhesion and spreading in vitro and response to podocyte injury in vivo. **A** Schematic diagram of generation of CRISPR mutant cells and mice. Different podocyte assays such as cell adhesion, cell spreading, and cell proliferation were performed using wildtype and mutant cells. The CRISPR-generated mice underwent NTS injury and urine was collected and analyzed. **B.** Immortalized human podocytes bearing bi-allelic TRIM8 variants causing N-terminal truncation (*TRIM8^Ex1-/Ex1-^*^)^ or C-terminal truncation (*TRIM8^Ex6-/Ex6-)^* were generated by CRISPR/Cas9 gene editing. **C.** Mutant cells labeled with a green fluorescent probe (calcein AM) exhibited reduced cell adhesion to tissue culture-treated dishes relative to wildtype immortalized podocytes. ****p<0.0001 by Mann Whitney U-test. **D.** Immortalized human podocytes bearing bi-allelic TRIM8 variants causing N-terminal truncation (*TRIM8^Ex1-/Ex1-^*) or C-terminal truncation (*TRIM8^Ex6-/Ex6-^*) were generated by CRISPR/Cas9 gene editing. Mutant cells labeled with a green fluorescent probe (Calcein AM) exhibited reduced cell adhesion to collagen-coated, tissue culture treated dishes relative to wildtype immortalized podocytes. **p<0.01, ****p<0.0001 by Mann Whitney U-test. **E.** Immortalized human podocytes bearing bi-allelic TRIM8 variants causing N-terminal truncation (*TRIM8^Ex1-/Ex1-^*) or C-terminal truncation (*TRIM8^Ex6-/Ex6-^*) were evaluated for altered cell spreading relative to wildtype control cells. **F.** Mutant cells whose surface was labeled with wheat germ agglutinin exhibited reduced cell spreading relative to wildtype immortalized podocytes. ****p<0.0001 by Mann Whitney U-test. **G.** Periodic acid–Schiff (PAS) stained sections of kidneys from 12-month-old *Trim8^Ex1m-/Ex1m-^*, *Trim8^Ex1m -/+^* and *Trim8^Ex1m +/+^* mice were generated. Representative images of PAS from each genotype are shown. **H.** 20x images of glomeruli were evaluated through an automated ImageJ pipeline to determine glomerular matrix deposition in a blinded manner. *Trim8^Ex1m-/Ex1m-^* and *Trim8^Ex1m -/+^* mice showed significantly increased glomerular matrix deposition. Mann-Whitney test; ***, p<0.001. **I.** C-terminal truncating TRIM8 variants were generated in FVB/N mice (*Trim8^Ex6m^*). Podocyte injury was induced with sheep nephrotoxic serum (NTS). Albuminuria was assessed to follow injury and recovery. Male heterozygous mice exhibited increased albuminuria relative to wildtype mice at 4 and 9 days after NTS treatment, while homozygotes did not exhibit significantly different albuminuria at any timepoints. *p<0.05 by Mann-Whitney U-test. **J.** In the NTS study in (I), total albuminuria defined as the area-under-curve was significantly increased in male heterozygotes (*Trim8^+/Ex6m^*) and collectively all male mutant animals (*Trim8^+/Ex6m^* and *Trim8^Ex6m/Ex6m^*) relative to littermate wildtype male control mice. **K.** In the NTS study in (I), peak albuminuria was significantly increased collectively for all male mutant animals (*Trim8^+/Ex6m^* and *Trim8^Ex6m/Ex6m^*) relative to littermate wildtype male control mice. **L.** Schematic summary showing mutant TRIM8 podocytes have reduced cell adhesion and spreading relative to the wildtype TRIM8 podocytes. The TRIM8 mutant mice also have elevated response to NTS injury compared to the wildtype TRIM8 mice.

We assessed two-dimensional cell growth using live cell imaging in different seeding densities. Upon seeding at lower density, *TRIM8* mutant cell lines exhibited increased cell number relative to wildtype immortalized podocytes, while no significant difference in cell number was observed when seeding at higher densities (**Suppl Figure S20K-N**). Cell proliferation was additionally assessed by MTT assay. No consistent significant differences were noted at low density (**Suppl Figure S20G**), while only *TRIM8^Ex1-/Ex1-^* podocytes exhibited consistently reduced proliferation at higher density. Overall, *TRIM8* truncating variants led to modest effects on cell growth and proliferation. We then turned to podocyte adhesion assays, because cell adhesion has been shown to be positively regulated by TAK1/NFκB signaling and implicated in podocyte disease^46,47^. All *TRIM8* mutant podocyte cell lines exhibited reduced cell adhesion relative to control wildtype podocytes (**Figure 7B-D**) and reduced cell spreading (**Figure 7E-F**), strongly arguing that *TRIM8* disease variants cause podocyte dysfunction.

Finally, we generated germline variants in *Trim8* in mice in order to assess the effects of these variants on podocyte homeostasis in animals. As with the immortalized podocytes, CRISPR/Cas9 gene editing was employed to generate N-terminal truncating variants within exon 1 to cause loss-of-function (*Trim8^Ex1m^*) or homozygous C-terminal truncating variants within exon 6 that mimick patient variants and truncate the TRIM8 IDR (*Trim8^Ex6m^*) (**Suppl Figure S21A-F, S22A**). Both heterozygous and homozygous *Trim8* mutant mice, exhibited normal survival, allowing us to assess age-related kidney disease (**Suppl Figure S21G-H, S22B-C**).

Therefore, we assessed podocyte homeostasis and dysfunction using proteinuria as a readout by measuring urine albumin-to-creatinine ratios in monthly collections. We observed that proteinuria was not significantly different between *Trim8* mutant mice and littermate controls through age 12 months (**Suppl Figure S21I, S22D**). At age 1 year, *Trim8* mutant mice and littermate controls were euthanized, and tissues were harvested. Kidney sections were stained using Periodic acid Schiff (PAS) to assess glomerular matrix deposition as a marker of chronic glomerular injury. *Trim8^Ex1m^* heterozygotes and homozygotes exhibited significantly increased matrix deposition relative to wildtype controls (**Figure 7G-H**), while the glomerular matrix was not significantly different between *Trim8^Ex6m^* mutant mice and wildtype controls (**Suppl Figure S22E-F**). This strongly argues that N-terminal truncating variants in *Trim8* cause mild chronic glomerular injury *in vivo*, consistent with the *TRIM8* syndrome phenotype in humans.

Because *Trim8^Ex6m^* mutant mice failed to develop a homeostatic phenotype, we posited that they may have an impaired response to podocyte injury in line with *Nfkb1* mutant mice^44^. We, therefore, employed sheep-derived nephrotoxic serum (NTS) with antibodies targeting podocytes to induce transient podocyte injury^48–52^. As a marker of podocyte injury and recovery, albuminuria was assessed at timed intervals through 21 days post-injection. Male heterozygote *Trim8^Ex6m/+^* mice exhibited higher albuminuria at 4 and 9 days post-injection and overall by area-under-curve (AUC) analysis (**Figure 7I-J**). All male mutant mice (heterozygotes and homozygotes collectively) exhibited increased albuminuria by AUC and peak albuminuria (**Figure 7J-K**). Female mice overall had more modest injury in response to NTS with lower albuminuria across genotypes, and no significant differences were observed between mutant mice and wildtype controls (**Suppl Figure S22G-I**). In conclusion, TRIM8 disease variants caused prolonged podocyte injury, in particular, in male heterozygote mice again consistent with the *TRIM8-*associated disease phenotype.

## DISCUSSION

As the human genome and subsequent annotation of molecular-scale features of the human proteome^53,54^ has revealed the diversity of the cellular protein landscape, it has also opened questions of how these molecular features result in the complex mesoscale organization within cells and how this may be perturbed in disease. The role of biomolecular condensation broadly in this organization remains critically unexplored. Here, we characterize the mesoscale cellular topography of TRIM family proteins and provide a multi-scale framework for exploring biomolecular condensation as a mechanism for this organization, leading to the molecular dissection of a human genetic condensatopathy.

Our TRIM screen identified that a majority of 72 of the TRIM family proteins exhibit a predominant mesoscale organization and to some extent localize to cellular bodies when expressed in U2OS and/or HELA cells (**Figure 1**). Systemic evaluation of fluorescence recovery of TRIM family cellular bodies revealed that a significant fraction are dynamic biomolecular condensates, suggesting this is a common mechanism for cellular compartmentalization (**Figure 2**). That TRIM cellular bodies were observed in a diversity of localization patterns underscores the potential for biomolecular condensates to mediate specialized functions across cellular compartments.

The diversity of mesoscale patterns exhibited by TRIM proteins is likely attributable to the highly multivalent nature of this protein class consisting of an RBBC motif, variable ordered C-terminal domains mediating polymer as well as IDR regions **(Figure 1A-B)**^8–12^. We provide a model example by comparing the mesoscale properties of a series of dynamic nuclear condensate forming TRIMs (8, 19, 33, 52) and demonstrating that increasing modular complexity through additional ordered domains can yield emergent cellular properties (**Figure 3-4**).

Thus, through cellular landscaping of the TRIM protein family, we provide evidence that the molecular features driving condensation (i.e. IDR) should be integrated into the annotations and larger knowledge of the proteome. This should occur through not only *in silico* tools to predicted disordered content but also through functional validation as we have displayed here. This is critical because, while disordered content does correlate with dynamic condensate behavior (**Figure 2D, Suppl Figure S12D**), it did not exclusively explain with the dynamism of TRIM proteins.

The need for broad evaluations of condensate driving forces in health and disease is best illustrated by our findings on TRIM8. We explore TRIM8 biomolecular condensates, as naturally occurring disease variants cause shortening of its IDR and aberrant phase separation (**Figure 4**). This has significant molecular consequences for the organization of protein quality control systems within these condensates (**Figure 5**) and TAK1/NFkB signaling which is critical to podocyte function^44,45^ (**Figure 6**). We, furthermore, evaluate the role of *TRIM8* variants on immortalized human podocyte function *in vitro* and mouse podocyte function *in vivo,* observing that these variants lead to defective cultured podocyte adhesion and aberrant responses to podocyte injury in mice (**Figure 7**). Collectively, these findings not only establish a direct link between IDR-driven condensation and human genetic disease.

While we posit that these molecular functions of TRIM8 are disrupted by loss of condensation in disease, future studies should explore if the cellular and organismal level phenotypes caused by *TRIM8* variants can be rescued through complementation of IDR function (e.g. TRIM8 transgenes with alternative IDRs or activation of downstream signaling). To support these studies, it will be important to establish under what developmental stages and/or injury settings that endogenous TRIM8 forms condensates.

These observations suggest that *de novo TRIM8* variants are complex and may cause disease through loss-of-function (aberrant recruitment of TAK1, stimulation of NFkB signaling, and aberrant cell adhesion), gain-of-function (increased stability of mutant TRIM8 protein), or aberrant function (reduced TAK1 levels). These findings must be reconciled with more N-terminal truncating variants in control subjects in the gnomAD database^23^, which would presumably cause haploinsufficiency. Our molecular findings also need to be considered in the context of the modest phenotypes in knockout *Trim8* mice (**Figure** 7) and *trim8a/b* mice^23^, although these observations should be interpreted with caution as there are compensatory mechanisms that could explain these organism level observations^55^ and other mouse models of genetic kidney disease have shown mild phenotypes^56,57^.

Additionally, through complementation studies, we provide evidence that TRIM8 IDR-dependent functions can be transferred to other paralogues and that another functional IDR can rescue certain TRIM8 molecular functions (**Figure 4D-E, 5G-H**, and **6E**). These observations raise questions about the specificity and grammar underlying IDR-directed phase separation.

An important part of our study is the creation of condensation diagrams for a subset of nuclear TRIMs. The question of localization will depend centrally on the concentration of a protein, and this will vary according to disease, cell type and cell cycle. Our approach suggests that a primary screen should be followed up by more detailed condensation diagrams.

In summary, this study establishes a robust framework for linking biomolecular condensation to mesoscale organization, protein regulation, and disease. By leveraging the TRIM family as a model system, we provide insights into the biophysical and functional principles governing condensate behavior and lay the groundwork for future exploration of *condensatopathies*.

## Acknowledgements

HRS thanks NOMIS foundation for funding. HRS thanks Technology development studio (TDS) especially Rico Barsacchi, Martin Stoeter, Nadine Tomschke and Claudia Moebius for technical support. HRS thanks LMF especially Britta Schroth-Diez, Catarina Nabais, Jan Peychl for technical support, Genome Engineering Facility (Ilka Reichardt-Gómez and Mihail Sarov) for generating TRIM33 and TRIM8 KO cell lines. HRS thanks Regis Lemaitre from the PEPC for providing purified mEGFP and mCherry used to make standard curve for the *in cellulo* protein concentration quantification. HRS thank Christina Eugster Oegama from the MPI-CBG central facility (OSCF) for providing routinely Mycoplasma tested and authenticated U2OS/HELA cell line used for this study. HRS thank Aliona Bogdanova for providing plasmid vectors for cloning. HRS thanks Julia Jarrells from the MPI-CBG, FACS facility. HRS thanks Paul J Taylor and Louis Flamand for the U2OS TRIM25 and TRIM19 KO cells. HRS thanks Adolfo garcia-sastre for many TRIM inserts containing plasmid DNA and Peter Dittrich for TRIM19 plasmids (please also see key resource table). HRS thank Emma Lundberg for the HPA antibodies.

A.J.M. was supported by the NIH (5K12HD052896-13, 1K08DK125768-01A1), American Society of Nephrology (Norman Siegel Research Scholar Career Grant 81542), Manton Center for Orphan Disease Research (Junior Faculty Award). This work was supported by a Boston Children’s Hospital Office of Faculty Development/Basic and Clinical Translational Research Executive Committees Faculty Career Development Fellowship (A.J.M). F.H. is the William E. Harmon Professor of Pediatrics and has grant support from the National Institutes of Health (5R01DK076683-13).

## MATERIALS AND METHODS

### Molecular scale methods analysis of the TRIM family proteins

#### Molecular scale *in silico* analysis of the TRIM family proteins

All alpha fold models of TRIM proteins were downloaded from the Alpha fold server^58,59^ and prepared using pymol for representation and analysis (**Suppl Figure S1**, **Suppl Files SF2-SF78**). Alpha fold3 server^60^ was used to visualize domain structure and interactions (e.g. between NHL repeats containing TRIMs with the RNA, Modified histone octamer and the Chromatin binding TRIMs, TRIM19-SIM and the Poly-Sumo interactions). It was also employed for the modelling of the effect of TRIM8 variants on its protein structure.

#### Mammalian cell culture

Both U2OS and HELA cells were grown in high glucose DMEM medium (Gibco-31966021) that was supplemented with both, 10% FBS (Sigma-S0615) as well as 100 units/ml Penicillin-Streptomycin (Gibco-15140112). Cells were seeded on the Ibidi 8 well imaging chambers (80826) or the 96 well plates, following trypsinization with Trypsin-EDTA (Gibco-25300054. Cells were grown overnight, then transfected with plasmids encoding the protein of interest and for the further colocalization purposes with other protein of interest encoding plasmids using the Fugene HD transfection reagent (Promega-E2311).

#### Plasmid constructions, fluorescent protein expression in mammalian cells

U2OS/HELA cells were transfected with plasmids encoding either TRIM/other proteins tagged with florescent tags at either the N-terminus or indicated otherwise. Some of the genes were codon optimized (for synthesis ease as well as to override any cellular regulation involving mRNA degradation of endogenous sequences) using synthesized g-blocks from Integrated DNA Technologies (IDT), Twist Biosciences and GenScript; restriction digested using NotI-HF and AscI enzymes (NEB-R3189 and NEB-R0558) and then ligated into the pre-digested vectors using T4 DNA ligase(M0202) and transformed in *E. coli* DH5-alpha cells. Positive clones were confirmed by insert release and correctness was verified by DNA sequencing.

#### Imaging

Cellular fluorescent images were recorded on a spinning disk confocal microscope at the LMF (Light Microscopy Facility) at the MPI-CBG, with FRAP capability using the 60x/1.2U-Plan-SApo, water immersion objective lens (Olympus) using 405nm, 488nm, 561nm and 640nm laser lines, on an Andor-iXon-897-EMCCD camera. High-throughput images were acquired at the TDS (technology development studio) at the MPI-CBG, cells were imaged in 96 well plates (Greiner Bio-one F-bottom-655090) using 60x objective lens (UPLSAPO60xW, NA = 1.2, WD = 0.28mm) with cmos camera (2560×2160 pixels 16 bit) at binning 1 on the automated Yokogawa CV7000 spinning disk microscope using 405nm, 488nm, 561nm and 640nm laser lines.

### Mesoscale cellular image-analysis methods

#### Manual image analysis

Image analyses, and representative image preparation of the colocalization as well as of the FRAP movies was done prepared using Fiji. The data was plotted using GraphPad prism software. All the images were finally prepared in Adobe Illustrator.

#### Automated image analysis

Condensatogram quantification, multiple calibration curves were generated using purified mEGFP and mCherry proteins utilizing same settings as the one’s used for the imaging of the cells. Fluorescence intensity of the nuclear foci as well as whole nuclear and the dilute phase was measured after segmentation, concentration of the total protein and in the condensed as well as dilute phase were determined by fitting against the generated calibration curve depending on the imaging settings (**Suppl Figure S14**).

One channel contained data on the TRIM protein of interest tagged with GFP or mCherry, while the other contained a nuclear marker protein, either H2B-mCherry or NLS-iRFP. Image analysis was performed using custom scripts written in R, largely using the EBImage library^61^. Nuclear marker fluorescence was used to extract regions of interest (ROIs) corresponding to individual cell nuclei. For this purpose, the nuclear marker channel was mildly blurred using a median-filtering approach (EBImage::medianFilter). Next, a threshold was applied to the image using the EBImage::thresh function, employing a moving rectangular window of 50×50 pixels.

Subsequently, each individual labelled nucleus was virtually cropped out, background pixels set to 0 and the moving rectangle threshold approach (now 10×10 pixels) was applied to the TRIM8 fluorescence intensities. Besides having the entire nucleus labelled (for detection of C_tot_), this generated 2 additional masks: one labeling any present condensates (for detection of C_in_), and one labeling the non-condensate area of the nucleus (for detection of C_out_). For non-condensate area masks a 3-pixel ring surrounding condensates was removed to reduce interface fluorescence effects. The coefficient of variance (CoV) was calculated for the entire nuclear TRIM8 signal. A CoV value bigger than 0.5 was chosen to define a cell as having condensates, while at a CoV ≤ 0.5 a cell was defined as not containing condensates. In the latter case, C_out_ was set to equal C_tot_, and C_in_ and condensate area were set to 0 (µM and %, respectively).

### Cellular scale phenotypic assays

#### Condensation Diagram Analyses

U2OS TRIM8 KO as well as U2OS TRIM33 KO cell lines were generated using CRISPR-Cas9 technology. U2OS TRIM19 Knockout cells were kind gift from Louis Flamand. U2OS TRIM25 Knockout cells kind gift from Paul J Taylor. Cells were transfected with fluorescent tagged TRIM8, TRIM19 and TRIM33 to record the transfected cells with varying level of expression.

#### Generation of stable immortalized podocyte cell lines

Immortalized human podocytes (HSPD) cells with stable integration of pInducer21_Puro_GFP_Hs TRIM8 and _MYC_TRIM8 constructs were generated using a lentiviral system. Lentiviruses were produced by transfecting HEK293T cells with pInducer21_Puro constructs (gift of Dr. Daniela Braun, cloned from pInducer21, Addgene # 87360) with wildtype or patient variant forms of MYC or GFP tagged TRIM8 as well as a packaging plasmids psPAX2 (Addgene #12260) and pMD2.G (Addgene #12259). Supernatants were collected at 24 hours post-transfection, pelleted, and filter sterilized, and were supplemented with 8 µg/mL polybrene (TR-1003-G, Sigma Aldrich). HSPD cells were then cultured with the supernatants for 24 hours and selected with puromycin (A11138-03, Thermo Fisher Scientific).

#### Immunoprecipitation

Cells were lysed using IP lysis buffer (87787, Thermo Fisher Scientific) supplemented with protease inhibitors (78429, Thermo Fisher Scientific) and phosphatase inhibitors (78426, Thermo Fisher Scientific). The lysates were sonicated to ensure thorough disruption and subsequently precleared overnight using Sepharose A beads (PA50-00-0005, Rockland). The precleared lysates were then incubated overnight with conditioned beads conjugated to specific antibodies: MYC (E6654, Sigma Aldrich), FLAG (F2426, Sigma Aldrich), or HA (26181, Thermo Fisher Scientific). Following incubation, the beads were washed thoroughly 10 times to remove nonspecific binding. The bound proteins were then analyzed via immunoblotting.

#### Protein Quantification and Immunoblotting

Following cell lysis, the lysates were sonicated to ensure complete disruption and centrifuged to pellet debris. Protein concentration was determined using the Lowry assay, performed according to the manufacturer’s protocol (Bio-Rad). For electrophoresis, 40–80 µg of protein per sample was loaded into each well of a precast protein gel (Thermo Fisher Scientific). After electrophoresis, proteins were transferred onto membranes using the iBlot system (Thermo Fisher Scientific) for subsequent immunoblotting.

#### Immunofluorescence and confocal imaging

Cells were fixed with 4% paraformaldehyde (PFA) (15710, Electron Microscopy Sciences) for 15 minutes at room temperature and permeabilized using 0.1% Triton X-100 in phosphate-buffered saline (PBS). Following permeabilization, cells were blocked in a solution containing 1% bovine serum albumin (5217100G, Fisher), 10% donkey serum (100-151-500, Gemini Bio-Products), and PBS for 1 hour at room temperature. Blocked cells were incubated with primary antibodies overnight at 4°C. The next day, cells were treated with appropriate secondary antibodies for 1 hour at room temperature. After secondary antibody incubation, cells were counterstained with DAPI (1 mg/mL, Thermo Fisher Scientific) for three 5-minute washes at room temperature. Samples were mounted using ProLong Diamond mountant (Thermo Fisher Scientific), dried overnight, and imaged using a Leica LSM confocal microscope.

#### Proteasome inhibition

pInducer21 puro immortalized cell lines were induced with 1000 ng/mL doxycycline (D9891-5G, Sigma Aldrich) to express MYC-tagged TRIM8 wild-type or patient variant TRIM8. Alternatively, wildtype HSPD cells were transiently transfected with FLAG-tagged TRIM8 wildtype or patient variant TRIM8 constructs. For proteasome inhibition assays, cells expressing MYC- or FLAG-tagged TRIM8 were treated with MG132 (J63250.LB0, Thermo Fisher Scientific), a 26S proteasome inhibitor. Transiently transfected cells were treated with MG132 at concentrations of 0–5 µM for 6 hours, whereas stable pIND21 cell lines expressing MYC-tagged TRIM8 were treated with 0–1 µM MG132 for 6 hours. After treatment, cells were harvested and subjected to immunoblotting to assess MYC- and FLAG-tagged TRIM8 levels. Mouse anti-MYC (2276S, Cell Signaling Technology) and rat anti-FLAG (NBP1-06712, Novus Biologicals) antibodies were used for detection.

#### Cycloheximide chase assay

pInducer21-puro immortalized cell lines were induced with 1000 ng/mL doxycycline to express wild-type TRIM8 or patient variant TRIM8. To assess protein stability, TRIM8-expressing cells were treated with 100 µg/mL cycloheximide (CHX, C1988, Sigma Aldrich) to inhibit protein synthesis. Cells were harvested and lysed at time points ranging from 0 to 5 hours post-treatment. Protein lysates were analyzed by immunoblotting, and TRIM8 levels were detected using a mouse anti-MYC antibody.

#### IP Ubiquitination assay

For ubiquitination studies, HSPD cells were co-transfected with wild-type TRIM8 or patient variant TRIM8 plasmids and HA ubiquitin plasmids using Lipofectamine 2000 transfection reagent (Thermo Fisher Scientific). Transfections were performed for 24 hours. Following transfection, cells were scraped, pelleted, and washed with PBS. The cell pellets were lysed in 200 µL of SDS ubiquitination lysis buffer (2% SDS, 20 mM N-Ethylmaleimide (E3876-5G, Sigma Aldrich), and IP lysis buffer) supplemented with protease and phosphatase inhibitors. The lysates were then boiled for 10 minutes, cooled on ice at 4°C for 20 minutes, and sonicated to ensure complete disruption. Immunoprecipitation was performed as previously described to analyze ubiquitinated proteins.

#### NF-κB reporter assay

Firefly luciferase activities were standardized according to Renilla luciferase activities. 293T cells (5 × 10⁴/well) were seeded into 96-well plates and transfected the following day using Lipofectamine 2000 (Thermo Fisher Scientific). Each well was transfected with 50 ng of either an empty vector, TRIM8 wild-type, or patient variant TRIM8 plasmids, along with 4xNF-κB luciferase reporter plasmid (#111216, Addgene). To standardize transfection efficiency, 5 ng of Renilla luciferase reporter plasmid was co-transfected into each well. Approximately 24 hours post-transfection, luciferase activity was measured using the Dual-Glo® Luciferase Assay System kit (E2940, Promega) according to the manufacturer’s protocol. Firefly luciferase activities were normalized to Renilla luciferase activities to account for variability in transfection efficiency.

#### Proteasome Probing

HSPD cells were transfected with TRIM8 wild-type or \patient variant TRIM8 plasmids and treated with MG132 for 6 hours. After treatment, cells were immunostained with antibodies specific to PSMD4 and PSMD12, followed by imaging as previously described. To observe active proteasomes, TRIM8-overexpressing cells were treated with 2 µM of the activity-based proteasome probe Me4BodipyFL-Ahx3Leu3VS (I-190-050, R&D Systems) for 1 hour at 37°C. Following probe incubation, cells were fixed and imaged to visualize active proteasomes.

#### CRISPR/Cas9 KO cell line generation

HSPD cells were transfected with pSpCas9 plasmids containing guide RNAs (gRNAs) targeting either exon 1 or exon 6 using Lipofectamine 2000 (Thermo Fisher Scientific) for 48 hours. Transfected cells were sorted for eGFP-positive populations via FACS and replated for further culture. After one week of growth, a subset of cells was harvested for genotyping. Genomic DNA was extracted using the Genomic DNA Isolation Kit (K182001, Thermo Fisher Scientific), and the targeted regions were Sanger sequenced to confirm allele loss. Cells were then plated at a low density to facilitate the formation of monoclonal colonies. Individual colonies were isolated using cloning rings and transferred to separate wells for expansion. Once allele loss was confirmed in monoclonal populations, the cells were either cryopreserved or used for subsequent experiments.

#### Characterization of CRISPR/Cas9 KO cell line by PCR

Genomic DNA was isolated using Kit and amplified using Hotstar Taq polymerase (Qiagen). Primers F1_CCAGTCCCACGTGCAGAC and R2_AGAGAAGAGTTTGGCCAGGG were used to detect the deletion in the Exon1. Primers IF2_ACACTCGGTGTGCGACGTG were designed within the deleted region of Exon1 and were used with R2_AGAGAAGAGTTTGGCCAGGG to further confirm the deletion. Primers F3_TCAACGGCCTTCCCAGAG and R3_CTCCCACTGTCACCTCTGC were designed to flank the region of Exon6 KO and were used as a tool to determine the deletion. To further confirm the deletion, we designed the internal primer IF3_CGTCTGTTCTGTGGACAACTGT in the region of deletion and used it with R3_CTCCCACTGTCACCTCTGC to amplify the deleted region. PCR products were confirmed by gel electrophoresis and/or Sanger sequencing.

#### Cell proliferation

For confluency analysis, 5 × 10³ and 1 × 10⁴ cells from Trim8^Ex1-/Ex1-^, Trim8^Ex6-/Ex6-^, and Trim8 wildtype HSPD lines were seeded in 48-well plates. Cells were imaged every 4 hours using light microscopy at 10× magnification with the Incucyte system (Incucyte 2022B Rev2). After 48 hours, the images were analyzed with Incucyte software to measure cell confluency.

For proliferation assays, the MTT assay was performed using the MTT reagent (3-(4,5-dimethylthiazol-2-yl)-2,5-diphenyltetrazolium bromide; Thermo Fisher Scientific) according to the manufacturer’s protocol. Briefly, 5 × 10³ cells were seeded in 96-well plates and cultured at 37°C for 48 hours. MTT reagent was added to each well and incubated for 3 hours at 37°C. The MTT formazan product was solubilized with DMSO, and absorbance was measured at 475 nm and 660 nm at predetermined intervals. Complete media without cells served as a negative control.

Each experiment was conducted in quadruplets with three biological replicates.

#### Cell adhesion assay

Equal numbers of Trim8^Ex1-/Ex1-^, Trim8^Ex6-/Ex6-^, and Trim8 wildtype HSPD cells were labeled with Calcein AM (c1430, Thermo Fisher Scientific) for 20 minutes. A total of 5 × 10³ cells per well were seeded onto 96-well plates, either non-coated or collagen-coated, and allowed to attach for 90 minutes. Pre-wash images were captured using the Incucyte imaging system. To assess cell adhesion, plates were shaken at 1500 rpm for 5 minutes to detach cells, followed by a PBS wash. Post-wash images were then captured. Both pre-wash and post-wash images were exported and analyzed using ImageJ. A custom macro was created to count fluorescent cells in each image. The percentage of cells attached after washing was calculated by normalizing the post-wash cell count to the pre-wash cell count. Data were plotted using Prism 9, and statistical significance between groups was analyzed using the Mann-Whitney test.

#### Cell spreading assay

Trim8^Ex1-/Ex1-^, Trim8^Ex6-/Ex6-^, and Trim8 wildtype HSPD cells were seeded at a density of 5 × 10³ cells per well in 96-well plates and allowed to attach for 60 minutes. After attachment, cells were stained with wheat germ agglutinin (WGA) for 10 minutes to fluorescently label the cell surface.

Images were captured using the Incucyte imaging system, exported, and analyzed using ImageJ. A custom analysis pipeline was created in ImageJ to calculate the fluorescently labeled area of each cell.

### Animal Modeling

#### Mouse Models

Trim8^Ex1m^ and Trim8^Ex6m^ mouse models were generated via CRISPR/CAS9 on the FVB/N background to study the effects of TRIM8 mutations. Mice were genotyped to identify homozygous (Trim8^Ex1m-/Ex1m-^, Trim8^Ex6m-/Ex6m-^), heterozygous (Trim8^Ex1m-/+^, Trim8^Ex6m-/+^), and wildtype (Trim8^+/+^) animals. All experiments were performed on age-matched m mice unless otherwise stated.

#### Periodic Acid–Schiff (PAS) Staining and Glomerular Matrix Deposition Analysis

Kidneys were harvested from 12-month-old mice and fixed in 10% neutral-buffered formalin. Samples were paraffin-embedded, sectioned at 5 µm thickness, and stained with periodic acid–Schiff (PAS) reagent to evaluate glomerular matrix deposition. Representative 20x images of glomeruli from each genotype were captured using a light microscope. An automated ImageJ analysis pipeline was used to quantify matrix deposition in a blinded manner. Statistical significance was determined using the Mann-Whitney U-test.

#### Nephrotoxic Serum (NTS)-Induced Podocyte Injury and Albuminuria Measurement

Sheep nephrotoxic serum (NTS) was used to induce podocyte injury in Trim8Ex6m mice. Mice were injected intravenously with NTS, and urine samples were collected at days 1 to 14 post-treatment to assess albuminuria.

#### Urine protein quantification

Urine was collected by placing mice in collection cages with *ad libitum* access to water overnight (16 h). Upon collection, samples were immediately frozen and stored at - 80 °C and only thawed on ice before urine albumin and creatinine measurements once. Urinary albumin levels were quantified using the Albumin Blue Fluorescent Assay Kit (Active Motif), with standard dilutions prepared from mouse serum albumin (Equitech Bio Inc.). Urine creatinine levels were measured using the QuantiChrom™ Creatinine Assay Kit (BioAssay Systems), according to the manufacturer’s instructions. Albumin-to-creatinine ratios were calculated to normalize urinary albumin levels.

## KEY RESOURCES AND MATERIALS

A comprehensive list of all key resources and materials can be found in Supplementary Document SD1.

## REFERENCES

1. Sabari, B. R., Dall’Agnese, A. & Young, R. A. Biomolecular Condensates in the Nucleus. Trends in Biochemical Sciences 45, 961–977 (2020).

2. Alberti, S. & Hyman, A. A. Biomolecular condensates at the nexus of cellular stress, protein aggregation disease and ageing. Nature Reviews Molecular Cell Biology 22, 196–213 (2021).

3. Wiegand, T. et al. Actin polymerization counteracts prewetting of N-WASP on supported lipid bilayers. Proceedings of the National Academy of Sciences 121, e2407497121 (2024).

4. Zhang, X. et al. Molecular mechanisms of stress-induced reactivation in mumps virus condensates. Cell 186, 1877–1894.e27 (2023).

5. Molliex, A. et al. Phase Separation by Low Complexity Domains Promotes Stress Granule Assembly and Drives Pathological Fibrillization. Cell 163, 123–133 (2015).

6. Patel, A. et al. A Liquid-to-Solid Phase Transition of the ALS Protein FUS Accelerated by Disease Mutation. Cell 162, 1066–1077 (2015).

7. Klein, I. A. et al. Partitioning of cancer therapeutics in nuclear condensates. Science 368, 1386–1392 (2020).

8. Kiss, L. et al. Trim-Away ubiquitinates and degrades lysine-less and N-terminally acetylated substrates. Nature Communications 14, 2160 (2023).

9. Reymond, A. et al. The tripartite motif family identifies cell compartments. EMBO J 20, 2140 (2001).

10. Esposito, D., Koliopoulos, M. G. & Rittinger, K. Structural determinants of TRIM protein function. Biochemical Society Transactions 45, 183–191 (2017).

11. Vunjak, M. & Versteeg, G. A. TRIM proteins. Current Biology 29, R42–R44 (2019).

12. Ozato, K., Shin, D.-M., Chang, T.-H. & Morse, H. C. TRIM family proteins and their emerging roles in innate immunity. Nature Reviews Immunology 8, 849–860 (2008).

13. Mascle, X. H. et al. Acetylation of SUMO1 Alters Interactions with the SIMs of PML and Daxx in a Protein-Specific Manner. Structure 28, 157–168.e5 (2020).

14. Shang, Z. et al. TRIM25 predominately associates with anti-viral stress granules. Nature Communications 15, 4127 (2024).

15. Tozawa, T. et al. Ubiquitination-coupled liquid phase separation regulates the accumulation of the TRIM family of ubiquitin ligases into cytoplasmic bodies. PLOS ONE 17, e0272700 (2022).

16. Lallemand-Breitenbach, V. & de Thé, H. PML Nuclear Bodies. Cold Spring Harbor Perspectives in Biology 2, a000661 (2010).

17. Kimura, T., Mandell, M. & Deretic, V. Precision autophagy directed by receptor regulators – emerging examples within the TRIM family. Journal of Cell Science 129, 881–891 (2016).

18. Borlepawar, A., Frey, N. & Rangrez, A. Y. A systematic view on E3 ligase Ring TRIMmers with a focus on cardiac function and disease. Trends in Cardiovascular Medicine 29, 1–8 (2019).

19. Hatakeyama, S. TRIM Family Proteins: Roles in Autophagy, Immunity, and Carcinogenesis. Trends in Biochemical Sciences 42, 297–311 (2017).

20. Van Tol, S., Hage, A., Giraldo, M. I., Bharaj, P. & Rajsbaum, R. The TRIMendous Role of TRIMs in Virus–Host Interactions. Vaccines 5, (2017).

21. Assoum, M. et al. Further delineation of the clinical spectrum of de novo TRIM8 truncating mutations. American Journal of Medical Genetics Part A 176, 2470–2478 (2018).

22. McClatchey, M. A. et al. Focal segmental glomerulosclerosis and mild intellectual disability in a patient with a novel de novo truncating TRIM8 mutation. European Journal of Medical Genetics 103972 (2020) doi:10.1016/j.ejmg.2020.103972.

23. Weng, P. L. et al. De novo TRIM8 variants impair its protein localization to nuclear bodies and cause developmental delay, epilepsy, and focal segmental glomerulosclerosis. The American Journal of Human Genetics 108, 357–367 (2021).

24. Sakai, Y. et al. De Novo Truncating Mutation of TRIM8 Causes Early-Onset Epileptic Encephalopathy. Annals of Human Genetics 80, 235–240 (2016).

25. Warren, M. et al. Association of a de novo nonsense mutation of the TRIM8 gene with childhood-onset focal segmental glomerulosclerosis. Pediatric Nephrology (2020) doi:10.1007/s00467-020-04525-3.

26. Li, W. & Guo, H. De novo truncating variants of TRIM8 and atypical neuro-renal syndrome: a case report and literature review. Italian Journal of Pediatrics 49, 46 (2023).

27. Badeńska, M. et al. A Rare De Novo Mutation in the TRIM8 Gene in a 17-Year-Old Boy with Steroid-Resistant Nephrotic Syndrome: Case Report. International Journal of Molecular Sciences 25, (2024).

28. Mann, N., Sun, H. & Majmundar, A. J. Mechanisms of podocyte injury in genetic kidney disease. Pediatric Nephrology (2024) doi:10.1007/s00467-024-06551-x.

29. Peng, K., Radivojac, P., Vucetic, S., Dunker, A. K. & Obradovic, Z. Length-dependent prediction of protein intrinsic disorder. BMC Bioinformatics 7, 208 (2006).

30. Uversky, V. N. Unreported intrinsic disorder in proteins: Building connections to the literature on IDPs. Intrinsically Disordered Proteins 2, e970499 (2014).

31. Hardenberg, M., Horvath, A., Ambrus, V., Fuxreiter, M. & Vendruscolo, M. Widespread occurrence of the droplet state of proteins in the human proteome. Proceedings of the National Academy of Sciences 117, 33254–33262 (2020).

32. Hadarovich, A. et al. PICNIC accurately predicts condensate-forming proteins regardless of their structural disorder across organisms. Nature Communications 15, 10668 (2024).

33. Rold, C. J. & Aiken, C. Proteasomal Degradation of TRIM5α during Retrovirus Restriction. PLOS Pathogens 4, e1000074 (2008).

34. Clift, D. et al. A Method for the Acute and Rapid Degradation of Endogenous Proteins. Cell 171, 1692–1706.e18 (2017).

35. Iriki, T. et al. Senescent cells form nuclear foci that contain the 26S proteasome. Cell Reports 42, 112880 (2023).

36. de Jong, A., Schuurman, K. G., Rodenko, B., Ovaa, H. & Berkers, C. R. Fluorescence-Based Proteasome Activity Profiling. in Chemical Proteomics: Methods and Protocols (eds. Drewes, G. & Bantscheff, M.) 183–204 (Humana Press, Totowa, NJ, 2012). doi:10.1007/978-1-61779-364-6_13.

37. Li, Q. et al. Tripartite motif 8 (TRIM8) modulates TNFα- and IL-1β–triggered NF-κB activation by targeting TAK1 for K63-linked polyubiquitination. Proc Natl Acad Sci USA 108, 19341 (2011).

38. Chen, L. et al. Tripartite Motif 8 Contributes to Pathological Cardiac Hypertrophy Through Enhancing Transforming Growth Factor β–Activated Kinase 1–Dependent Signaling Pathways. Hypertension 69, 249–258 (2017).

39. Tomar, D. et al. Nucleo-Cytoplasmic Trafficking of TRIM8, a Novel Oncogene, Is Involved in Positive Regulation of TNF Induced NF-κB Pathway. PLOS ONE 7, e48662 (2012).

40. Tao, Q. et al. Tripartite Motif 8 Deficiency Relieves Hepatic Ischaemia/reperfusion Injury via TAK1-dependent Signalling Pathways. Int J Biol Sci 15, 1618–1629 (2019).

41. Yan, F. et al. The E3 ligase tripartite motif 8 targets TAK1 to promote insulin resistance and steatohepatitis. Hepatology 65, (2017).

42. Liu, R., Wu, H. & Song, H. Knockdown of TRIM8 Attenuates IL-1β-induced Inflammatory Response in Osteoarthritis Chondrocytes Through the Inactivation of NF-κB Pathway. Cell Transplant 29, 0963689720943604 (2020).

43. Bai, X., Zhang, Y.-L. & Liu, L.-N. Inhibition of TRIM8 restrains ischaemia-reperfusion-mediated cerebral injury by regulation of NF-κB activation associated inflammation and apoptosis. Experimental Cell Research 388, 111818 (2020).

44. Fearn, A. et al. The NF-κB1 is a key regulator of acute but not chronic renal injury. Cell Death & Disease 8, e2883–e2883 (2017).

45. Kim, S. I. et al. TGF-β–Activated Kinase 1 Is Crucial in Podocyte Differentiation and Glomerular Capillary Formation. Journal of the American Society of Nephrology 25, (2014).

46. Lennon, R., Randles, M. J. & Humphries, M. J. The Importance of Podocyte Adhesion for a Healthy Glomerulus. Frontiers in Endocrinology 5, (2014).

47. Sachs, N. & Sonnenberg, A. Cell–matrix adhesion of podocytes in physiology and disease. Nature Reviews Nephrology 9, 200–210 (2013).

48. George, B. et al. Crk1/2-dependent signaling is necessary for podocyte foot process spreading in mouse models of glomerular disease. J Clin Invest 122, 674–692 (2012).

49. Verma, R. et al. Nephrin is necessary for podocyte recovery following injury in an adult mature glomerulus. PLOS ONE 13, e0198013 (2018).

50. Topham, P. S. et al. Lack of chemokine receptor CCR1 enhances Th1 responses and glomerular injury during nephrotoxic nephritis. J Clin Invest 104, 1549–1557 (1999).

51. Yanagita, M. et al. Essential role of Gas6 for glomerular injury in nephrotoxic nephritis. J Clin Invest 110, 239–246 (2002).

52. Chugh, S. et al. Aminopeptidase A: A nephritogenic target antigen of nephrotoxic serum. Kidney International 59, 601–613 (2001).

53. Lander, E. S. et al. Initial sequencing and analysis of the human genome. Nature 409, 860– 921 (2001).

54. The UniProt Consortium. The Universal Protein Resource (UniProt). Nucleic Acids Research 35, D193–D197 (2007).

55. Ma, Z. et al. PTC-bearing mRNA elicits a genetic compensation response via Upf3a and COMPASS components. Nature 568, 259–263 (2019).

56. Butt, L. et al. Super-Resolution Imaging of the Filtration Barrier Suggests a Role for Podocin R229Q in Genetic Predisposition to Glomerular Disease. Journal of the American Society of Nephrology 33, (2022).

57. Subramanian, B. et al. Mice with mutant Inf2 show impaired podocyte and slit diaphragm integrity in response to protamine-induced kidney injury. Kidney International 90, 363–372 (2016).

58. Jumper, J. et al. Highly accurate protein structure prediction with AlphaFold. Nature 596, 583– 589 (2021).

59. Varadi, M. et al. AlphaFold Protein Structure Database: massively expanding the structural coverage of protein-sequence space with high-accuracy models. Nucleic Acids Research 50, D439–D444 (2022).

60. Abramson, J. et al. Accurate structure prediction of biomolecular interactions with AlphaFold 3. Nature 630, 493–500 (2024).

61. Pau, G., Fuchs, F., Sklyar, O., Boutros, M. & Huber, W. EBImage—an R package for image processing with applications to cellular phenotypes. Bioinformatics 26, 979–981 (2010).

