## Supplementary Appendix and Figures for "Mesoscale landscaping of the TRIM protein family reveals a novel human condensatopathy"

### Supplementary Figures

**Supplementary Figure S1. Alpha-fold models for TRIM family proteins.** Images of Alpha-fold derived structures are shown corresponding to structure files (**see Suppl Files SF2-SF72**).

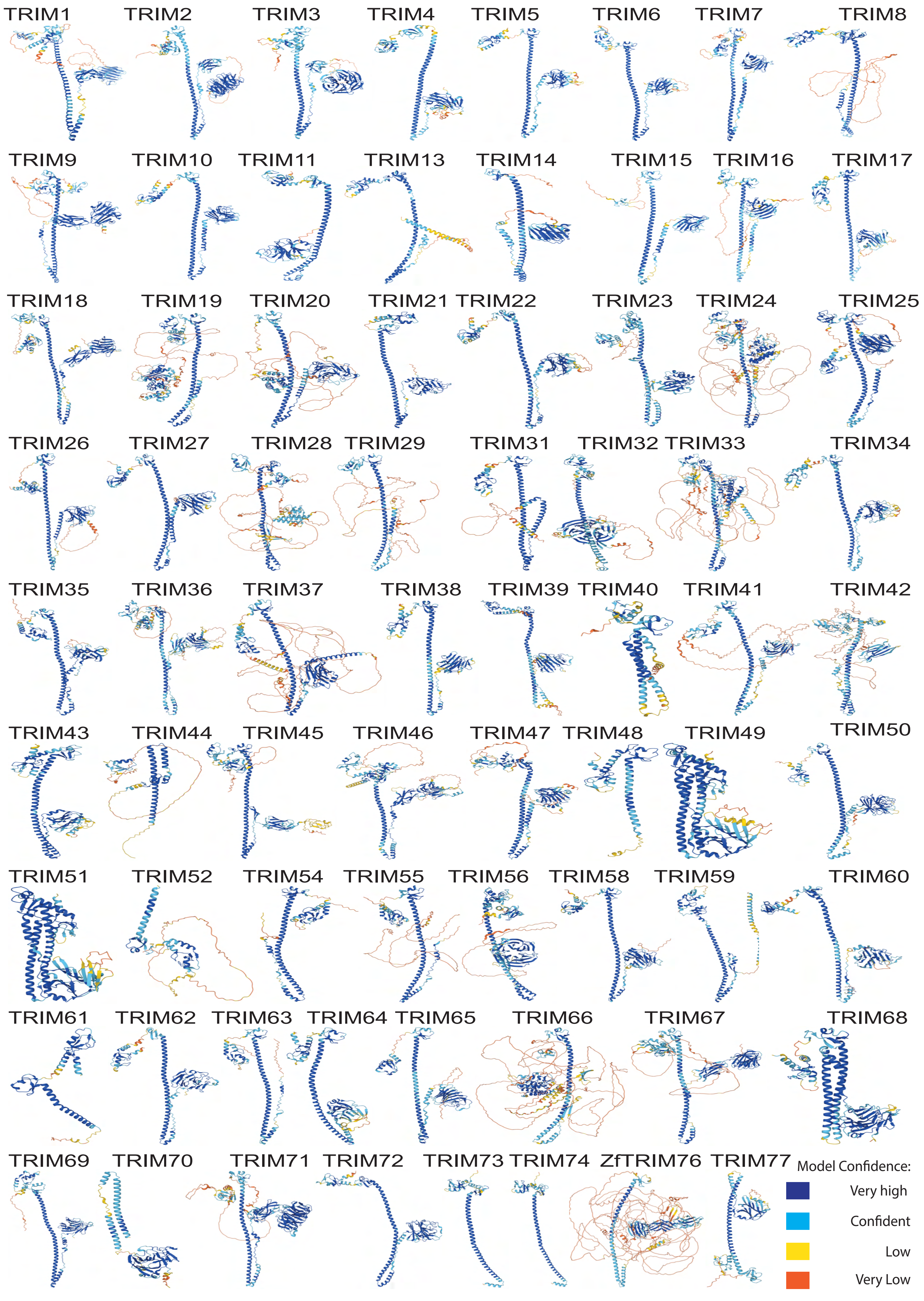

TRIM1
TRIM2
TRIM3
TRIM4
TRIM5
TRIM6
TRIM7
TRIM8
TRIM9
TRIM10
TRIM11
TRIM13
TRIM14
TRIM15
TRIM16
TRIM17
TRIM18
TRIM19
TRIM20
TRIM21
TRIM22
TRIM23
TRIM24
TRIM25
TRIM26
TRIM27
TRIM28
TRIM29
TRIM31
TRIM32
TRIM33
TRIM34
TRIM35
TRIM36
TRIM37
TRIM38
TRIM39
TRIM40
TRIM41
TRIM42
TRIM43
TRIM44
TRIM45
TRIM46
TRIM47
TRIM48
TRIM49
TRIM50
TRIM51
TRIM52
TRIM54
TRIM55
TRIM56
TRIM58
TRIM59
TRIM60
TRIM61
TRIM62
TRIM63
TRIM64
TRIM65
TRIM66
TRIM67
TRIM68
TRIM69
TRIM70
TRIM71
TRIM72
TRIM73
TRIM74
ZfTRIM76
TRIM77
Model Confidence:
Very high
Confident
Low
Very Low

**Supplementary Figure S2. Overview of the TRIM family proteins that display a variety of specific mesoscale organizations in U2OS cells.**

**A.** Representative examples of the TRIM family proteins that predominantly localize to the nucleus in bodies or diffuse patterns in U2OS cells are shown. (Note that while localization patterns were generally concordant, TRIM52 in HELA cells show diffused localization, while in U2OS cells it shows nuclear bodies.). See further analysis of these proteins in **Figure 2E**.

**B.** Representative examples are shown of the TRIM family proteins that localize to the cytoplasm, including cytoplasmic filaments and the cytoplasmic bodies, in U2OS cells. (For HELA cells, **see Figure 1.**)

**C.** Representative examples are shown of the TRIM family proteins that localize to the membranous compartments in U2OS cells.

**D.** A visual summary of the consensus global cellular mesoscale localization patterns that were observed in HELA and/or U2OS cells (**see Suppl Figure S2**) for the TRIM family.

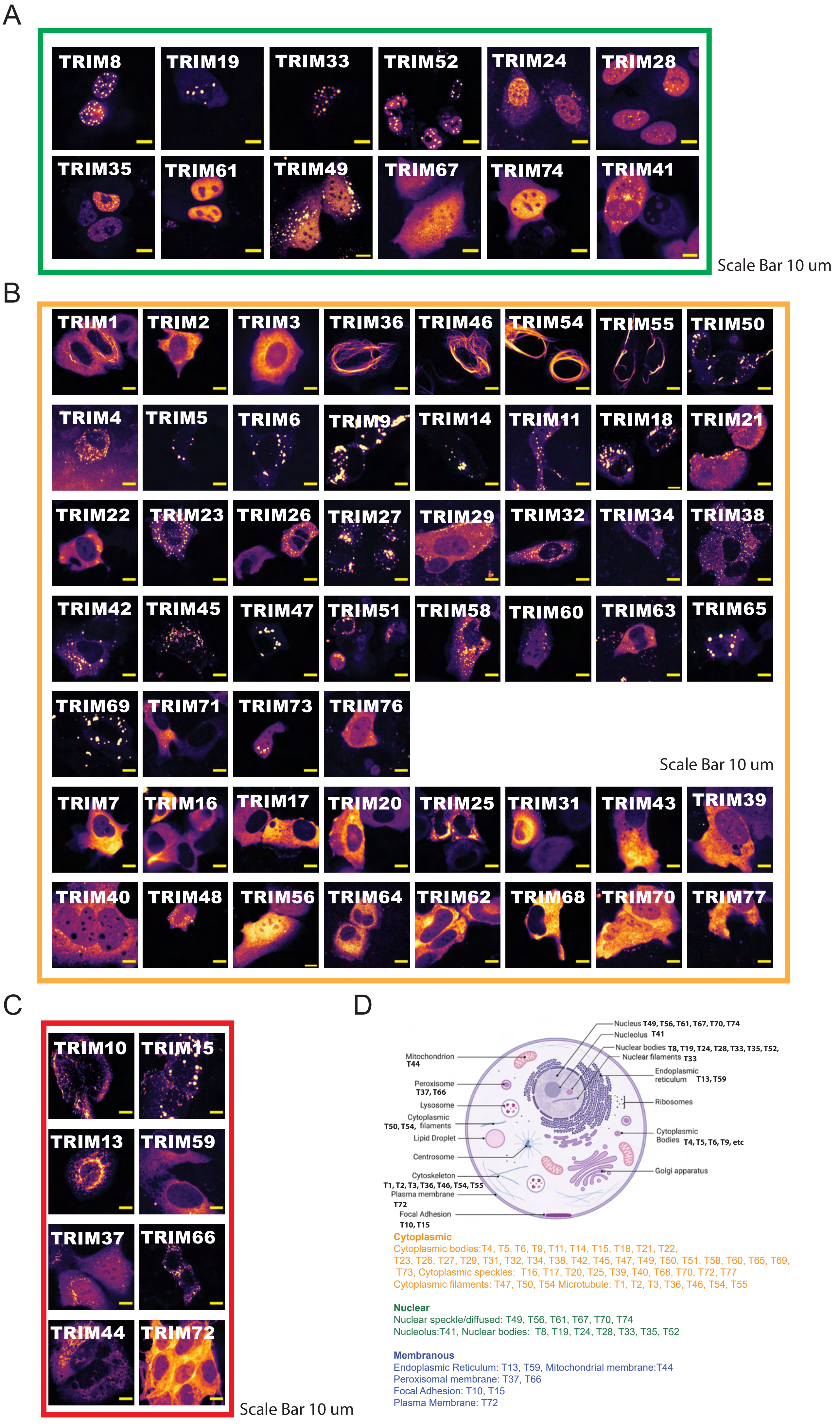
A
TRIM8
TRIM19
TRIM33
TRIM52
TRIM24
TRIM28
TRIM35
TRIM61
TRIM49
TRIM67
TRIM74
TRIM41
Scale Bar 10 um
B
TRIM1
TRIM2
TRIM3
TRIM36
TRIM46
TRIM54
TRIM55
TRIM50
TRIM4
TRIM5
TRIM6
TRIM9
TRIM14
TRIM11
TRIM18
TRIM21
TRIM22
TRIM23
TRIM26
TRIM27
TRIM29
TRIM32
TRIM34
TRIM38
TRIM42
TRIM45
TRIM47
TRIM51
TRIM58
TRIM60
TRIM63
TRIM65
TRIM69
TRIM71
TRIM73
TRIM76
Scale Bar 10 um
TRIM7
TRIM16
TRIM17
TRIM20
TRIM25
TRIM31
TRIM43
TRIM39
TRIM40
TRIM48
TRIM56
TRIM64
TRIM62
TRIM68
TRIM70
TRIM77
C
TRIM10
TRIM15
TRIM13
TRIM59
TRIM37
TRIM66
TRIM44
TRIM72
Scale Bar 10 um
D
Nucleus T49, T56, T61, T67, T70, T74
Nucleolus T41
Nuclear bodies T8, T19, T24, T28, T33, T35, T52,
Nuclear filaments T33
Endoplasmic reticulum T13, T59
Ribosomes
Cytoplasmic Bodies T4, T5, T6, T9, etc
Golgi apparatus
Mitochondrion T44
Peroxisome T37, T66
Lysosome
T50, T54, Cytoplasmic filaments
Lipid Droplet
Centrosome
Cytoskeleton T1, T2, T3, T36, T46, T54, T55
Plasma membrane T72
Focal Adhesion T10, T15
Cytoplasmic
Cytoplasmic bodies:T4, T5, T6, T9, T11, T14, T15, T18, T21, T22, T23, T26, T27, T29, T31, T32, T34, T38, T42, T45, T47, T49, T50, T51, T58, T60, T65, T69, T73, Cytoplasmic speckles: T16, T17, T20, T25, T39, T40, T68, T70, T72, T77
Cytoplasmic filaments: T47, T50, T54 Microtubule: T1, T2, T3, T36, T46, T54, T55
Nuclear
Nuclear speckle/diffused: T49, T56, T61, T67, T70, T74
Nucleolus:T41, Nuclear bodies: T8, T19, T24, T28, T33, T35, T52
Membranous
Endoplasmic Reticulum: T13, T59, Mitochondrial membrane:T44
Peroxisomal membrane: T37, T66
Focal Adhesion: T10, T15
Plasma Membrane: T72

**Supplementary Figure S3. Domain structure and dynamic recovery for TRIM1, TRIM5, and TRIM6.**

**A.** Domain structure of TRIM1 shown.

**B.** Fluorescence recovery after photo-bleaching curve shown for TRIM1 filaments.

**C.** Representative images of fluorescence recovery after photo-bleaching shown for TRIM1 filaments.

**D.** Alpha-fold structure for TRIM1 shown.

**E.** Domain structure of TRIM5 shown.

**F.** Fluorescence recovery after photo-bleaching curve shown for TRIM5.

**G.** Representative images of fluorescence recovery after photo-bleaching shown for TRIM5 filaments.

**H.** Alpha-fold structure for TRIM5 shown.

**I.** Domain structure of TRIM6 shown.

**J.** Fluorescence recovery after photo-bleaching curve shown for TRIM6.

**K.** Representative images of fluorescence recovery after photo-bleaching shown for TRIM6 filaments.

**L.** Alpha-fold structure for TRIM6 shown.

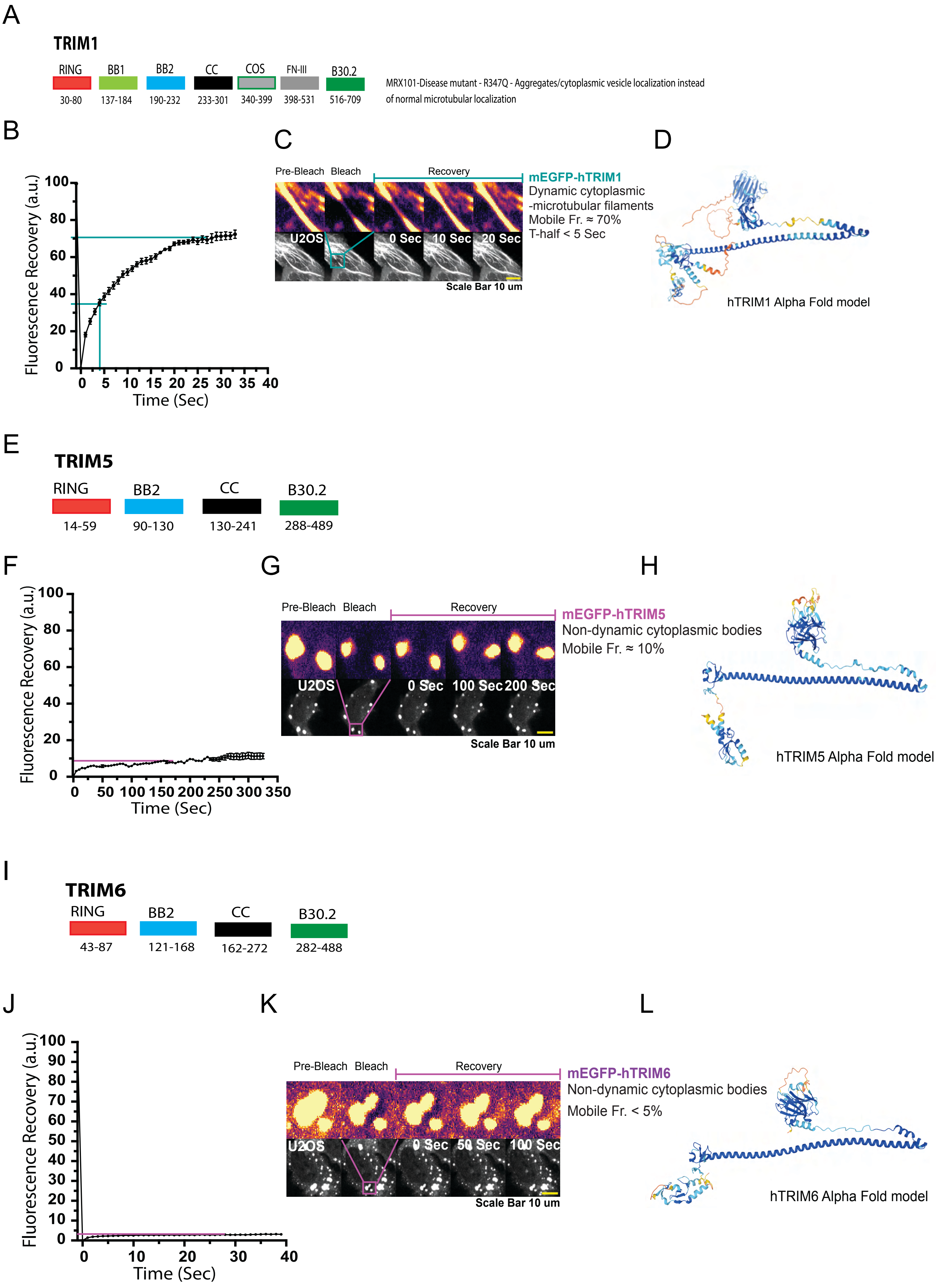

A
TRIM1
RING
BB1
BB2
CC
COS
FN-III
B30.2
30-80
137-184
190-232
233-301
340-399
398-531
516-709
MRX101-Disease mutant - R347Q - Aggregates/cytoplasmic vesicle localization instead of normal microtubular localization
B
Fluorescence Recovery (a.u.)
Time (Sec)
C
Pre-Bleach
Bleach
Recovery
U2OS
0 Sec
10 Sec
20 Sec
Scale Bar 10 um
mEGFP-hTRIM1
Dynamic cytoplasmic -microtubular filaments
Mobile Fr. ≈ 70%
T-half < 5 Sec
D
hTRIM1 Alpha Fold model
E
TRIM5
RING
BB2
CC
B30.2
14-59
90-130
130-241
288-489
F
Fluorescence Recovery (a.u.)
Time (Sec)
G
Pre-Bleach
Bleach
Recovery
U2OS
0 Sec
100 Sec
200 Sec
Scale Bar 10 um
mEGFP-hTRIM5
Non-dynamic cytoplasmic bodies
Mobile Fr. ≈ 10%
H
hTRIM5 Alpha Fold model
I
TRIM6
RING
BB2
CC
B30.2
43-87
121-168
162-272
282-488
J
Fluorescence Recovery (a.u.)
Time (Sec)
K
Pre-Bleach
Bleach
Recovery
U2OS
0 Sec
50 Sec
100 Sec
Scale Bar 10 um
mEGFP-hTRIM6
Non-dynamic cytoplasmic bodies
Mobile Fr. < 5%
L
hTRIM6 Alpha Fold model

**Supplementary Figure S4. Dynamic recovery for TRIM7.**

**A.** Domain structure of TRIM7 shown.

**B.** Representative images of fluorescence recovery after photo-bleaching shown for TRIM7.

**C.** Fluorescence recovery after photo-bleaching curve shown for TRIM7.

A

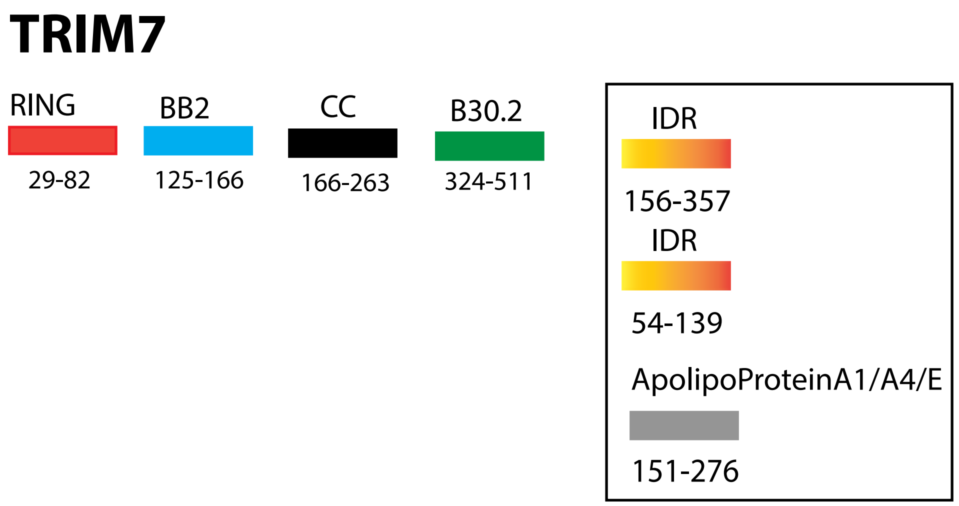

TRIM7
RING
29-82
BB2
125-166
CC
166-263
B30.2
324-511
IDR
156-357
IDR
54-139
ApolipoProteinA1/A4/E
151-276


B

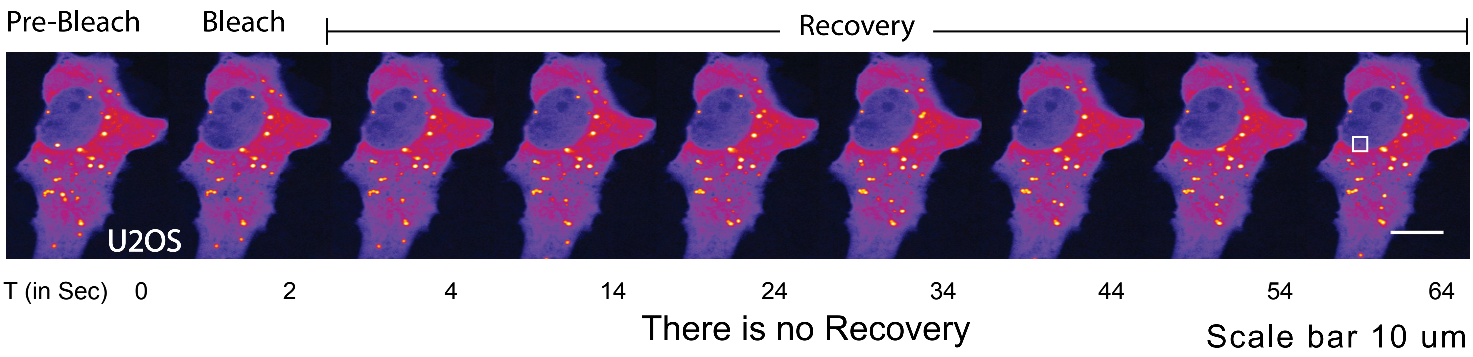

Pre-Bleach
Bleach
Recovery
U2OS
T (in Sec) 0 2 4 14 24 34 44 54 64
There is no Recovery
Scale bar 10 um


C

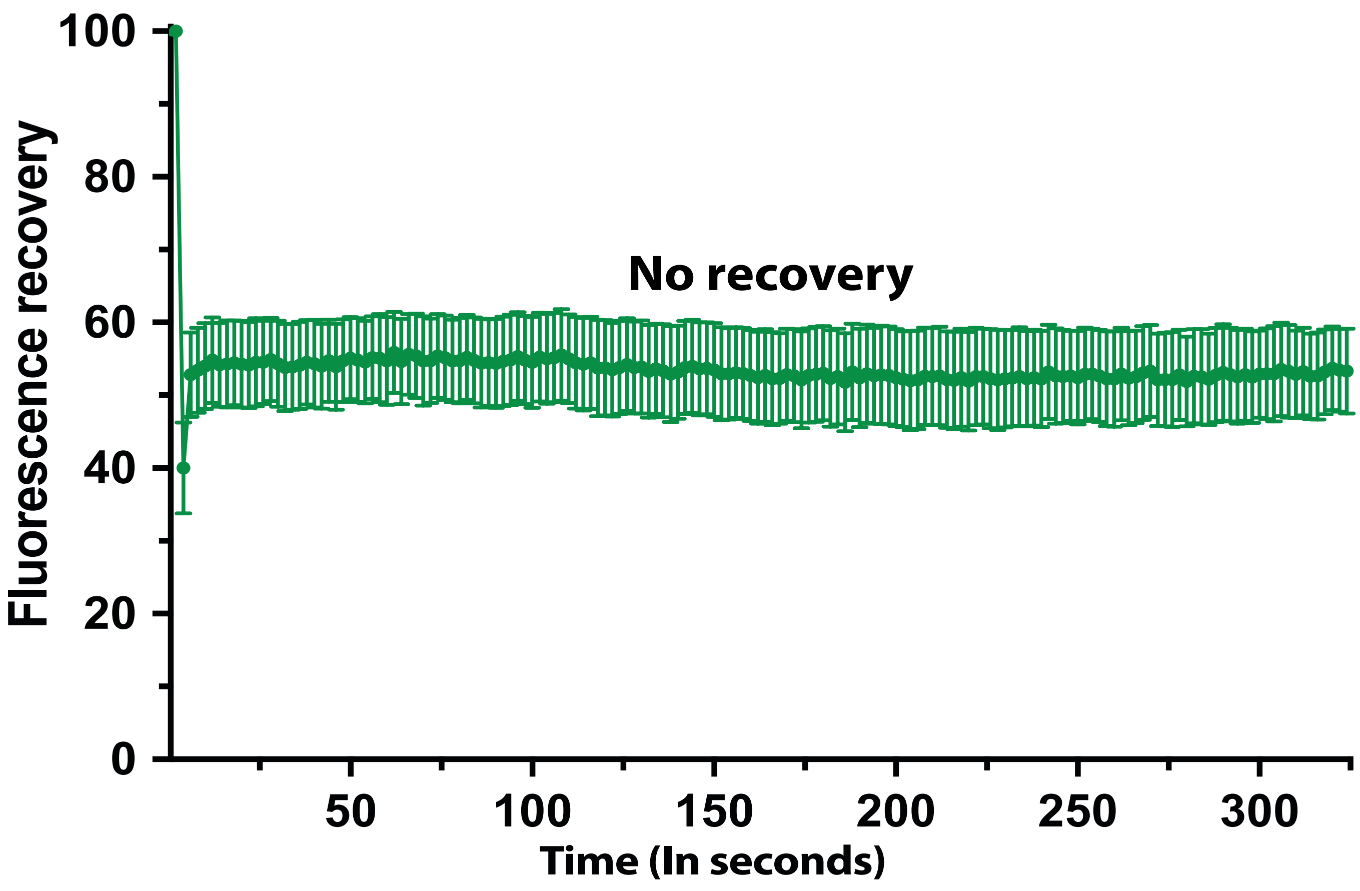

Fluorescence recovery
100
80
60
40
20
0
No recovery
50
100
150
200
250
300
Time (In seconds)

**Supplementary Figure S5. Dynamic recovery for TRIM8 and TRIM9.**

**A.** Domain structure of TRIM8 shown.

**B.** Representative images of fluorescence recovery after photo-bleaching shown for TRIM8.

**C.** Domain structure of TRIM9 shown.

**D.** Fluorescence recovery after photo-bleaching curve shown for TRIM9.

**E.** Representative images of fluorescence recovery after photo-bleaching shown for TRIM9.

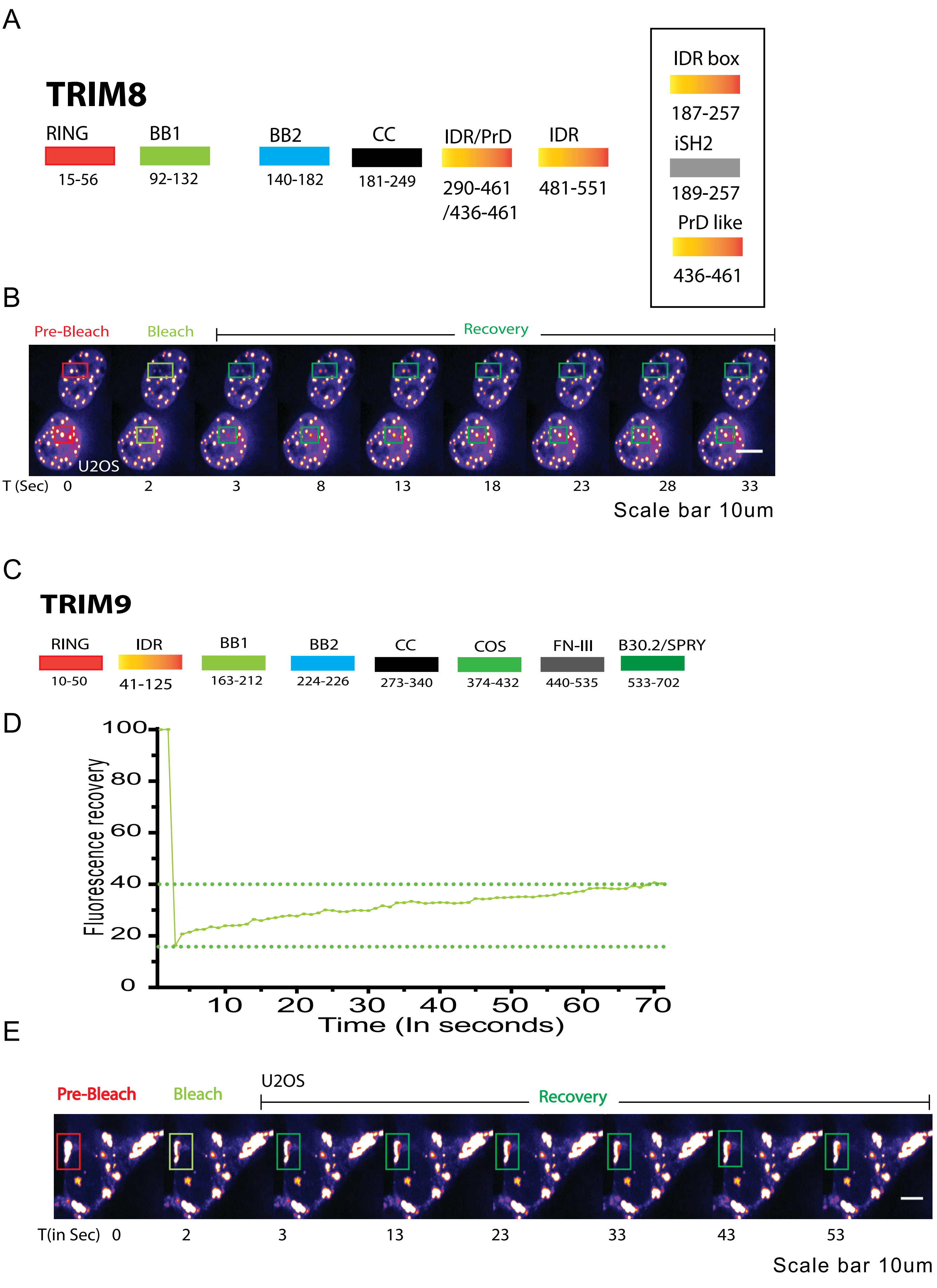

A
TRIM8
RING
15-56
BB1
92-132
BB2
140-182
CC
181-249
IDR/PrD
290-461
/436-461
IDR
481-551
IDR box
187-257
iSH2
189-257
PrD like
436-461
B
Pre-Bleach
Bleach
Recovery
U2OS
T (Sec) 0 2 3 8 13 18 23 28 33
Scale bar 10um
C
TRIM9
RING
10-50
IDR
41-125
BB1
163-212
BB2
224-226
CC
273-340
COS
374-432
FN-III
440-535
B30.2/SPRY
533-702
D
Fluorescence recovery
0
20
40
60
80
100
10 20 30 40 50 60 70
Time (In seconds)
E
Pre-Bleach
Bleach
U2OS
Recovery
T(in Sec) 0 2 3 13 23 33 43 53
Scale bar 10um

**Supplementary Figure S6. Dynamic recovery for TRIM10 and TRIM11.**

**A.** Domain structure of TRIM10 shown.

**B.** Representative images of fluorescence recovery after photo-bleaching shown for TRIM10.

**C.** Domain structure of TRIM11 shown.

**D.** Fluorescence recovery after photo-bleaching curve shown for TRIM11.

**E.** Representative images of fluorescence recovery after photo-bleaching shown for TRIM11.

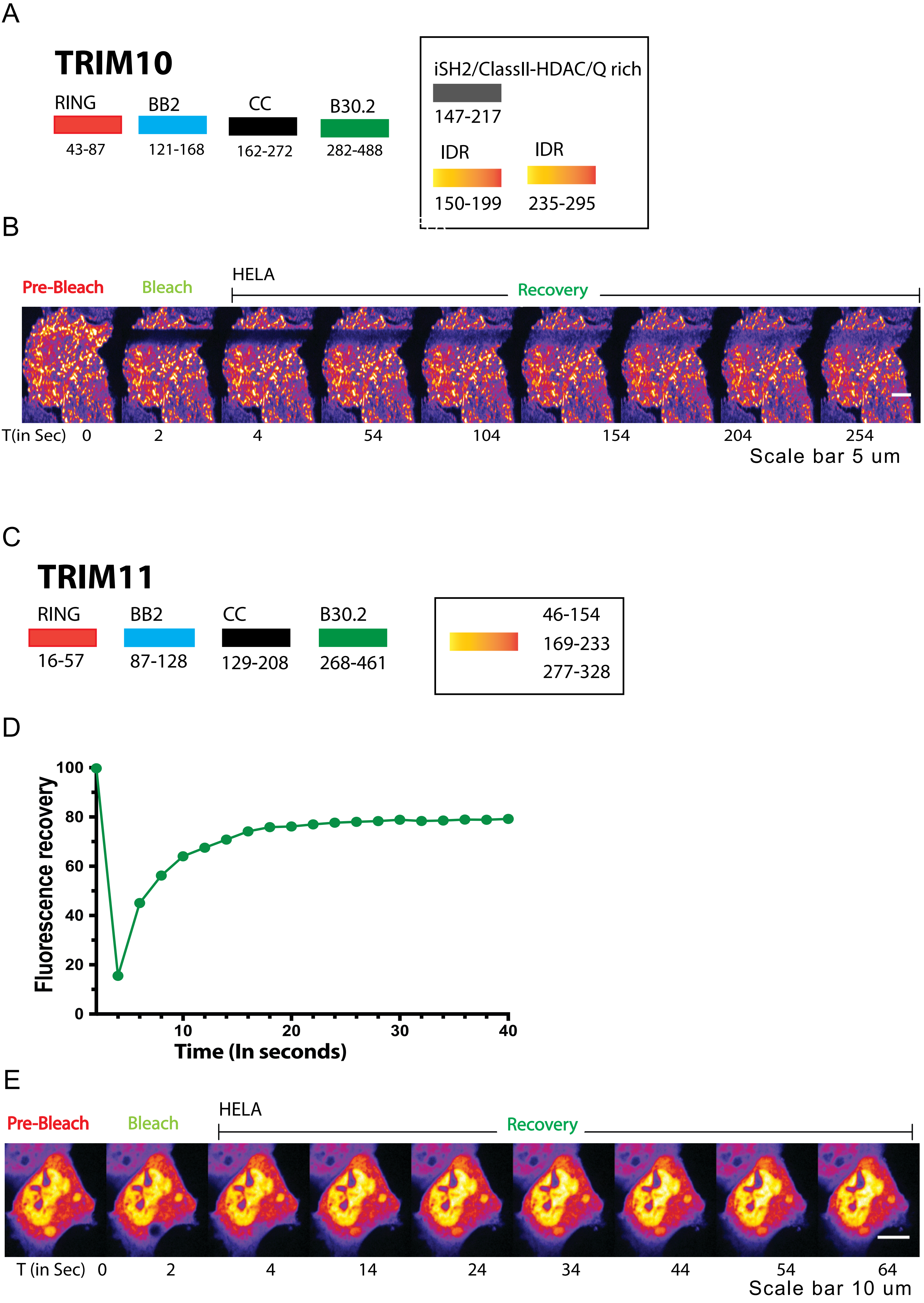
A
TRIM10
RING
43-87
BB2
121-168
CC
162-272
B30.2
282-488
iSH2/ClassII-HDAC/Q rich
147-217
IDR
150-199
IDR
235-295
B
Pre-Bleach
Bleach
HELA
Recovery
T(in Sec) 0 2 4 54 104 154 204 254
Scale bar 5 um
C
TRIM11
RING
16-57
BB2
87-128
CC
129-208
B30.2
268-461
46-154
169-233
277-328
D
Fluorescence recovery
100
80
60
40
20
0
10
20
30
40
Time (In seconds)
E
Pre-Bleach
Bleach
HELA
Recovery
T (in Sec) 0 2 4 14 24 34 44 54 64
Scale bar 10 um

### Supplementary Figure S7. Dynamic recovery for TRIM13 and TRIM16.

**A.** Domain structure of TRIM13 shown.

**B.** Representative images of fluorescence recovery after photo-bleaching shown for TRIM13.

**C.** Representative images of fluorescence recovery after photo-bleaching shown for TRIM16.

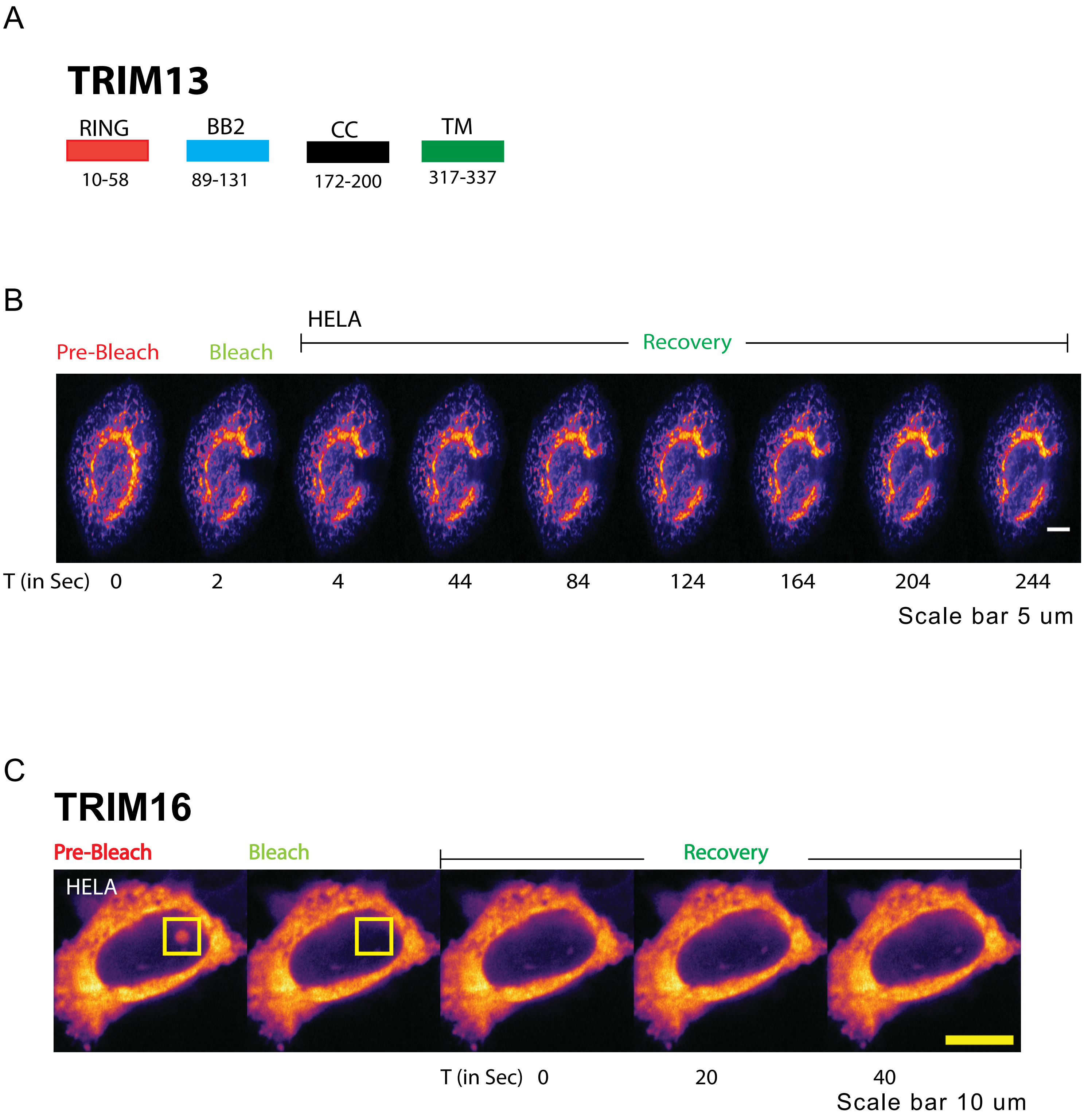
A
TRIM13
RING
10-58
BB2
89-131
CC
172-200
TM
317-337
B
HELA
Pre-Bleach
Bleach
Recovery
T (in Sec) 0 2 4 44 84 124 164 204 244
Scale bar 5 um
C
TRIM16
Pre-Bleach
Bleach
Recovery
HELA
T (in Sec) 0 20 40
Scale bar 10 um

### Supplementary Figure S8. Dynamic recovery for TRIM15.

**A.** Domain structure of TRIM15 shown.

**B.** Representative fluorescent images of TRIM15 expressed in wildtype and TRIM25 KO U2OS cells.

**C.** Representative images of fluorescence recovery after photo-bleaching shown for TRIM15 in wildtype U2OS cells.

**D.** Representative images of fluorescence recovery after photo-bleaching shown for TRIM15 in TRIM25 KO U2OS cells.

**E.** Curve of fluorescence recovery after photo-bleaching shown for TRIM15 in TRIM25 KO U2OS cells.

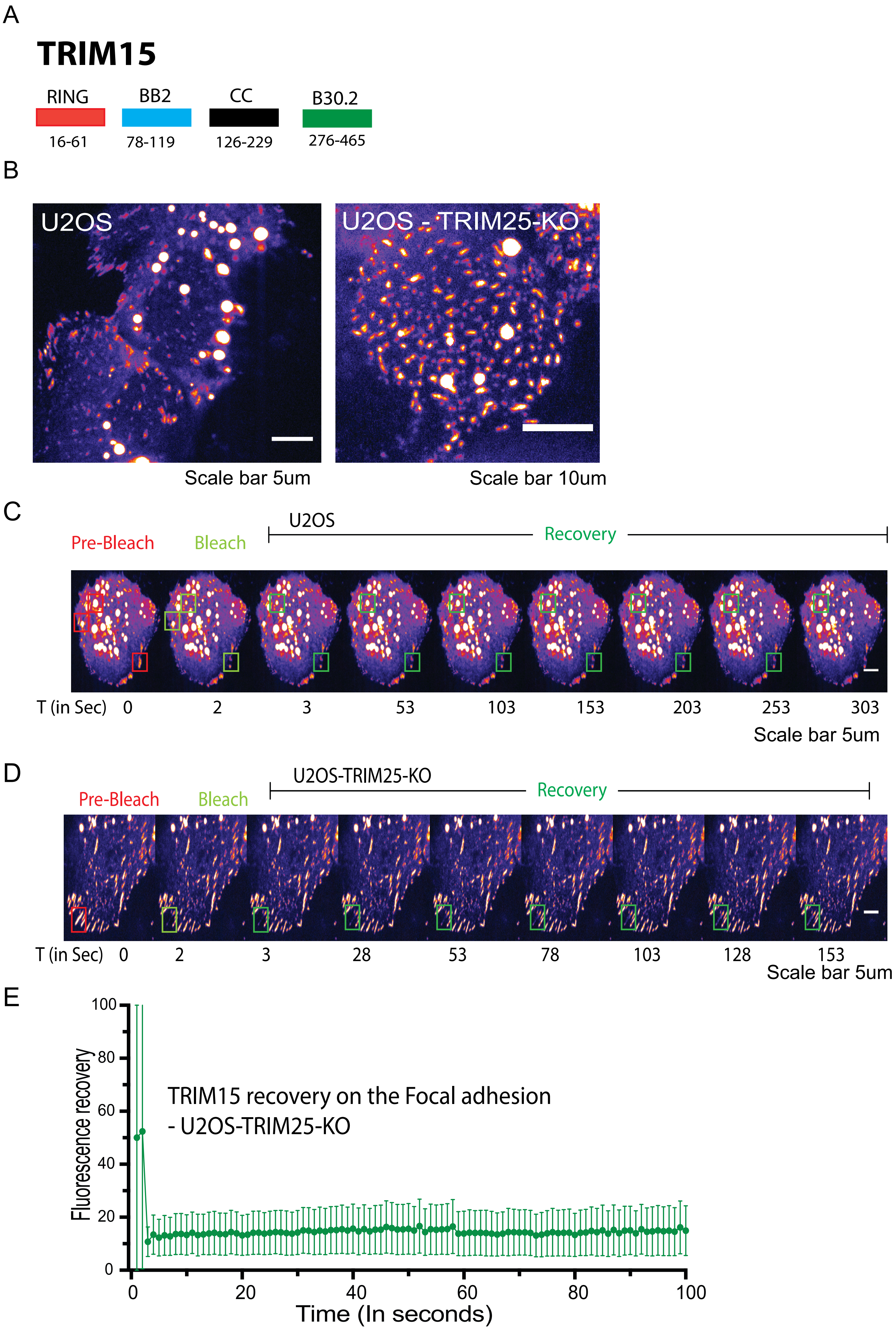
A
TRIM15
RING
BB2
CC
B30.2
16-61
78-119
126-229
276-465
B
U2OS
U2OS - TRIM25-KO
Scale bar 5um
Scale bar 10um
C
U2OS
Pre-Bleach
Bleach
Recovery
T (in Sec) 0 2 3 53 103 153 203 253 303
Scale bar 5um
D
U2OS-TRIM25-KO
Pre-Bleach
Bleach
Recovery
T (in Sec) 0 2 3 28 53 78 103 128 153
Scale bar 5um
E
Fluorescence recovery
TRIM15 recovery on the Focal adhesion
- U2OS-TRIM25-KO
Time (In seconds)
0
20
40
60
80
100

### Supplementary Figure S9. Dynamic recovery for TRIM33.

**A.** Domain structure of TRIM33 shown.

**B.** Representative fluorescent images of TRIM33 localization in U2OS, A549 and HCT-116 cells.

**C.** Fluorescence recovery after photo-bleaching curve shown for TRIM33.

**D.** Representative images of fluorescence recovery after photo-bleaching shown for TRIM33.

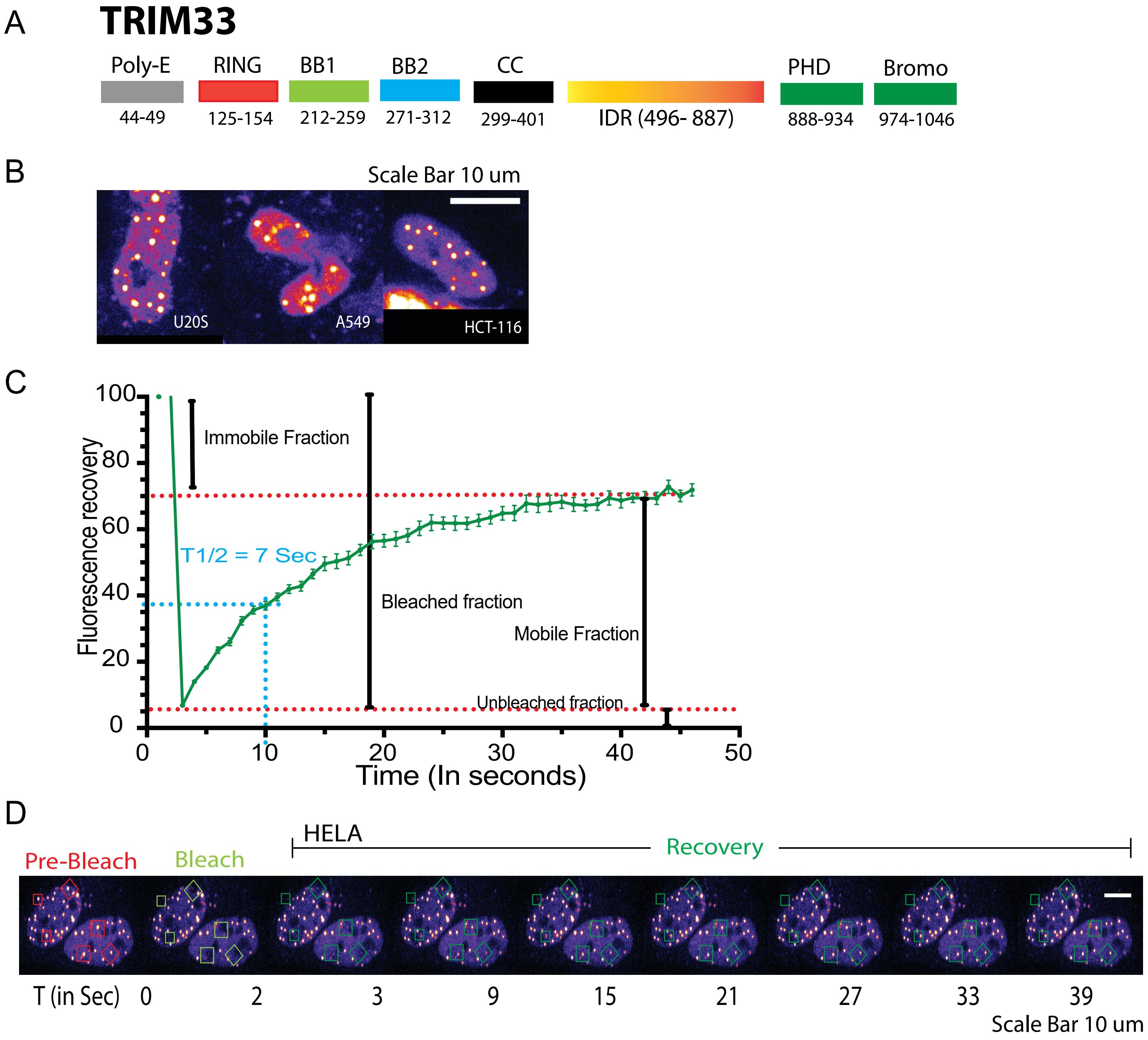
A
TRIM33
Poly-E
RING
BB1
BB2
CC
PHD
Bromo
44-49
125-154
212-259
271-312
299-401
IDR (496- 887)
888-934
974-1046
B
Scale Bar 10 um
U20S
A549
HCT-116
C
Fluorescence recovery
Immobile Fraction
T1/2 = 7 Sec
Bleached fraction
Mobile Fraction
Unbleached fraction
Time (In seconds)
D
HELA
Pre-Bleach
Bleach
Recovery
T (in Sec) 0 2 3 9 15 21 27 33 39
Scale Bar 10 um

**Supplementary Figure S10. Dynamic recovery for TRIM52.**

**A.** Domain structure of TRIM52 shown.

**B.** Representative fluorescent images of TRIM52 localization in HCT-116 cells.

**C.** Fluorescence recovery after photo-bleaching curve shown for TRIM52.

**D.** Representative images of fluorescence recovery after photo-bleaching shown for TRIM52.

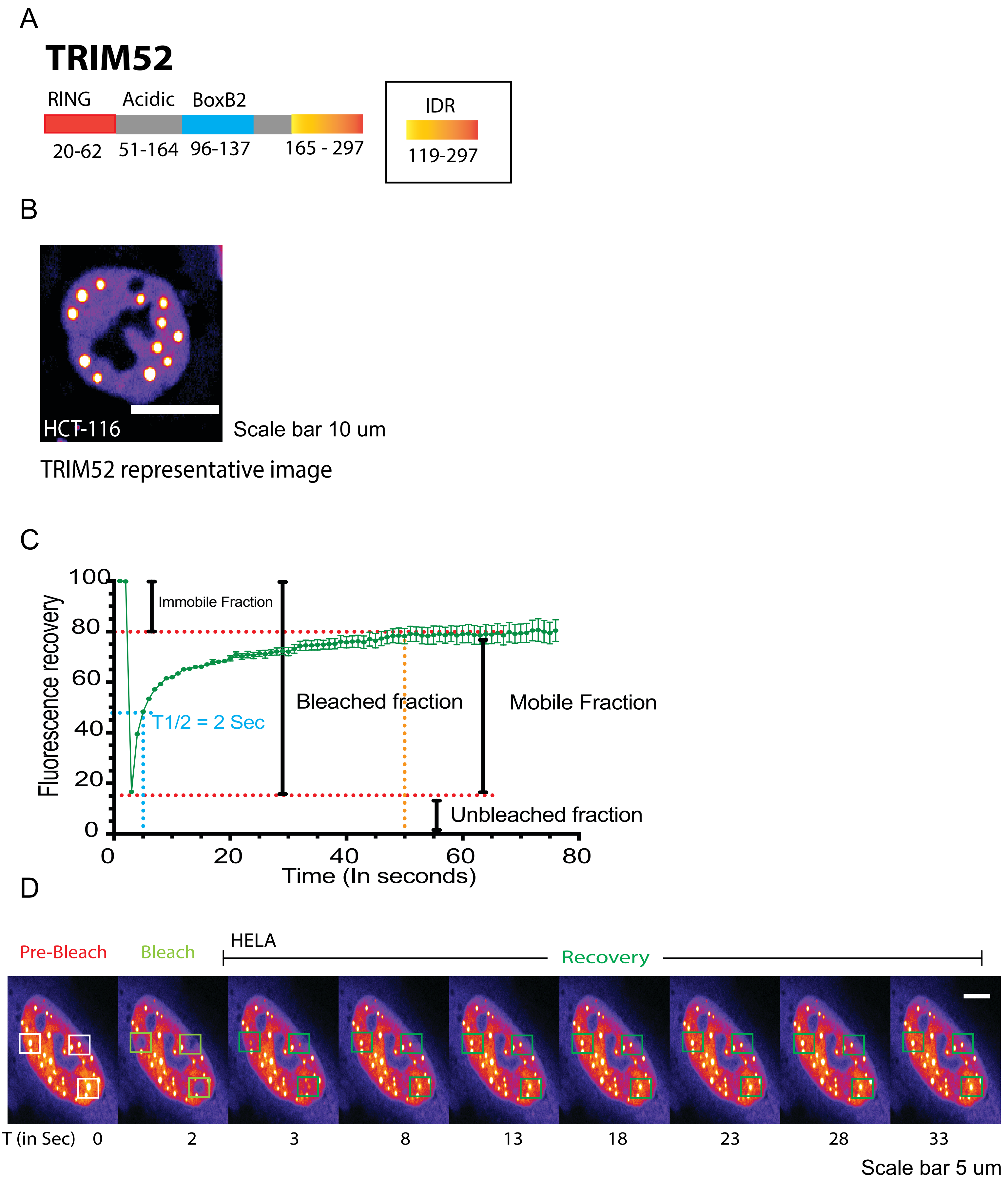

A
TRIM52
RING
Acidic
BoxB2
20-62
51-164
96-137
165 - 297
IDR
119-297
B
HCT-116
Scale bar 10 um
TRIM52 representative image
C
Fluorescence recovery
Immobile Fraction
Bleached fraction
Mobile Fraction
T1/2 = 2 Sec
Unbleached fraction
0
20
40
60
80
100
Time (In seconds)
D
Pre-Bleach
Bleach
HELA
Recovery
T (in Sec)
0
2
3
8
13
18
23
28
33
Scale bar 5 um

#### Supplementary Figure S11. Dynamic recovery for TRIM28.

**A.** Domain structure of TRIM28 shown.

**B.** Representative images of fluorescence recovery after photo-bleaching shown for TRIM28.

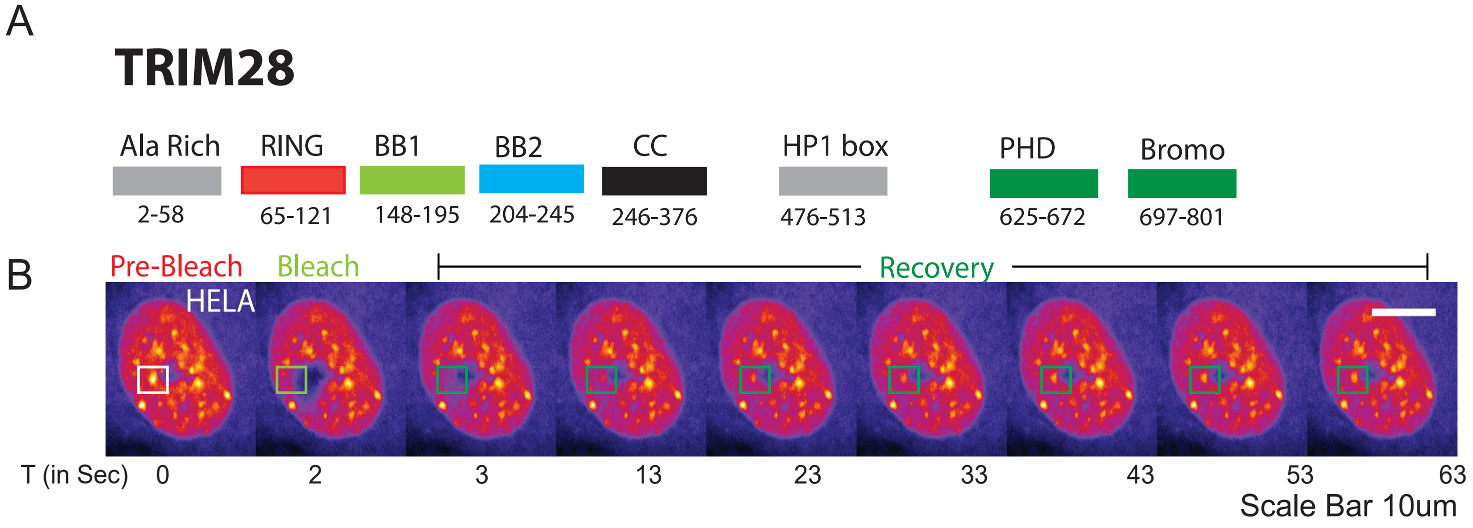
A
TRIM28
Ala Rich
2-58
RING
65-121
BB1
148-195
BB2
204-245
CC
246-376
HP1 box
476-513
PHD
625-672
Bromo
697-801
B
Pre-Bleach
Bleach
Recovery
HELA
T (in Sec)
0
2
3
13
23
33
43
53
63
Scale Bar 10um

**Supplementary Figure S12. *In silico* prediction of disordered content across TRIM family.**

**A.** Heatmap showing relative scores or percentages generated from IDR tools FuzDrop-pLLPS, PICNIC, and PONDR-VSL2 (percent disordered).

**B.** PICNIC scores were plotted from highest to lowest by TRIM family protein.

**C.** PICNIC scores were plotted for subset of TRIM family proteins.

**D.** Based on recovery dynamics after photobleaching, TRIMs forming nuclear and/or cytoplasmic bodies were grouped into dynamic or non-dynamic bodies. Plots show disordered content scores/percentages for each group. Disordered content/percentages are significantly higher in the dynamic bodies, and in particular in dynamic nuclear bodies, than in non-dynamic ones. Mann-Whitney U test. *$p<0.05$, **$p<0.01$, ****$p<0.0001$.

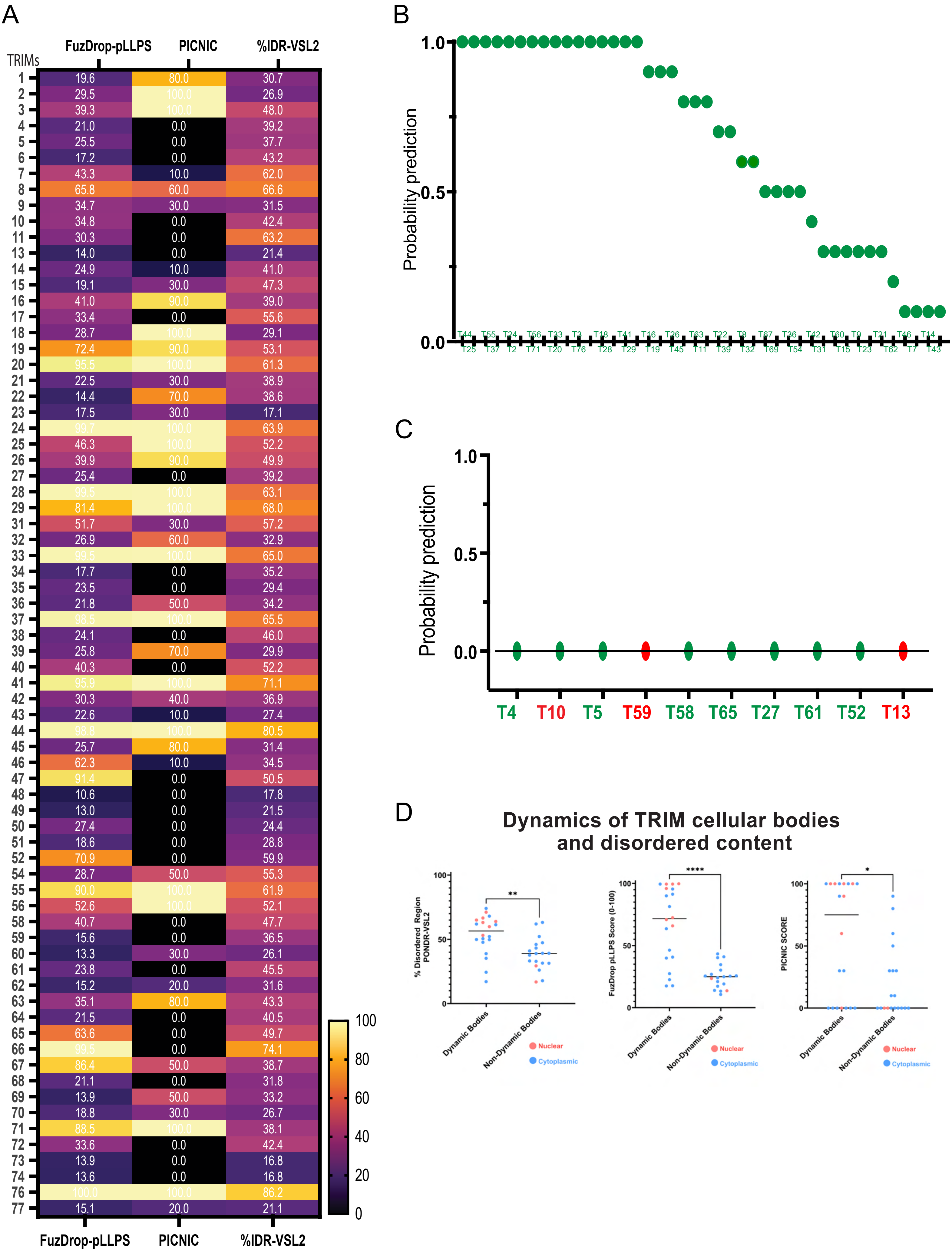

A
FuzDrop-pLLPS
PICNIC
%IDR-VSL2
TRIMs
1 19.6 80.0 30.7
2 29.5 100.0 26.9
3 39.3 100.0 48.0
4 21.0 0.0 39.2
5 25.5 0.0 37.7
6 17.2 0.0 43.2
7 43.3 10.0 62.0
8 65.8 60.0 66.6
9 34.7 30.0 31.5
10 34.8 0.0 42.4
11 30.3 0.0 63.2
13 14.0 0.0 21.4
14 24.9 10.0 41.0
15 19.1 30.0 47.3
16 41.0 90.0 39.0
17 33.4 0.0 55.6
18 28.7 100.0 29.1
19 72.4 90.0 53.1
20 95.5 100.0 61.3
21 22.5 30.0 38.9
22 14.4 70.0 38.6
23 17.5 30.0 17.1
24 99.7 100.0 63.9
25 46.3 100.0 52.2
26 39.9 90.0 49.9
27 25.4 0.0 39.2
28 99.5 100.0 63.1
29 81.4 100.0 68.0
31 51.7 30.0 57.2
32 26.9 60.0 32.9
33 99.5 100.0 65.0
34 17.7 0.0 35.2
35 23.5 0.0 29.4
36 21.8 50.0 34.2
37 98.5 100.0 65.5
38 24.1 0.0 46.0
39 25.8 70.0 29.9
40 40.3 0.0 52.2
41 95.9 100.0 71.1
42 30.3 40.0 36.9
43 22.6 10.0 27.4
44 98.8 100.0 80.5
45 25.7 80.0 31.4
46 62.3 10.0 34.5
47 91.4 0.0 50.5
48 10.6 0.0 17.8
49 13.0 0.0 21.5
50 27.4 0.0 24.4
51 18.6 0.0 28.8
52 70.9 0.0 59.9
54 28.7 50.0 55.3
55 90.0 100.0 61.9
56 52.6 100.0 52.1
58 40.7 0.0 47.7
59 15.6 0.0 36.5
60 13.3 30.0 26.1
61 23.8 0.0 45.5
62 15.2 20.0 31.6
63 35.1 80.0 43.3
64 21.5 0.0 40.5
65 63.6 0.0 49.7
66 99.5 0.0 74.1
67 86.4 50.0 38.7
68 21.1 0.0 31.8
69 13.9 50.0 33.2
70 18.8 30.0 26.7
71 88.5 100.0 38.1
72 33.6 0.0 42.4
73 13.9 0.0 16.8
74 13.6 0.0 16.8
76 100.0 100.0 86.2
77 15.1 20.0 21.1
FuzDrop-pLLPS PICNIC %IDR-VSL2
100 80 60 40 20 0
B
Probability prediction
1.0 0.5 0.0
T44 T55 T24 T56 T33 T3 T18 T41 T16 T26 T63 T22 T8 T67 T36 T42 T60 T9 T21 T46 T14
T25 T37 T2 T71 T20 T76 T28 T29 T19 T45 T11 T39 T32 T69 T54 T31 T15 T23 T62 T7 T43
C
Probability prediction
1.0 0.5 0.0
T4 T10 T5 T59 T58 T65 T27 T61 T52 T13
D
Dynamics of TRIM cellular bodies and disordered content
% Disordered Region PONDR-VSL2
FuzDrop pLLPS Score (0-100)
PICNIC SCORE
100 50 0
**
****
*
Dynamic Bodies
Non-Dynamic Bodies
Nuclear
Cytoplasmic

**Supplementary Figure S13. Variation in IDR caused by alternative splicing of *TRIM19.*** Shown are the last coding positions (arrows) for TRIM19 naturally occurring splice isoforms, resulting in gradual loss of the C-terminal structured domain and IDR. Representative images of TRIM19 localization in HELA cells are shown. Alpha-fold structures are shown below.

### Physiological IDR changes via alternative splicing

Human TRIM19 - WT

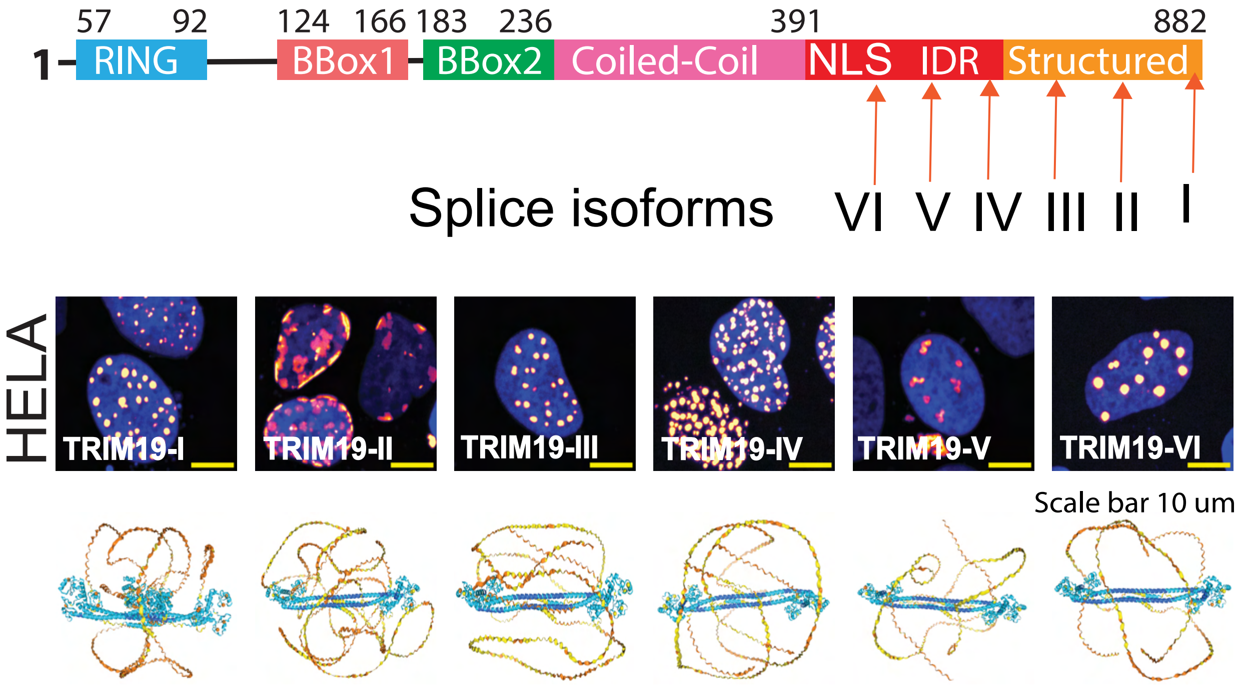


AF3 dimers of TRIM19 splice isoforms

**Supplementary Figure S14. *In cellulo* concentration measurements.** Image analysis pipeline (**A**) and the concentration estimation standard curve **(B)** showing the linear concentration range are presented.

A Microscopy image analysis pipeline

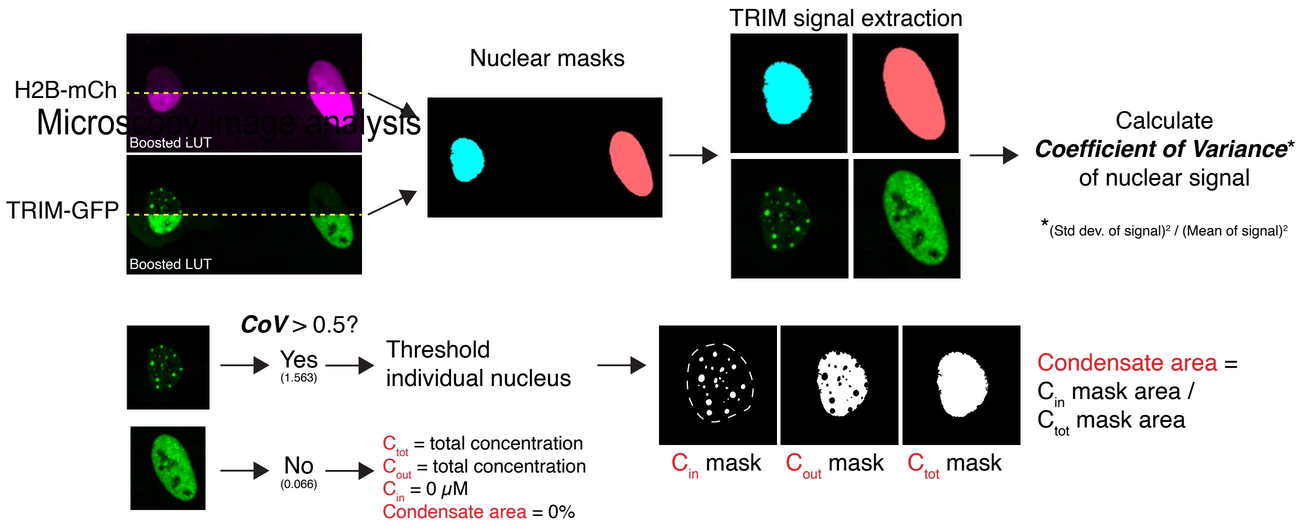


B *In cellulo* concentration estimation pipeline

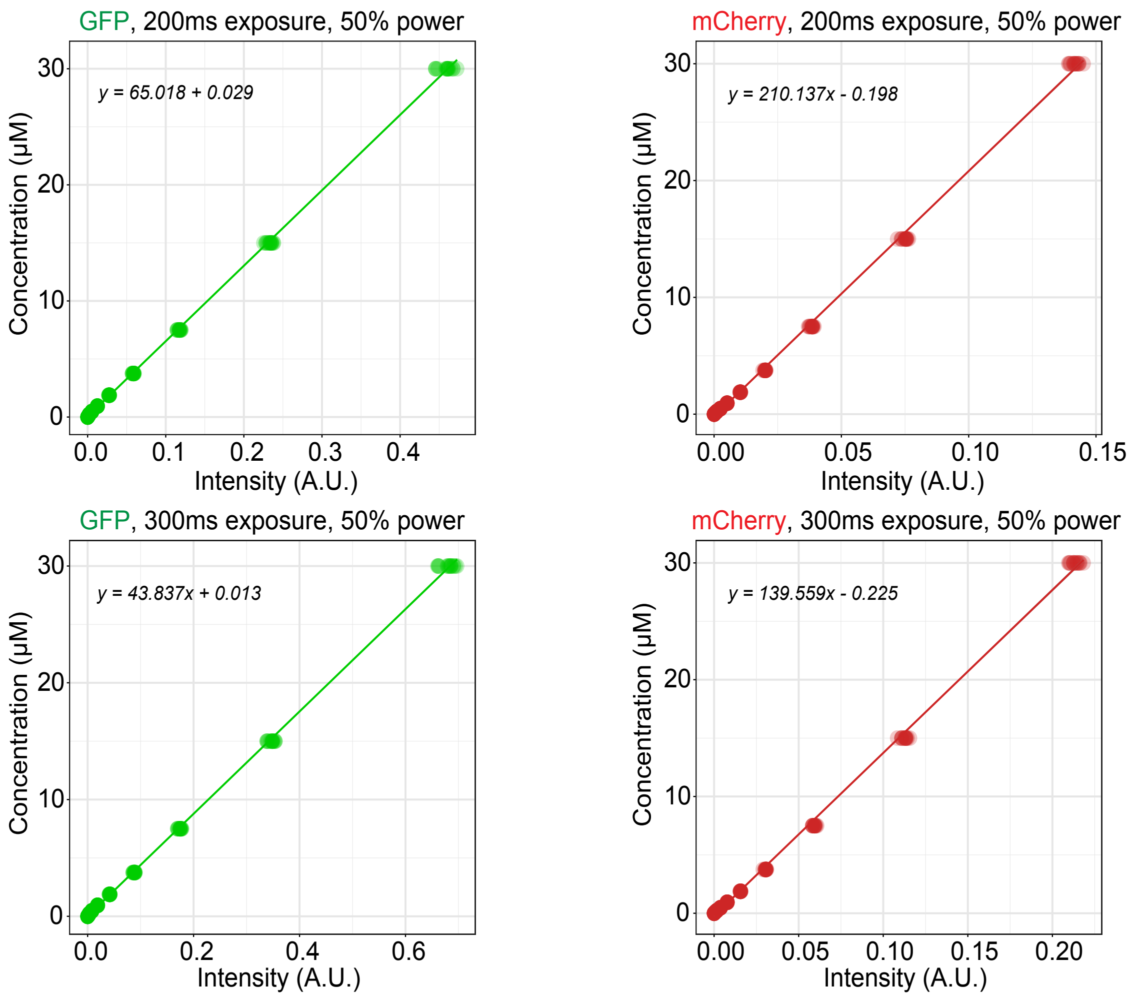

**Supplementary Figure S15. TRIM8 protein concentration-localization relationship and impact of patient variants.**

**A.** Relationship between TRIM8 concentration in U2OS cells over time and representative images showing increasing condensate formation.

**B.** Shown are the representative images of expression constructs of TRIM8 and mutants based on the TRIM8 syndrome caused by IDR deletions resulting from mutations leading to the stop codons in the IDR of the TRIM8 protein causing premature truncations. To the right are Alpha fold3 models of the corresponding disease mutants. C1461G is same as TRIM8-487* and del-T1163 is same as TRIM8-388*. All truncations cause partial to complete loss of TRIM8 nuclear condensate formation.

**C**. A representative summary of all the tested TRIM8 disease variants at the level of mRNA that led to the corresponding TRIM8 C-terminal IDR truncations., TRIM8-WT (1-551) showing nuclear bodies. TRIM8-IDR scramble also showing nuclear bodies saying IDR amino acid content is a more dominant driver of condensation than sequence patterns. shows the representative images for the transfections of the exact nucleotide change in the full-length sequence of the TRIM8 coding gene as in the patients, to see if any post-transcriptional mechanisms are at play, all of them show loss of condensation without causing any post-transcriptional change like non-sense mediated decay.

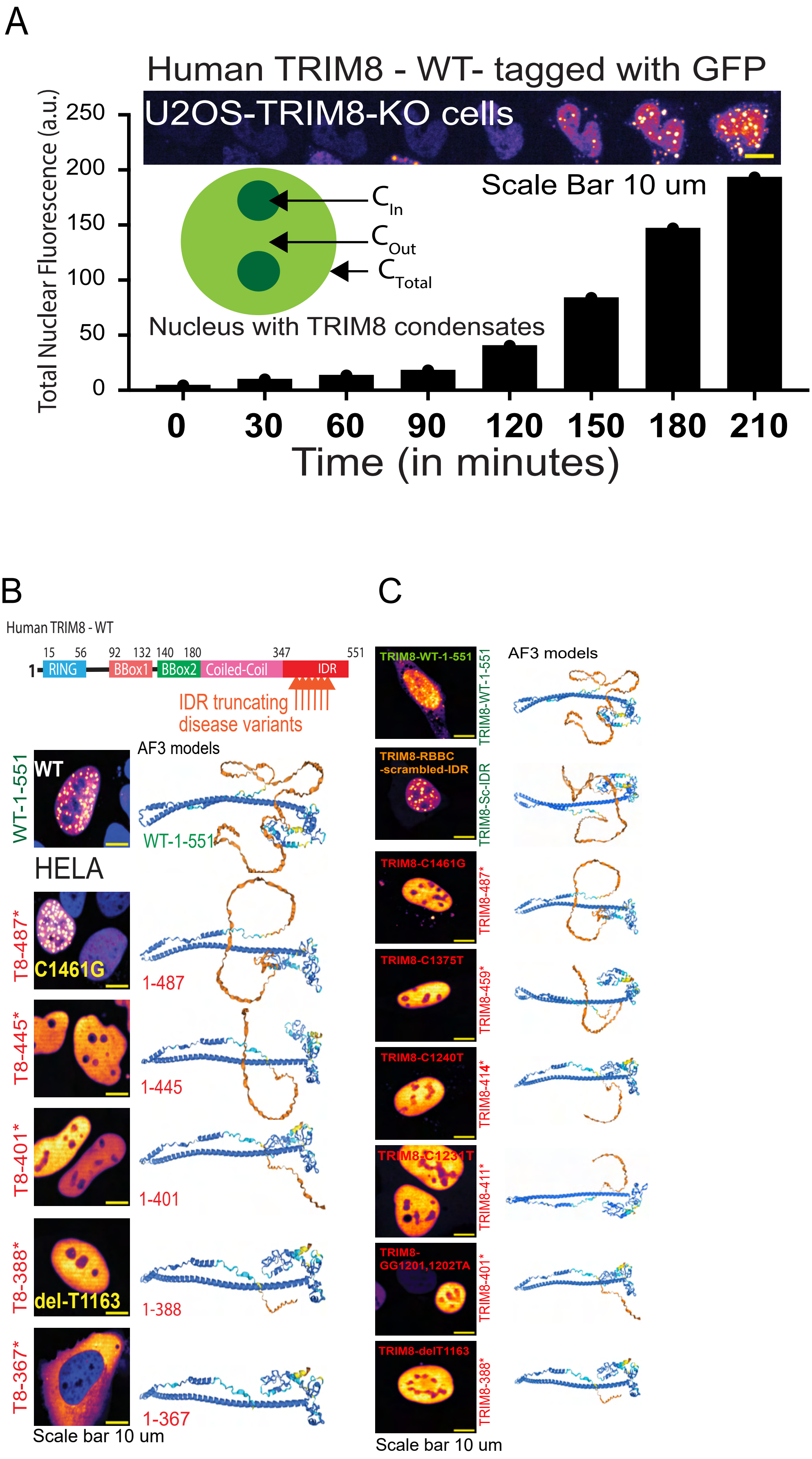

A
Human TRIM8 - WT- tagged with GFP
U2OS-TRIM8-KO cells
Scale Bar 10 um
Total Nuclear Fluorescence (a.u.)
250
200
150
100
50
0
C In
C Out
C Total
Nucleus with TRIM8 condensates
0
30
60
90
120
150
180
210
Time (in minutes)
B
Human TRIM8 - WT
15 56 92 132 140 180 347 551
1
RING
BBox1
BBox2
Coiled-Coil
IDR
IDR truncating disease variants
AF3 models
WT-1-551
WT
WT-1-551
HELA
T8-487*
C1461G
1-487
T8-445*
1-445
T8-401*
1-401
T8-388*
del-T1163
1-388
T8-367*
1-367
Scale bar 10 um
C
AF3 models
TRIM8-WT-1-551
TRIM8-WT-1-551
TRIM8-RBBC -scrambled-IDR
TRIM8-Sc-IDR
TRIM8-C1461G
TRIM8-487*
TRIM8-C1375T
TRIM8-459*
TRIM8-C1240T
TRIM8-414*
TRIM8-C1231T
TRIM8-411*
TRIM8-GG1201,1202TA
TRIM8-401*
TRIM8-delT1163
TRIM8-388*
Scale bar 10 um

**Supplementary Figure S16. TRIM8 disease variants disrupt the turnover and condensation of TRIM8.**

**A.** Upon transfection in immortalized podocytes, protein levels of Flag-tagged wildtype TRIM8 but not patient variant TRIM8 were increased by the 26S proteasome inhibitor MG132 in a dose dependent manner, while TRIM8 patient variant protein levels were unaffected by MG132 treatment, indicating a loss of proteasome-dependent regulation. 13-actin levels demonstrate equal loading.

**B.** Densitometry of cycloheximide studies in **Figure 5C** were performed from 3 biological replicates showing wildtype TRIM8 has a half-life of ~3 hours while mutant TRIM8 half-life was >5 hours. Significantly different levels were observed at 3-5 hours by the Mann–Whitney U test.

**C.** Ectopic-tagged TRIM8 protein was expressed in immortalized podocytes upon doxycycline administration. In correlation with increased total wildtype TRIM8 levels, proteasome inhibition causes increased nuclear condensation of wildtype TRIM8. In contrast, modulation of the proteasome with MG132 does not affect patient variant TRIM8 protein condensation and localization. Localization of TRIM8 is quantified from multiple biological replicates and demonstrates the selective regulation of wildtype TRIM8 condensation by the 26S proteasome.

**D.** In correlation with increased total wildtype TRIM8 levels in **Figure 5B**, proteasome inhibition causes increased nuclear condensation of transiently transfected wildtype TRIM8. In contrast, modulation of the proteasome with MG132 did not markedly affect patient variant TRIM8 protein condensation and localization. Localization of TRIM8 is quantified from multiple biological replicates and demonstrates the selective regulation of wildtype TRIM8 condensation by the 26S proteasome.

**E.** Immunoblots are shown of inputs for TRIM8 poly-ubiquitination assays performed in Figure 5D and demonstrate appropriate expression of FLAG-tagged TRIM8 (T8) and HA-tagged ubiquitin (Ubi).

**F.** Schematic representation of the TRIM8 wildtype protein structure, illustrating its key domains. Below each black line indicates the position of a lysine residue in lysine residue including two lysine residues after positions of nonsense mutations identified in TRIM8-associated syndrome.

**G.** Upon transient transfection in immortalized podocytes, protein levels of Flag-tagged wildtype TRIM8 and C-terminal lysine variants were increased by the 26S proteasome inhibitor MG132 in a dose-dependent manner, suggesting these lysine residues are not required for proteasomal degradation of TRIM8. β-actin levels demonstrate equal loading.

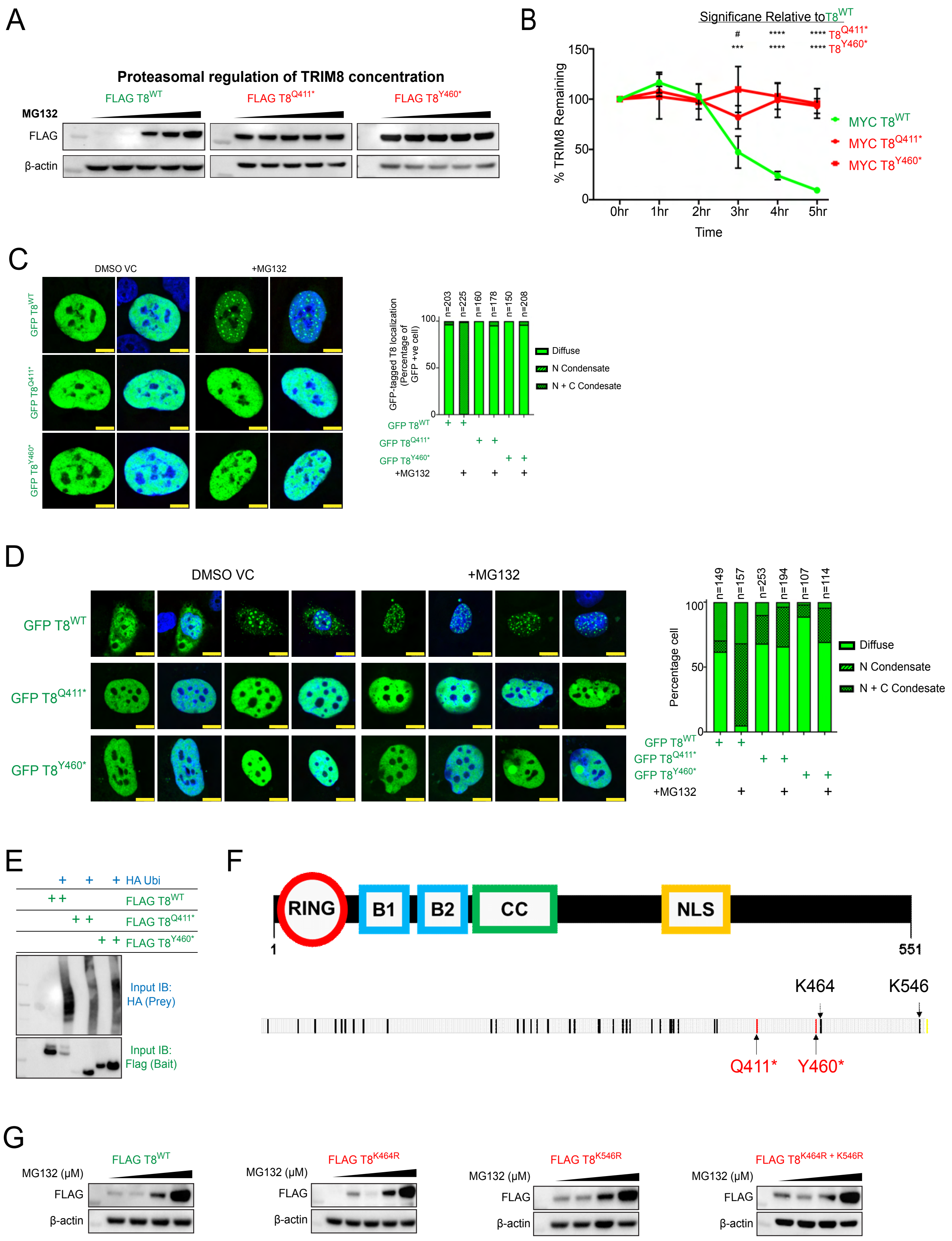
A
Proteasomal regulation of TRIM8 concentration
FLAG T8WT
FLAG T8Q411*
FLAG T8Y460*
MG132
FLAG
β-actin
B
Significane Relative to T8WT
# **** **** T8Q411*
*** **** **** T8Y460*
% TRIM8 Remaining
150
100
50
0
0hr 1hr 2hr 3hr 4hr 5hr
Time
MYC T8WT
MYC T8Q411*
MYC T8Y460*
C
DMSO VC
+MG132
GFP T8WT
GFP T8Q411*
GFP T8Y460*
GFP-tagged T8 localization (Percentage of GFP +ve cell)
n=203 n=225 n=160 n=178 n=150 n=208
Diffuse
N Condensate
N + C Condesate
+MG132
D
DMSO VC
+MG132
GFP T8WT
GFP T8Q411*
GFP T8Y460*
Percentage cell
n=149 n=157 n=253 n=194 n=107 n=114
Diffuse
N Condensate
N + C Condesate
+MG132
E
HA Ubi
FLAG T8WT
FLAG T8Q411*
FLAG T8Y460*
Input IB: HA (Prey)
Input IB: Flag (Bait)
F
RING
B1
B2
CC
NLS
1
551
K464
K546
Q411*
Y460*
G
FLAG T8WT
FLAG T8K464R
FLAG T8K546R
FLAG T8K464R + K546R
MG132 (μM)
FLAG
β-actin

**Supplementary Figure S17. Components of ubiquitin-proteasome system co-localize to TRIM8 condensates in immortalized podocytes.**

**A.** GFP-tagged ubiquitin co-localizes to Cherry-tagged wildtype TRIM8 condensates. Quantification of co-localization is displayed in the histogram.

**B.** Cherry-tagged wildtype TRIM8 promotes localization of GFP-tagged ubiquitin to nuclear condensates in immortalized podocytes, which is abrogated by patient variants lacking the TRIM8 IDR. The number and size of GFP-tagged ubiquitin nuclear condensates are quantified, showing wildtype TRIM8 promotes the size of these condensates while patient variant TRIM8 impairs their number and size [ ] (TRIM8 = T8, Ubiquitin = Ubi, etc)

**C.** Endogenous proteasome component PSMD4 co-localizes to Cherry-tagged wildtype TRIM8 condensates. Quantification of co-localization is displayed in the histogram.

**D.** Endogenous proteasome component PSMD12 co-localizes to Cherry-tagged wildtype TRIM8 condensates. Quantification of co-localization is displayed in the histogram.

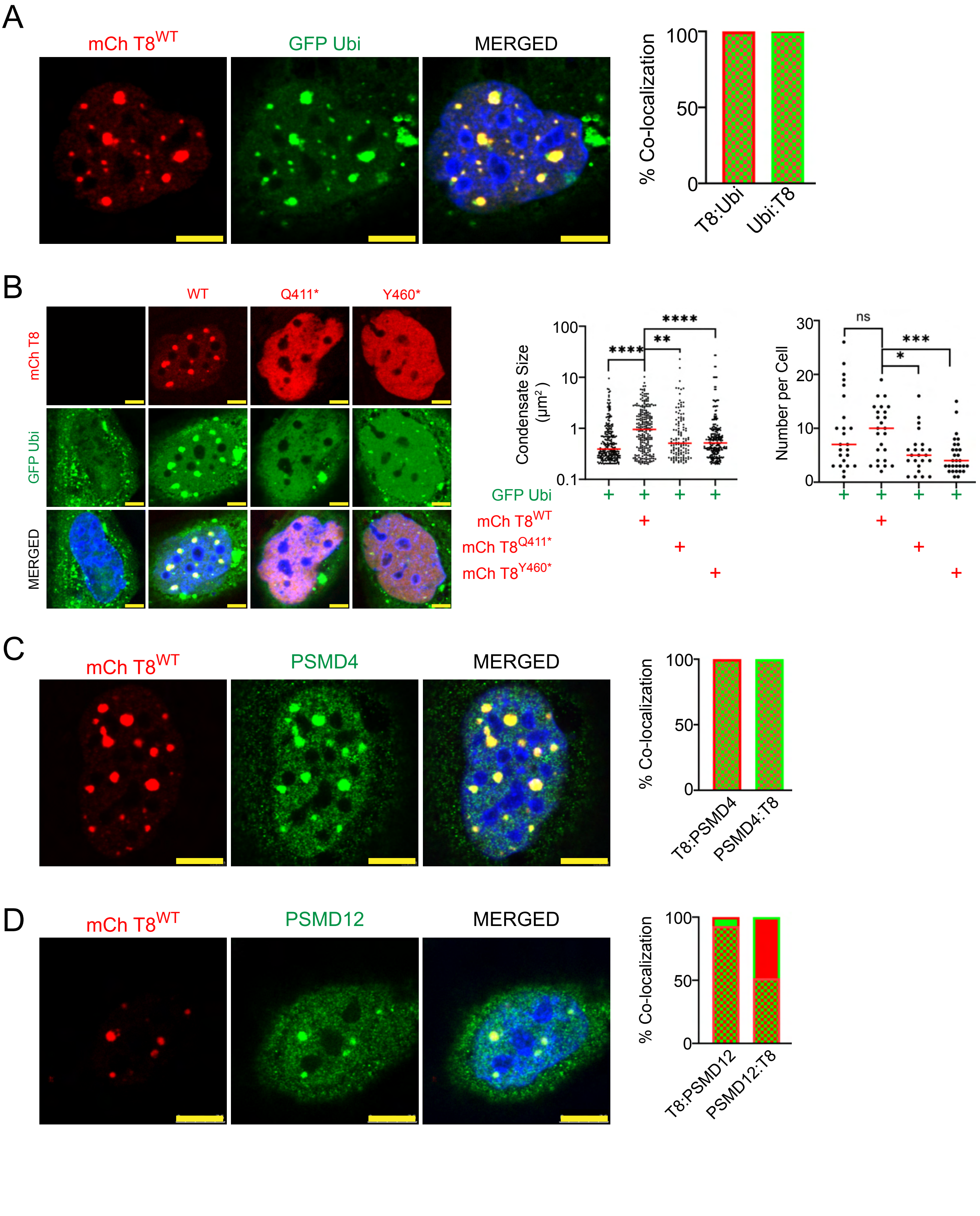
A
mCh T8WT
GFP Ubi
MERGED
% Co-localization
100
50
0
T8:Ubi
Ubi:T8
B
WT
Q411*
Y460*
mCh T8
GFP Ubi
MERGED
Condensate Size (μm²)
100
10
1
0.1
****
**
****
Number per Cell
30
20
10
0
ns
***
*
GFP Ubi
mCh T8WT
mCh T8Q411*
mCh T8Y460*
C
mCh T8WT
PSMD4
MERGED
% Co-localization
100
50
0
T8:PSMD4
PSMD4:T8
D
mCh T8WT
PSMD12
MERGED
% Co-localization
100
50
0
T8:PSMD12
PSMD12:T8

**Supplementary Figure S18. Components of ubiquitin-proteasome system co-localize to TRIM8 condensates in U2OS cells.**

This figure shows double transfection of RFP-tagged ubiquitin with GFP tagged TRIM8 constructs including the wildtype protein, a disease mutant and synthetic constructs. Localization of ubiquitin to TRIM8 condensates was impaired by patient variants.

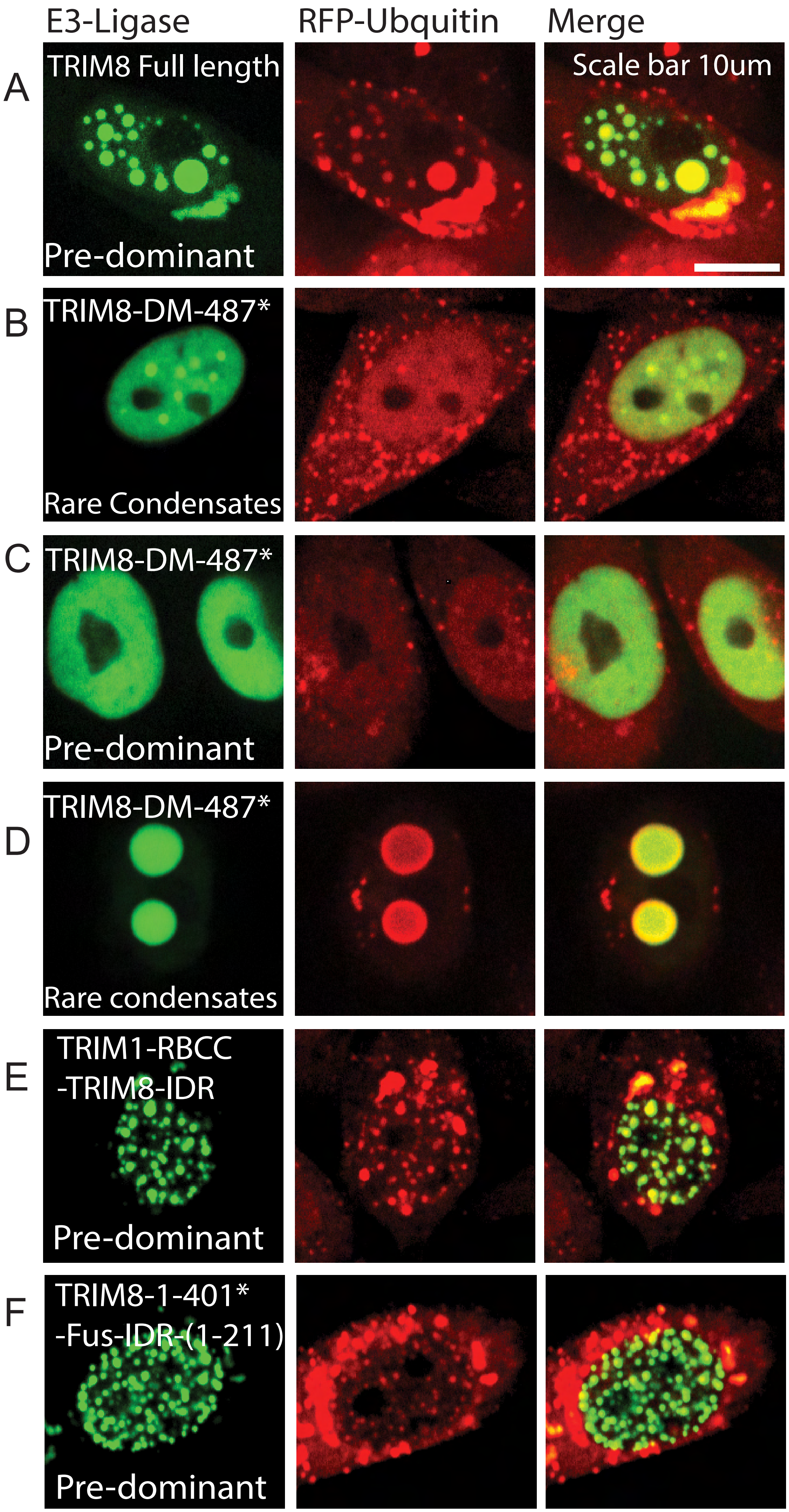
E3-Ligase
RFP-Ubquitin
Merge
A
TRIM8 Full length
Pre-dominant
Scale bar 10um
B
TRIM8-DM-487*
Rare Condensates
C
TRIM8-DM-487*
Pre-dominant
D
TRIM8-DM-487*
Rare condensates
E
TRIM1-RBCC
-TRIM8-IDR
Pre-dominant
F
TRIM8-1-401*
-Fus-IDR-(1-211)
Pre-dominant

**Supplementary Figure S19. TAK1/NFκB signaling impacted by TRIM8 disease mutations.**

**A.** GFP-tagged wildtype TRIM8 co-localizes to Cherry-tagged TAK1 condensates. Quantification of colocalization studies from is displayed in the histogram.

**B.** NF-kB transcriptional activity was assessed by a 4xNFkB binding site firefly luciferase reporter in immortalized human podocytes (HsPd) with renilla luciferase co-transfected as an internal control. The ratio of firefly: renilla activity normalized to the level for the mock MYC-tagged plasmid transfection group of all three biological replicates is shown. NF-kB is increased by co-transfection with wildtype TRIM8 while TRIM8 patient variants failed to modulate reporter activity, indicating this effect is dependent on the TRIM8 IDR. **$p<0.01$ by t-test.

**C.** Co-localization of TAK1 as a function of the indicated TRIM8 protein variants in physiology (WT), disease (disease mutants) and engineered rescue (TRIM1 RBBC swap and Fus IDR swap) in U2OS cells is shown.

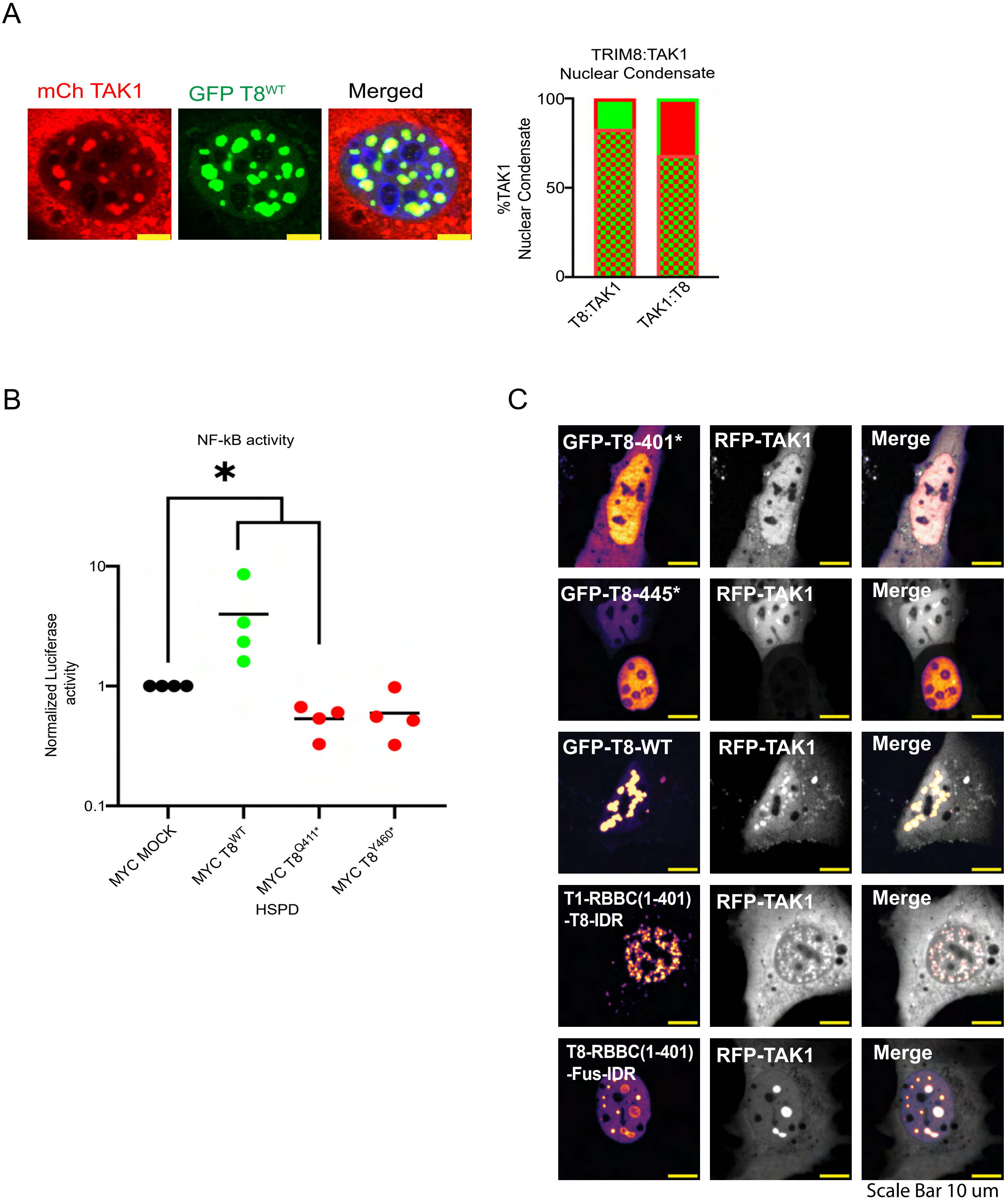
A
mCh TAK1
GFP T8WT
Merged
TRIM8:TAK1
Nuclear Condensate
%TAK1
Nuclear Condensate
100
50
0
T8:TAK1
TAK1:T8
B
NF-kB activity
*
Normalized Luciferase
activity
10
1
0.1
MYC MOCK
MYC T8WT
MYC T8Q411*
MYC T8Y460*
HSPD
C
GFP-T8-401*
RFP-TAK1
Merge
GFP-T8-445*
RFP-TAK1
Merge
GFP-T8-WT
RFP-TAK1
Merge
T1-RBBC(1-401)
-T8-IDR
RFP-TAK1
Merge
T8-RBBC(1-401)
-Fus-IDR
RFP-TAK1
Merge
Scale Bar 10 um

**Supplementary Figure S20. CRISPR/Cas9 mediated generation of *TRIM8* variants in immortalized human podocytes and cellular consequences.**

**A.** Diagram shows exon structure of human TRIM8 locus with relative position of gRNA (pink arrows) employed for CRISPR/Cas9 gene editing in exon 1 and exon 6 to generate N-terminal and C-terminal truncating variants, respectively. Primer positions are shown (blue-green bars), which were employed for validation of genetic variants.

**B.** Gel electrophoresis is shown of PCR products (343bp) of genomic DNA isolated from immortalized podocytes CRISPR-generated N-terminal TRIM8 truncating variants (*TRIM8$^{Ex1-/Ex1-}$*). *TRIM8$^{Ex1-/Ex1-}$* podocytes (two independent cell lines) exhibited PCR products of reduced size relative to wildtype genomic DNA using primers bounding an exon 1 deletion region.

**C.** *TRIM8$^{Ex1-/Ex1-}$* podocytes (two independent cell lines) showed no PCR product (160bp) when using a forward primer that was internal to the deletion.

**D.** Gel electrophoresis is shown of PCR products (845bp) of genomic DNA isolated from immortalized podocytes CRISPR-generated C-terminal TRIM8 truncating variants (*TRIM8$^{Ex6-/Ex6-}$*). *TRIM8$^{Ex6-/Ex6-}$* podocytes (two independent cell lines) exhibited PCR products of reduced size relative to wildtype genomic DNA using primers bounding an exon 6 deletion region.

**E.** *TRIM8$^{Ex6-/Ex6-}$* podocytes (two independent cell lines) showed no PCR product (291bp) when using a forward primer that was internal to the deletion.

**F.** Representative Sanger sequencing chromatogram of PCR product of genomic DNA from *TRIM8$^{Ex1-/Ex1-}$* podocytes.

**G.** Representative Sanger sequencing chromatogram of PCR product of genomic DNA from *TRIM8$^{Ex6-/Ex6-}$* podocytes.

**H.** Diagram of human TRIM8 protein is shown with bars above representing immunogens for N-terminal NOVUS (aa 205-aa285) and C-terminal Santa Cruz antibodies in relation to disease associated truncating variants.

**I.** Western blotting of total cell lysates from *TRIM8$^{Ex1-/Ex1-}$*and *TRIM8$^{Ex6-/Ex6-}$* podocytes was performed. Using an antibody raised against the TRIM8 N-terminus, wildtype lysates exhibited a predominant band at ~60-65 kDa consistent with the wildtype protein that was absent in *TRIM8$^{Ex1-/Ex1-}$* podocytes and reduced in size in *TRIM8$^{Ex6-/Ex6-}$* podocytes.

**J.** Using an antibody raised against the TRIM8 C-terminus, the expected wildtype protein was observed in wildtype lysates and absent in TRIM8 mutant cell lines.

**K.** Cell proliferation was assessed by daily cell count over time in *TRIM8$^{Ex1-/Ex1-}$* podocytes. Cell proliferation increased in TRIM8 mutant podocytes relative to control wildtype podocytes upon cell seeding at low density (5,000 cells/well).

**L.** Cell proliferation was assessed by daily cell count over time in *TRIM8$^{Ex6-/Ex6-}$* podocytes. Cell proliferation increased in TRIM8 mutant podocytes relative to control wildtype podocytes upon cell seeding at low density (5,000 cells/well).

**M.** Cell proliferation was assessed by daily cell count over time in *TRIM8*$^{Ex1-/Ex1-}$ podocytes. Cell proliferation was not significantly different between TRIM8 mutant podocytes and control wildtype podocytes upon cell seeding at higher density (10,000 cells/well).

**N.** Cell proliferation was assessed by daily cell count over time in *TRIM8*$^{Ex6-/Ex6-}$ podocytes. Cell proliferation was not significantly different between TRIM8 mutant podocytes and control wildtype podocytes upon cell seeding at higher density (10,000 cells/well).

**O.** Cell metabolic activity by MTT assay, as measure of proliferation, was not consistently increased in independent *TRIM8*$^{Ex1-/Ex1-}$ and *TRIM8*$^{Ex6-/Ex6-}$ podocytes at low density (5,000 cells/well). **p<0.01 by Mann Whitney U-test.

**P.** Cell metabolic activity by MTT assay, as measure of proliferation, was consistently reduced in independent *TRIM8*$^{Ex1-/Ex1-}$ podocytes relative to control wildtype podocytes, upon seeding at higher cell density (10,000 cells/well). MTT-based cell metabolic activity was not increased consistently in two independent *TRIM8*$^{Ex6-/Ex6-}$ podocytes. **p<0.05, ****p<0.0001 by Mann Whitney U-test.

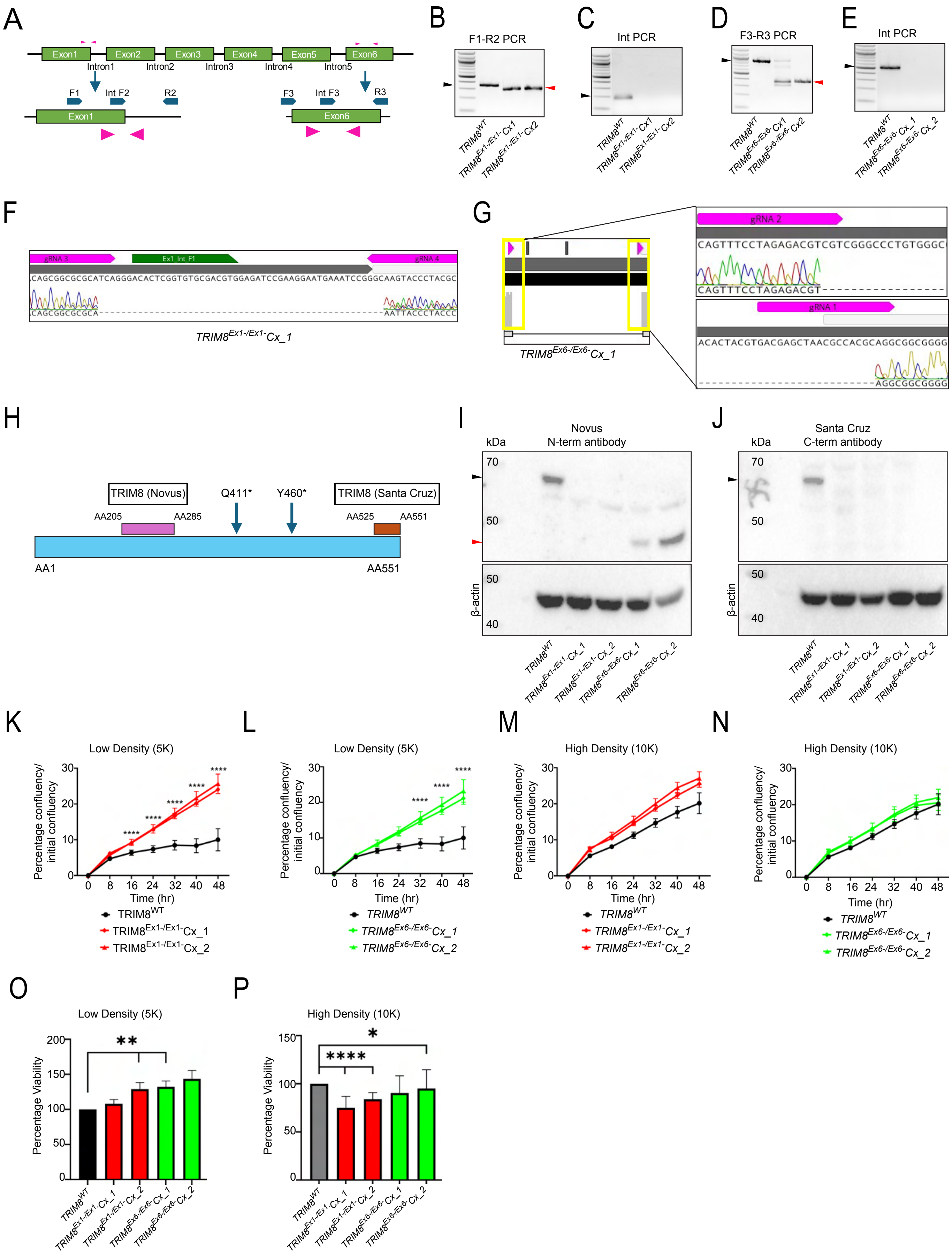
A
Exon1
Exon2
Exon3
Exon4
Exon5
Exon6
Intron1
Intron2
Intron3
Intron4
Intron5
F1
Int F2
R2
F3
Int F3
R3
B
F1-R2 PCR
C
Int PCR
D
F3-R3 PCR
E
Int PCR
F
gRNA 3
Ex1_Int_F1
gRNA 4
CAGCGGCGCGCATCAGGGACACTCGGTGTGCGACGTGGAGATCCGAAGGAATGAAATCCGGGCAAGTACCCTACGC
CAGCGGCGCGCA
AATTACCCTACCC
G
gRNA 2
CAGTTTCCTAGAGACGTCGTCGGGCCCTGTGGGC
CAGTTTCCTAGAGACGT
gRNA 1
ACACTACGTGACGAGCTAACGCCACGCAGGCGGCGGGG
AGGCGGCGGGG
H
TRIM8 (Novus)
Q411*
Y460*
TRIM8 (Santa Cruz)
AA205
AA285
AA525
AA551
AA1
AA551
I
kDa
Novus
N-term antibody
70
50
50
40
β-actin
J
kDa
Santa Cruz
C-term antibody
70
50
50
40
β-actin
K
Low Density (5K)
L
Low Density (5K)
M
High Density (10K)
N
High Density (10K)
Percentage confluency/ initial confluency
Time (hr)
0 8 16 24 32 40 48
0 10 20 30
****
O
Low Density (5K)
Percentage Viability
0 50 100 150 200
**
P
High Density (10K)
Percentage Viability
0 50 100 150
****
*

**Supplementary Figure S21. Derivation and characterization of FVB/N mice bearing CRISPR/Cas9 mediated N-terminal TRIM8 truncating variants by CRISPR/Cas9 editing.**

**A.** Diagram shows exon structure of mouse Trim8 locus (Chr19:46490087-46504894 bp) with N-terminal deletions (yellow bars) generated by multiguide CRISPR/Cas9 gene editing approach in exon 1 and PCR primers used for genotyping (blue bars).

**B.** Gel electrophoresis is shown of PCR product from neonatal toe tissue genomic DNA from wildtype (127 bp), heterozygous, and homozygous mice (514 bp).

**C.** Diagram shows exon structure of mouse Trim8 and PCR Primer spanning through exon1(Ex1) to exon3(Ex3) (pink arrow) and exon3 to exon5(Ex5) (blue arrow) used for RT-PCR.

**D.** Total RNA from neonatal kidney was extracted followed by reverse transcription of first strand cDNA. PCR amplification of regions spanning exon 1-3 and exon 3-5 of Trim8 transcripts are shown. RT-PCR products are reduced in $Trim8^{Ex1m/Ex1m}$ homozygote mice.

**E.** Western blotting of total lysates from $Trim8^{Ex1m}$ mouse brain and kidney tissue was performed. Using an antibody raised against the TRIM8 N-terminus, wildtype lysates exhibited a predominant band at ~60-65 KDa consistent with the wildtype protein that was absent in $Trim8^{Ex1-/Ex1-}$ lysates.

**F.** Western blotting of total lysates from $Trim8^{Ex1m}$ mouse brain and kidney tissue was performed. Using an antibody raised against the TRIM8 C-terminus, wildtype lysates exhibited a predominant band at ~60-65 KDa consistent with the wildtype protein that was absent in $Trim8^{Ex1-/Ex1}$ lysates.

**G.** Mendelian ratios observed at weaning age (chi square values = 2.297 with 2 degrees of freedom).

**H.** Kaplan-Meyer Curve shows no difference in survival of wildtype, heterozygote and homozygote $Trim8^{Ex1m}$ mice over 400 days of life.

**I.** Urine albumin-to-creatine ratios as a measure of proteinuria were assessed monthly in $Trim8^{Ex1m}$ mice and were not significantly different across genotype groups over 12 months of life.

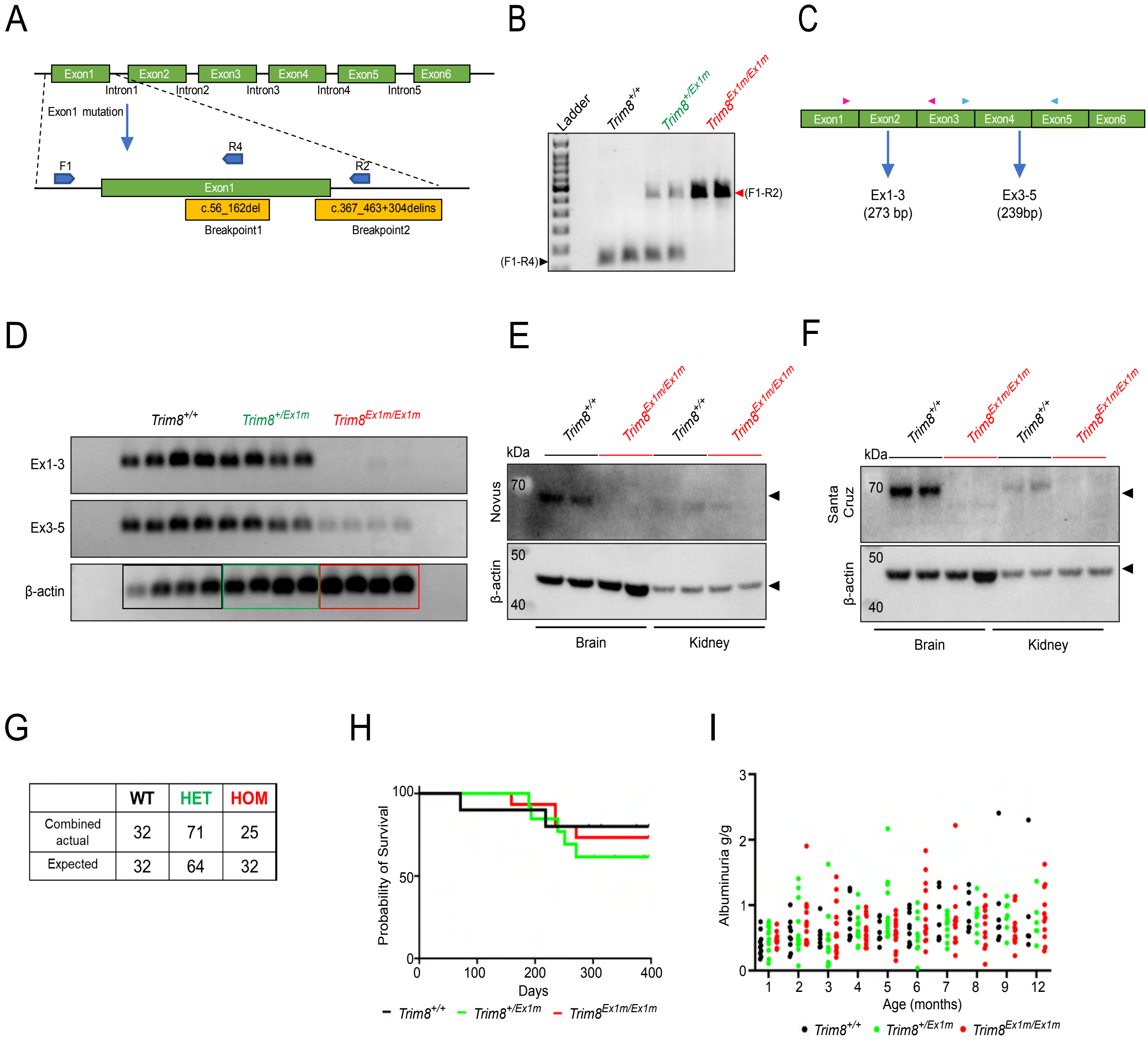

A
Exon1
Exon2
Exon3
Exon4
Exon5
Exon6
Intron1
Intron2
Intron3
Intron4
Intron5
Exon1 mutation
F1
R4
R2
c.56_162del
c.367_463+304delins
Breakpoint1
Breakpoint2
B
Ladder
Trim8+/+
Trim8+/Ex1m
Trim8Ex1m/Ex1m
(F1-R2)
(F1-R4)
C
Ex1-3
(273 bp)
Ex3-5
(239bp)
D
Ex1-3
Ex3-5
β-actin
E
kDa
Novus
70
50
40
Brain
Kidney
F
Santa Cruz
G
WT
HET
HOM
Combined actual
32
71
25
Expected
32
64
32
H
Probability of Survival
Days
I
Albuminuria g/g
Age (months)

**Supplementary Figure S22. Derivation and characterization of FVB/N mice bearing CRISPR/Cas9 mediated C-terminal TRIM8 truncating variants by CRISPR/Cas9 editing.**

**A.** Sanger sequencing chromatogram shows truncating allele within Trim8 exon 6 that is present in heterozygous and homozygous mice.

**B.** Kaplan-Meyer Curve shows no difference in survival of wildtype, heterozygote and homozygote *Trim8*$^{Ex6m}$ mice over 400 days of life.

**C.** C-terminal truncating TRIM8 variants were generated in FVB/N mice (*Trim8*$^{Ex6m}$). Mendelian ratios observed at weaning age (Chi squared equals 0.631 with 2 degrees of freedom).

**D.** Urine albumin-to-creatine ratios as measure of proteinuria were assessed monthly in *Trim8*$^{Ex6m}$ mice and were not significantly different across genotype groups over 12 months of life.

**E.** Periodic acid–Schiff (PAS) stained sections of kidneys from 12-month-old *Trim8*$^{Ex6m-/Ex6m-}$, *Trim8*$^{Ex6m\ -/+}$ and *Trim8*$^{Ex6m\ +/+}$ mice were generated. Representative images of PAS from each genotype are shown.

**F.** 20x images of glomeruli were evaluated through an automated ImageJ pipeline to determine glomerular matrix deposition in a blinded manner. *Trim8*$^{Ex6m-/Ex6m-}$ and *Trim8*$^{Ex6m\ -/+}$ mice showed no significant changes in glomerular matrix deposition.

**G.** C-terminal truncating TRIM8 variants were generated in FVB/N mice (*Trim8*$^{Ex6m}$). Podocyte injury was induced with sheep nephrotoxic serum (NTS). Albuminuria was assessed to follow injury and recovery. Female heterozygous and homozygous mice did not exhibit significantly different albuminuria at any timepoints in a three-week time course after injury.

**H.** In the NTS study in (G), total albuminuria defined as the area-under-curve was not significantly different between *Trim8*$^{Ex6m}$ female mice and littermate wildtype controls. *Trim8*$^{Ex6m}$.

**I.** In the NTS study in (E), peak albuminuria was not significantly different between *Trim8*$^{Ex6m}$ female mice and littermate wildtype controls.

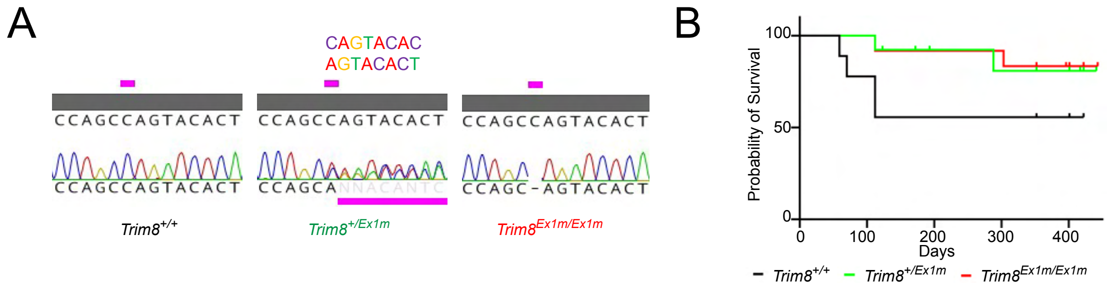


C

| | WT | HET | HOM |
|---|---|---|---|
| Combined actual | 15 | 31 | 19 |
| Expected | 16.25 | 32.5 | 16.25 |

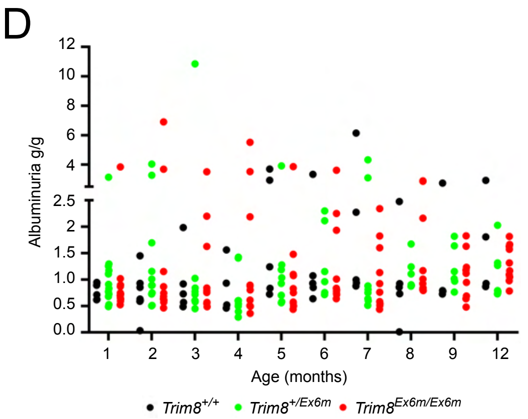


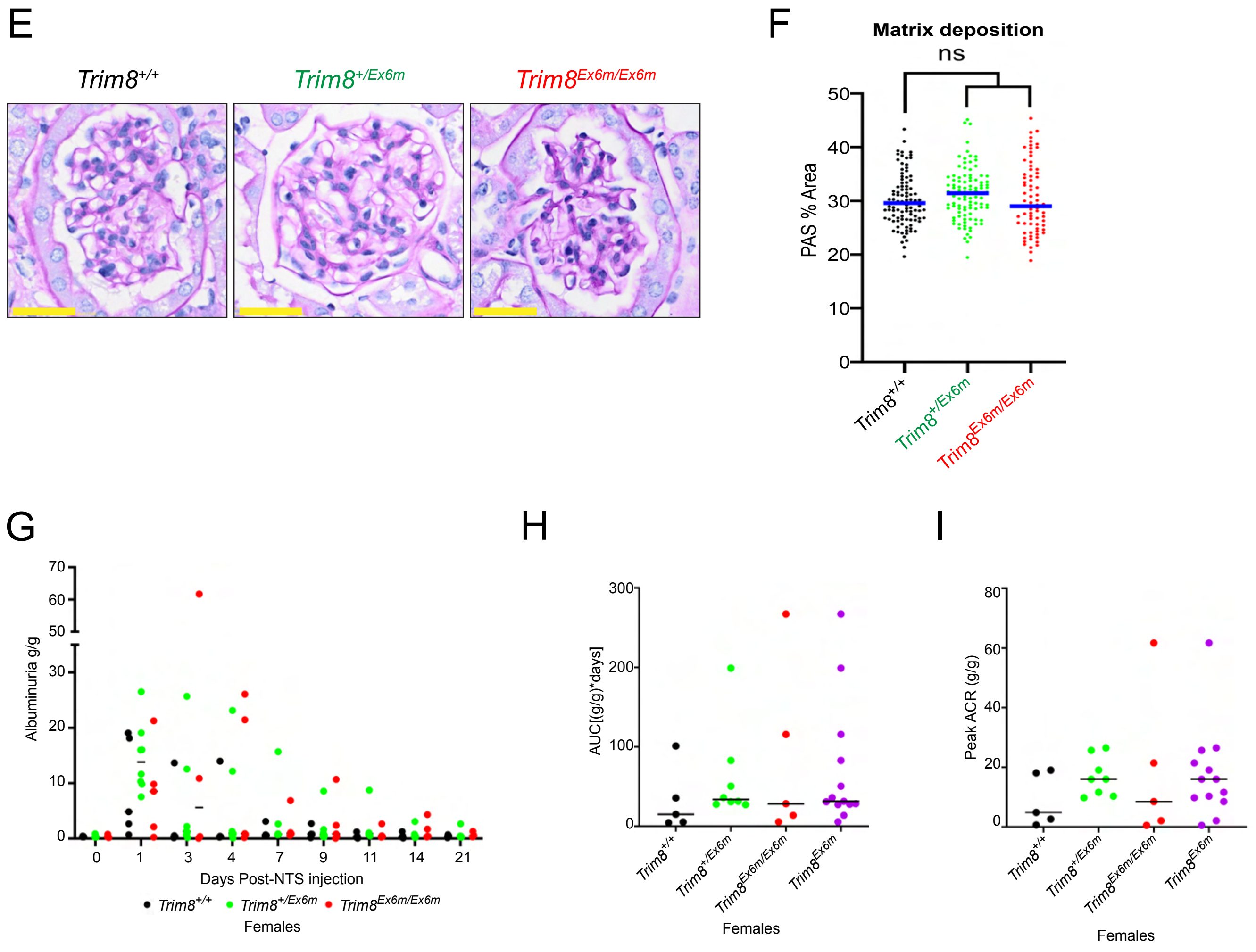

**All supplementary tables, files, movies and documents can be found online at:**

https://urldefense.com/v3/__https://cloud.mpi-cbg.de/index.php/s/jNcp3pzD6t6c04L__;!!NZvER7FxgEiBAiR_!uoqZoH5rzbOhrdm15Ly-3DiEkStrp3cqXv-a0CR9xbuMqVTqFZwN1kYW4dEzkkOGkILq9UtKi0VtKk_rimXAEB2fKmEFwubjfwtdpkB9$

### Supplementary Tables

Supplementary Table ST1. Domain structures and disordered content (PONDR-VSL2, FuzDrop, PICNIC) are displayed for each TRIM family protein with corresponding gene symbol and Uniprot symbol given.

Supplementary Table ST2. Human genetic diseases associated with TRIM family genes based on Online Mendelian Inheritance in Man (OMIM) curation or literature sources is shown.

Supplementary Table ST3. Summary cell biological phenotypes are shown for TRIM protein screen including predominant mesoscale localization and dynamics after photo-bleaching (in relation to disordered content).

### Supplementary Files

Supplementary File SF1. Primary amino acid sequences are listed in FASTA format for all human TRIM proteins based on the Uniprot ID found in **Suppl Table ST1**.

Supplementary File SF2-SF72. Alpha-fold derived structures (.pdb format) are provided for all human TRIM protein that informed domain mapping as listed in **Suppl Table ST1** and are basis for structures shown in main figures.

### Supplementary Movies

Supplementary Movies SM1-SM51 are shown for 48 TRIM proteins evaluated for fluorescence recovery after photobleaching to define dynamics of these mesoscale entities.

### Supplementary Documents

Supplementary Document SD1 contains the Key Materials and Equipment used in this study.
